# Structural polymorphism and population-variable coding capacity of HERV-K(HML-2) in human pangenomes

**DOI:** 10.64898/2026.09.22.753638

**Authors:** Daniel A. Murimi-Worstell, Michael Freeman, Anna Minkina, Mitchell R. Vollger, Andrew B. Stergachis, John M. Coffin

## Abstract

Approximately 8% of the human genome is derived from ancient retroviral infections. The most recently integrated of these endogenous retroviruses is the HERV-K(HML-2) clade, whose expression has been associated with cancer, amyotrophic lateral sclerosis, and embryogenesis. Studies of HERV expression, particularly HML-2, have relied predominantly on short-read sequencing. However, the high similarity among HML-2 proviruses prevents many short reads from being assigned uniquely to individual loci. We therefore compared haplotype-resolved long-read genome assemblies from 292 donors to resolve variation in proviral structure and coding capacity. Several loci previously thought to be fixed were structurally polymorphic. Tandem arrays occurred at 13 loci and contained up to six proviral copies in a single array. At 8q11.23, we identified a previously undescribed full-length provirus in one haplotype. All 583 other haplotypes carried a solo-LTR. We found that standard reference genomes failed to represent the coding capacity retained in many individuals, whose proviruses contained intact open reading frames despite disruptive mutations in the reference sequences. Short-read genotypes left 32.5% of the tested donor–variant pairs unresolved at sites associated with viral reading frames. These findings show why HML-2 expression must be interpreted in the context of the structural and coding alleles each individual carries.

**Graphical abstract:** Alt text: A schematic summarizes haplotype-resolved HML-2 structural variation and differences in predicted coding capacity across individuals.

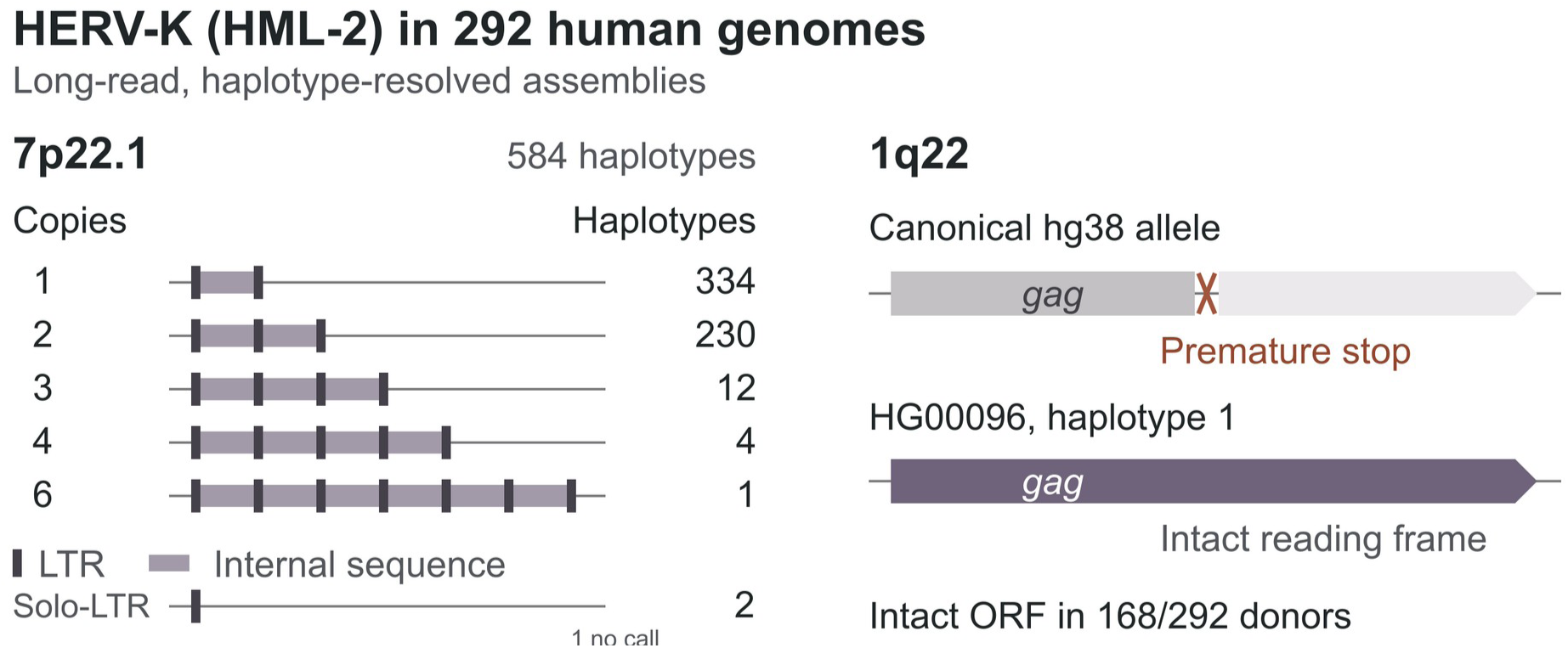

## Introduction

Human endogenous retroviruses (HERVs) represent the remnants of ancient germline retroviral infections in human ancestors. These remnants are usually lost via negative selection or genetic drift, but occasionally, by positive selection or drift, they become fixed in the human genome. Over millions of years, HERVs and related long terminal repeat (LTR) elements have grown to comprise approximately 8% of the human genome (1). Since their initial integrations, most HERVs have undergone extensive mutation and are usually epigenetically silenced. However, younger HERVs have had less time to accumulate mutations, and many can still encode viral proteins. Several elements of the most recently integrated clade of HERVs, HERV-K(HML-2) (hereafter, HML-2), retain open reading frames (ORFs) for viral proteins, which has driven particular interest in HERV research regarding this group (2,3), although no naturally occurring provirus has been shown to encode an infectious virus. Altered expression of transcripts and proteins derived from HML-2 elements has been identified in myriad physiological and pathological tissue states such as embryogenesis, aging, amyotrophic lateral sclerosis (ALS) and cancer (3–8).

Interpreting these expression patterns requires knowing which viral genes each expressed HML-2 element retains and which of these genes maintain coding capacity. The integrated viral genome, or provirus, contains gag, pro, pol, and env genes. The pol gene encodes reverse transcriptase and integrase. Two long terminal repeats (LTRs) flank these coding regions and contain regulatory sequences. Recombination between the LTRs can remove the internal viral genes entirely, leaving a solo-LTR at the same genomic site. Short stretches of host DNA duplicated during integration remain on either side of the provirus or solo-LTR. These target-site duplications (TSDs) mark the insertion boundaries (2,3). During translation, canonical frameshift events are required to generate gag-pro and gag-pro-pol polyproteins for protein-level expression of pro and pol.

Most HML-2 proviruses have lost some viral coding capacity through accumulated mutations. Type-I HML-2 proviruses carry a shared 292-base-pair deletion at the pol-env boundary that distinguishes them from Type II. Type-II proviruses retain this region and can produce Rec, an accessory protein that promotes viral RNA export from the nucleus. The deletion in Type I removes the splice donor used to produce Rec, and alternative splicing instead allows expression of Np9 (2,3).

Determining which of these structures and coding sequences an individual carries is complicated by the extensive similarity among HML-2 elements. Short reads often match multiple loci and cannot be assigned uniquely to their genomic source. This has driven the development of sophisticated statistical methods and tools to assign multimapping RNA-seq reads to individual loci (9). Although these methods have facilitated transcriptomic quantification at individual HERV loci, they do not reconstruct the full structural alleles present in each individual. Modern techniques based on short reads aligned to reference genomes are especially limited in their capacity to characterize the most recently integrated, and therefore most similar, HML-2 elements.

Recent analyses of HML-2 highlight this limitation. Researchers have identified HML-2 transcripts that lack inactivating mutations present in reference genomes (7), as well as proviruses absent from standard human reference assemblies (10,11). These findings suggest that relying on short-read sequencing data and standard human reference genomes masks the genetic diversity and coding capacity of HML-2, and therefore, the possible mechanisms by which HML-2 may affect health and disease. To address this limitation, we systematically explored HML-2 genetic variation in long-read genome assemblies from the Human Pangenome Reference Consortium (HPRC, releases 1 and 2) and the Human Genome Structural Variation Consortium (HGSVC3) (12–14). These datasets span 292 donors. Across these genomes, we find that the HML-2 clade is a structurally polymorphic and individually variable component of the human genome. HML-2 element structure, copy number, coding potential, and integration itself differ from person to person. Our results suggest that to elucidate HML-2’s biological roles in future studies, researchers must consider individual elements on a genome-by-genome basis rather than assuming concordance with standard genomic references.

## Materials and Methods

### Genomic Sequence Extraction and Pangenome Graph Processing

We established genomic coordinates for HML-2 loci from the LTR5Hs, LTR5A, and LTR5B subfamilies using the linear T2T-CHM13v2.0, hg38, and hg19 reference genomes (15). We obtained graph-based pangenomes from HGSVC3 and assemblies from HPRC releases 1 and 2 from the Human Pangenome Reference Consortium and Human Genome Structural Variation Consortium data portals (see Data Availability) (12–14). For the graph-based pangenome assemblies, we extracted local subgraphs encompassing 15 kb of flanking sequence around each target locus with the odgi toolkit (16). We used odgi untangle to project haplotype paths onto the reference paths. Graph-processing parameters are provided in Supplemental Methods.

For linear FASTA assemblies, we mapped contigs to the references with minimap2 using the asm20 preset (17), extended locus coordinates by 15 kb on each side, verified alignment orientation with pysam, and extracted the regions with samtools faidx (18).

The dataset contained 292 donor genomes and four reference assemblies. The HML-2 catalog comprised 99 named loci and four groups of copies lacking a unique genomic assignment (Table S1). Donors contributed 584 copies of each autosome. The 288 donors with known sex contributed 432 X and 144 Y copies. Structural fractions used chromosome copies with a locus-level call, reported alongside the total chromosome count. Matched analyses used only individuals and loci represented in both datasets (Table 1).

Published sequence classifications and direct sequence matches identify 8p22 and 17p13.1 as HML-11 rather than HML-2 (2). We retained those records for reference but excluded them from the HML-2 analyses (Supplemental Methods).

**Table 1.** Cohorts and denominators used in the main analyses.

| Analysis set | Denominator | Use |
| --- | --- | --- |
| Donor assemblies | 292 donors, 584 autosomal copies, 432 X and 144 Y copies | Structural calls, coding distributions, and 7p22.1 carrier counts |
| Reference assemblies | 4 reference genomes | Coordinate and reference-allele comparisons |
| Published short-read resources | 282 shared donors at 83 loci; 564 mapped ORF-associated variants at 36 loci. Callset-specific overlap for structural variants | Matched VCF/assembly comparisons (Figure S1) |

### HML-2 structural classification and motif analysis

We aligned sequences to the KCON (19) reconstructed intact HML-2 sequence to define proviruses, solo-LTRs, fragments, and tandem arrays. We annotated array members individually. We identified a provirus as Type I by directly detecting the canonical 292-bp pol-env deletion. Supplemental Methods describe additional alignment and boundary criteria.

We identified candidate TSDs by extracting sequences directly adjacent to the LTRs. We retained equal-length 4–6-bp pairs with up to two mismatches. We compared candidate lengths before interpreting differences (Supplemental Methods).

### Read depth and array support

We obtained Oxford Nanopore whole-genome reads for the matched donors from the sample-indexed raw-read files in the public HPRC sequence archive (Data Availability and Table S2 supporting data). Array support required reads spanning both outer boundaries at ≥80% alignment identity. To assess suspected extra copies at separate assembled positions, we measured read depth across each provirus and its adjacent host sequence. For each donor, we used the median positive flank-depth measurements as the diploid baseline. We divided each candidate’s local-flank depth by this baseline. We excluded candidates with ratios ≤0.5 from population counts unless independent sequence and flank evidence established duplication of the provirus together with its surrounding host DNA (segmental duplication). We also excluded copies crossing assembly gaps and counted repeated labels for the same assembled interval once. Supplemental Methods describe the depth model and the screen for mixed bases within assembled haplotypes.

### Validation of structural calls

We evaluated candidate additional copies using spanning reads, local and genome-wide read-depth comparisons, assembly continuity, and host-flank alignments. We excluded copies unsupported by these checks from population summaries. We also identified records representing the same assembled interval under different locus labels and counted each interval once. We used NucFreq to screen for collapsed sequence copies. Supplemental Methods and Table S2 provide thresholds, locus-assignment rules, and record-level decisions.

### New HML-2 detection

We screened each assembly with RepeatMasker to identify HML-2 proviral sequence absent from the corresponding T2T reference interval (RepeatMasker Open-4.0, https://www.repeatmasker.org). This search could recover a new occupied site or an internal proviral sequence at a known solo-LTR site. We aligned candidates to KCON for ORF annotation. We compared their locations with known loci using bedtools intersect (20) and excluded sequences that matched a known locus without a structural change.

### Open reading frame (ORF) integrity analysis

At each locus, we aligned haplotype sequences with the KCON reference using MAFFT (21) and projected gag, pro, pol, env, rec, and np9 annotations onto the alignment. We scored sequence coverage and ORF integrity with Biopython (22).

We recorded additional sequence frameshifts, premature stops, missing start codons, and deletions. We scored each gene in its own annotated reading frame. We did not count the canonical programmed frameshifts between gag, pro, and pol as disruptive sequence changes. We defined coverage below 30% as a major deletion. Intact annotations had no additional frameshift or premature stop and met the applicable start-codon and length criteria. To retain candidates that might encode altered proteins, we also used a combined ORF screen. This included intact annotations and sequences lacking a premature stop despite a first additional frameshift at or beyond 40% of the reference protein, or a non-triplet length leaving a terminal incomplete codon. These additional candidates do not imply retention of the usual protein function. Gag–Pro and Gag–Pro–Pol assignments required the corresponding annotations on one provirus. We also mapped substitutions and frameshift positions and identified each copy’s longest single-frame ATG-starting ORF of at least 200 amino acids. We classified translated sequences shorter than 60% of the reference protein as fragments and excluded them from the combined screen. Type-I Env denotes the theoretical N-terminally truncated product of the remaining env reading frame after Δ292. Translation of this product has not been demonstrated. Supplemental Methods give the remaining scoring rules.

### Matched long-read and short-read comparison

We compared long-read alleles with native, unphased genotypes from the high-coverage 1000 Genomes short-read callset (23). The shared catalog panel contained 282 donors and 83 loci. We identified single-nucleotide changes that gained or removed a premature stop codon in the canonical Gag, Pro, Pol, or Env frame, and indels shorter than 50 bp whose length change was not divisible by three. The latter were frame-changing variants relative to hg38; they could disrupt or restore a reading frame. We annotated each gene separately, so we did not count canonical Gag–Pro–Pol frameshifts as sequence defects. We compared each normalized allele once per donor and locus, irrespective of haplotype or copy number. Supported short-read calls required a complete genotype, depth of at least 10, genotype quality of at least 20, and passing site and sample filters. Missing records, missing or partial genotypes, inadequate quality, and ambiguous variant representations remained unresolved. We reported alternate-allele support, explicit reference calls, other alternate calls, and unresolved observations separately (Table 2, Table S3). These variant calls do not establish complete ORFs or link variants across a provirus.

We also examined four structural-variant callsets for 1000 Genomes Project samples. These were the phased high-coverage panel, the Illumina ensemble structural-variant callsets from freezes V1 and V3, and the HGSVC2 PanGenie callset (23–25). Figure S1 compares the Illumina ensemble calls with matched long-read assemblies. We averaged disagreement across records within each donor–locus combination and weighted each combination equally.

### Phylogenetic analyses

We aligned sequences with MAFFT (21) and built neighbor-joining trees (26) separately for the LTR, gag, pro, pol, and env regions, collapsing identical sequences within each locus. For each sequence pair, nucleotide divergence was defined as the number of nucleotide differences divided by the number of aligned positions with A, C, G, or T in both sequences. Gaps and ambiguous bases were excluded. For each tandem array, we counted nucleotide substitutions and contiguous alignment gaps between every pair of internal proviral sequences. We recorded the base at each variable nucleotide site in the copies’ order along the array. For arrays of at least three copies, we identified the pair or pairs with the lowest nucleotide divergence to assess whether expansion involved repeated duplication of particular copies (Supplemental Methods).

We assessed unrooted LTR and Pol trees with 1,000 site-bootstrap replicates and recorded nearest-neighbor frequencies within the 64-locus comparison shared by both trees. We assigned each tied neighbor an equal fraction of that replicate’s count. We used midpoint positioning only for display. We assigned telomeric copies to chromosome short arms using linked host sequence and kept ambiguous copies in separate groups.

We connected two loci in the sequence-sharing network if they shared at least one identical gag, pro, pol, or env sequence. For each connected pair of loci, we compared every sampled copy at one locus with every sampled copy at the other, separately for each available gene. We averaged the nucleotide-divergence values from these comparisons to label each connection. At Xq28a and Xq28b, this comparison used only env, the gene represented in the sequence-sharing dataset for those loci. We aligned their env sequences directly with MAFFT.

We additionally compared paired LTRs and flanking host sequences in the telomeric Type-II family and quantified nucleotide sharing across assigned population copies (Figure S2A–C).

### Chromatin accessibility analysis

We analyzed bonFIRE consensus peaks and Fiber-seq inferred regulatory element (FIRE) calls from lymphoblastoid cell lines representing a matched subset of 39 genome assembly donors (13,27,28). For each haplotype, we calculated accessibility as pooled FIRE coverage divided by pooled total read coverage across the assayed regulatory peaks. Supplemental Methods specify sequence-selection and coverage criteria.

### Type-I deletion analyses

To compare sequence conservation around Δ292 with the rest of the provirus, we used a fixed window containing 501 bp immediately upstream and 500 bp immediately downstream of the deletion in KCON. We concatenated the two flanks and excluded the deleted interval. The 1,001-bp window provided a comparison on a scale similar to six 1-kb windows elsewhere in the provirus. We selected one population-derived representative per locus and compared nucleotide divergence and nearest neighbors across these windows. The panel contained 15 Type-I and 30 Type-II representatives. We analyzed Type-II both as a pooled group and after restriction to its 15 LTR5Hs representatives. Direct calls of the Δ292 interval distinguished Type-I from Type-II sequences. Supplemental Methods provide the window coordinates, coverage thresholds, and representative-selection procedure.

To test whether neutral conversion after integration could explain the absence of Type-I/II polymorphism within loci, we modeled recurrent conversion to Type I and genetic drift at loci that were initially Type II. We calculated the probability of sampling only Type-I alleles at each observed Type-I locus, conditional on detecting Type I. We assumed independent loci and multiplied these probabilities across the observed Type-I loci. The central scenario used an effective population size of Nₑ=10,000 and a conversion rate of 3.2 × 10⁻⁶ per generation. The sensitivity grid varied conversion rate, population size, and locus-age bounds across 54 scenarios, including doubled older age bounds. A permissive scenario used Nₑ=5,000 and the highest tested rate and granted complete Type-I fixation to every older or unaged locus.

Supplemental Methods specify the diffusion equations and the full parameter grid. Table S13 reports the scenarios and numerical validation. We separately compared the numbers of ancestral proviral integrations carrying Δ292 or six other deletions of at least 50 bp detected in orangutan, siamang, or macaque HML-2 sequences. We treated sequences missing the same viral interval as one deletion class. We compared the host DNA on both sides of each provirus to identify insertions inherited from a shared ancestral integration and count them once across species. Each deletion class received a model weight for its cumulative contribution to those integrations. One model allowed those weights to differ among classes. The second also allowed a multiplier for Δ292 to represent a deletion-specific propagation advantage. These weights summarize contributions over the sampled history; they do not identify individual source loci, their lifetimes, or a transmission genealogy. We integrated each model’s likelihood over its specified priors and compared the resulting marginal likelihoods. Supplemental Methods describe the orthology-based counting, deletion-class contribution priors, detection-odds ranges, and likelihood integration (Table S11).

### Functional association analyses

We screened HML-2 structural states and ORF-annotation burden for associations with gene expression, growth, Epstein–Barr virus (EBV) DNA abundance, and anti-CD20 response in lymphoblastoid cell lines (LCLs). We used EBV DNA data from Mandage et al. (29) and Houldcroft et al. (30), growth data from Im et al. (31), expression data from MAGE (32) and GEUVADIS (33), and anti-CD20 viability data from Small et al. (34). We used covariate-adjusted regression and Benjamini–Hochberg correction (35). Supplemental Methods detail cohort selection, exposure definitions, covariates, testing families, and sensitivity analyses.

### Sequence variation at 8q11.23

At 8q11.23, we compared the two LTRs of the full provirus and measured sequence variation among the solo-LTR alleles. To compare solo-LTR diversity, we first aligned sequences to the 968-bp HML-2 LTR reference using minimap2. We calculated nucleotide diversity as the sum of nucleotide differences across all sequence pairs divided by the total number of comparable A/C/G/T positions across those pairs. Each donor haplotype contributed once, so identical sequences retained their observed counts. We compared solo-LTR diversity across loci with at least 20 retained sequences and examined its relationship to sequence-divergence expectations under two published pairwise LTR rates (2) (Supplemental Methods, Table S4). To place the sequence variation in the history of the occupied site, we examined reads spanning the host–LTR junctions in published Neanderthal and Denisovan genomes (36–39).

## Results

### HML-2 copy number and structure vary between people

Across 292 donor genomes, we identified population-variable HML-2 tandem arrays at 13 loci. Such arrays had previously only been described at 6q14.1 and 7p22.1 (40,41). Their consecutive proviral copies share intervening LTRs, and long-read assemblies resolved each copy’s sequence. Reads spanning the array distinguish structures that collapse into a single apparent element when short reads are mapped against a single-copy reference (Figure 1A). Across the 24 most structurally variable loci, we observed proviruses, solo-LTRs, fragments, and multi-copy states, as well as noncarrier calls (Figure 1B, Tables S1 and S5).

We checked candidate additional copies against spanning reads, read depth, assembly continuity, and host-flank alignments to distinguish true duplications from assembly errors and misassigned loci. These checks excluded 35 unsupported additional-copy records and counted 78 repeated observations of assembled intervals only once. We retained nine independently supported segmental duplications (Figure S3, Table S2). We retained copies without a unique supported host-flank assignment in separate groups (Table S5).

**Figure 1:**
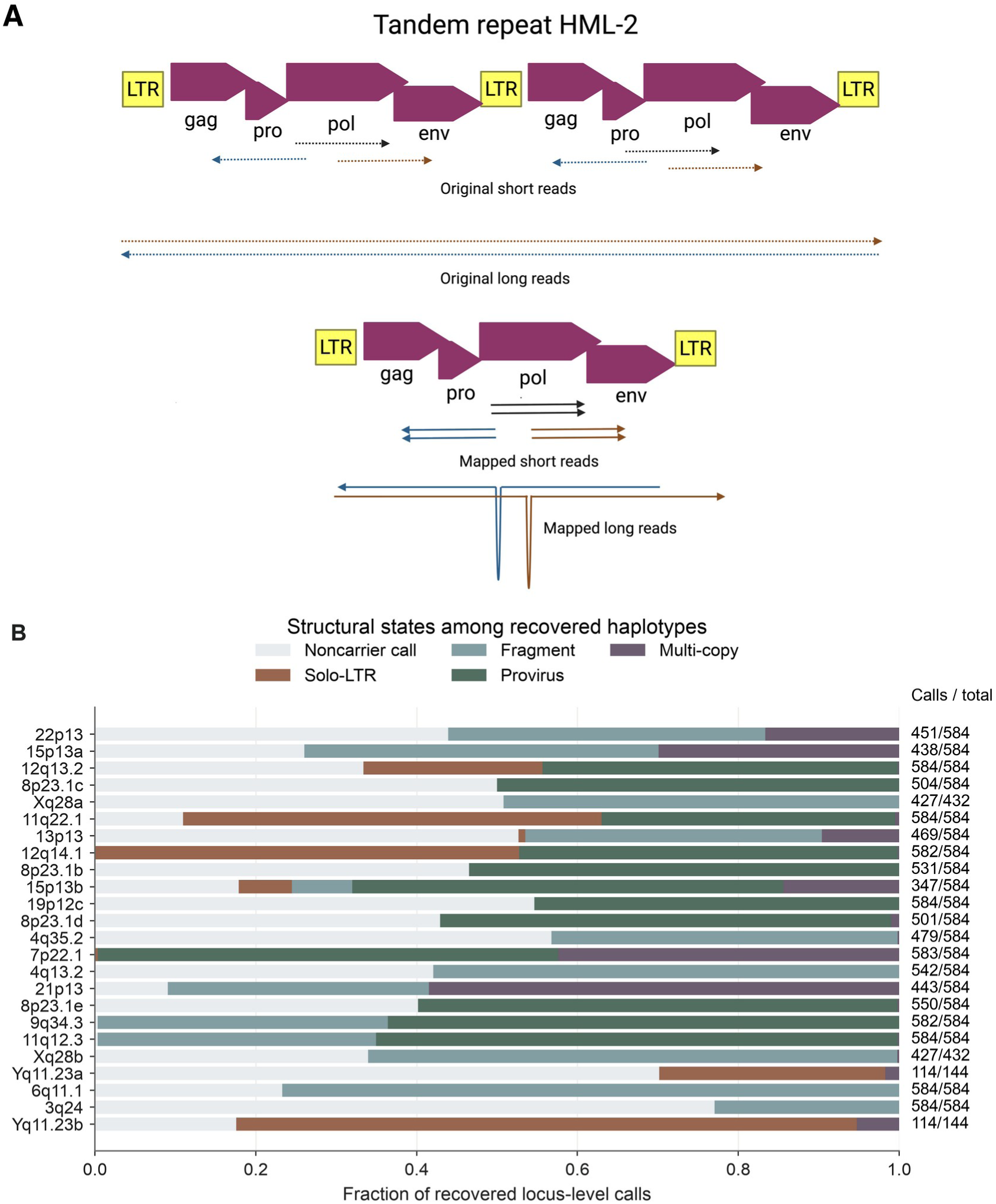
Structural variation in the HML-2 pangenome. **(A)** Recovery of tandem-array structure with spanning long reads. **(B)** Structural-state fractions at the 24 most variable loci, ordered from top to bottom by decreasing fraction of called haplotypes outside the most common structural state. Labels give calls/total chromosome copies. Totals are 584 for autosomes, 432 for X, and 144 for Y. Per-locus counts and counting rules are provided in Table S1 and the Supplemental Methods. Panel A: Created in BioRender. Murimi-Worstell, D. (2026) https://BioRender.com/2iwl6rp. Alt text: Long reads distinguish tandem copies hidden by short-read mapping. A stacked bar chart compares provirus, solo-LTR, fragment, multi-copy, and noncarrier states at 24 variable loci.

### Tandem arrays carry distinct viral coding sequences

We next asked how many proviral copies each tandem array contained and whether those copies differed in sequence or coding capacity. Across the 13 loci, arrays contained up to six copies at 7p22.1 and four at 1p31.1b (Figure 2A, Table S6). Multi-copy states occurred in 247 of 584 sampled haplotypes at 7p22.1 (42.3%, or 42.4% of the 583 called haplotypes), 13 at 12q24.33 (2.2%), nine at 1p31.1b (1.5%), and one to four at each of the other ten loci (0.2–0.7%, Figure 2B).

Individual elements within arrays differed in amino-acid substitutions and ORF annotations. At 7p22.1, these annotations differed among copies in 181 of 247 arrays for Gag, 180 for Pro and 172 for Pol (Figure 2C–E). Most Gag differences altered the sequence without changing the ORF classification. Env varied less. It differed in 15 of 247 arrays at 7p22.1, one of nine at 1p31.1b, and the single array at 14q11.2 (Figure 2F).

ORF retention also differed by position within arrays. Under the combined screen defined in Methods, 72/247 first-position and 122/247 second-position copies at 7p22.1 passed for Gag, compared with 77/247 and 244/247 for Pro (Figure S4A). Among array-bearing haplotypes, 230/247 arrays at 7p22.1 contained two proviruses, whereas seven of nine at 1p31.1b contained three or four (Figure S4B, Table S7).

We compared nucleotide differences among array members to ask whether their expansion involved repeated duplication of particular copies (Figure S5, Table S8).

Copies 2 and 3 were the closest pair in all twelve three-copy 7p22.1 arrays. In the six-copy array, copies 2–6 were identical internally, and each differed from copy 1 at ten nucleotide sites (Figure S5C). This pattern is consistent with expansion of a downstream copy lineage after divergence from copy 1. At 1p31.1b, copies differed by at most one substitution, and several arrays contained identical internal sequences (Figure S5D). Supplemental Results and Table S8 provide detailed sequence comparisons and conditional divergence estimates.

**Figure 2:**
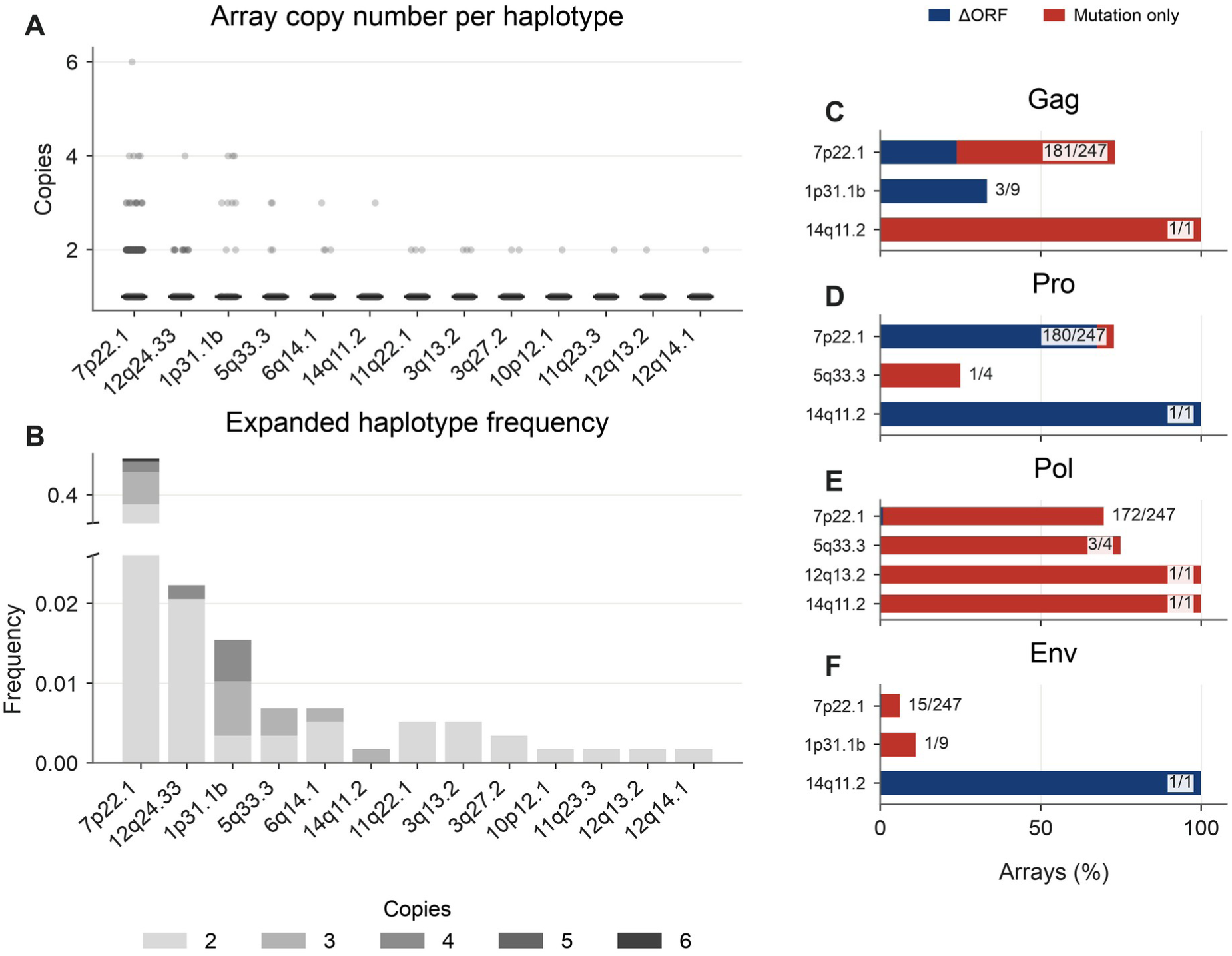
Copy number and coding variation in HML-2 arrays. **(A)** Copy-count distributions at 13 loci. The denominator is 584 haplotypes except at 7p22.1, where 583 were called. **(B)** Multi-copy frequencies among all 584 haplotypes. **(C–F)** Arrays containing more than one Gag, Pro, Pol, or Env sequence profile. Bars distinguish changes in ORF state from sequence differences that preserve that state. Labels indicate arrays containing differing profiles/arrays tested. At Type-I loci, Env refers to the theoretical N-terminally truncated product. Alt text: Copy-count distributions and array frequencies show extensive variation at 7p22.1 and rarer arrays elsewhere. Within-array differences are more frequent for Gag, Pro, and Pol than Env.

### Short-read genotypes leave many ORF-associated variants unresolved

To determine whether existing short-read calls more generally captured coding differences observed in the assemblies, we tested 564 ORF-associated small-variant alleles at 36 loci in 282 donors. These comprised 20,500 donor–variant pairs. Short-read genotypes supported the assembly-derived allele in 13,717 pairs (66.9%). Another 118 had a supported reference genotype, and 10 supported a different alternate allele. The remaining 6,655 pairs (32.5%) were unresolved. Among the 13,845 pairs that passed the genotype and representation checks, 99.1% agreed with the long-read allele (Table 2).

The distinction was particularly important at 1q22. Long-read sequences identified 272 donors carrying the single-nucleotide variant that removes a premature Gag stop codon in hg38. Short reads supported this allele in 37 donors and supported the reference genotype in seven. The other 228 calls were unresolved because of low depth, low genotype quality, or a missing genotype (Table S3). The separate Illumina ensemble structural-variant comparison showed disagreement about allele presence (Figure S1A) and structural state. Structural-state disagreement was highest at 11q12.1 in 194 of 223 comparisons (87.0%, Figure S1B).

We then asked how these gaps affected evidence for ORF intactness. Among 2,330 long-read-intact gene copies carrying at least one targeted variant, 2,166 (93.0%) had an unresolved call at one or more of those sites. This subset spanned nine loci. Another 10,934 intact gene copies had no qualifying target and were outside this denominator (Table S3). Thus, even known ORF-associated differences often lacked sufficient short-read support to confirm the intact sequence. Support at every targeted site would still leave unreported bases and unassessed linkage across the ORF.

**Table 2.** Short-read genotype support for ORF-associated variants observed in long-read assemblies.

| Coding region | Donor-variant pairs | Alternate supported | Reference supported | Other alternate | Unresolved |
| --- | --- | --- | --- | --- | --- |
| Gag | 5,516 | 3,617 (65.6%) | 48 | 8 | 1,843 |
| Pro | 2,719 | 1,146 (42.1%) | 43 | 0 | 1,530 |
| Pol | 7,774 | 4,649 (59.8%) | 26 | 2 | 3,097 |
| Type-II Env | 5,110 | 4,355 (85.2%) | 5 | 0 | 750 |
| Type-I Env* | 518 | 484 (93.4%) | 0 | 0 | 34 |
| Total | 20,500 | 13,717 (66.9%) | 118 | 10 | 6,655 |

Totals count unique donor-locus-variant pairs because a variant can affect overlapping genes. \*Type-I Env denotes a theoretical N-terminally truncated product whose translation has not been demonstrated.

### HML-2 loci share identical gene sequences but differ in regional ancestry

We examined both insertion boundaries and internal viral sequences for evidence of how recombination and duplication had shaped HML-2 loci. TSDs flanking an insertion originate from the same pre-integration sequence. Differences between them can therefore identify changes at insertion boundaries. We compared the sequences immediately outside the left and right boundaries of 12,095 retained element records at 26 loci. Both boundaries were available for each record. In 12,075 records, the two sequences matched at one or more candidate TSD lengths from 4 to 6 bp. The remaining 20 records at five loci differed at every tested length. These counts included repeated observations of inherited alleles. Supplemental Results and Figure S6 report the sequences and boundary checks.

Within proviruses, recombination can join sequences from different ancestral viruses, so an element’s LTR and coding regions may have different closest relatives. We compared trees for these regions to look for such ancestry changes. The most frequent 19p12c and 10q24.2 sequences had different nearest neighbors in the LTR and Pol trees, although bootstrap support for those specific relationships was limited (Figure 3A). Supplemental Results and Figures S7– S8 report the regional comparisons and their support.

Some copies at distinct loci shared an identical complete viral gene sequence (Figure 3B). This established a connection between loci even when other copies differed. The divergence shown for each connection averages all eligible between-locus copy pairs, not just the identical sequences. All six pairs among 8p23.1b–e shared complete gene-region sequences; their mean between-locus divergence ranged from 0.49% to 1.03%. Figure 3B and Table S9 give the remaining comparisons. Host-flank alignments provided additional evidence of duplication at 4q35.2 and the telomeric Type-II loci (Figure 3C). The most frequent 4q35.2 Pol sequence differed from a 21p13 sequence at only 4 of 2,741 jointly called bases. Reference alignments linked the provirus and at least 5 kb of host sequence on each side, supporting descent from a shared duplicated segment.

### Telomeric proviruses trace an older insertion through duplicated chromosome segments

The shared viral and host sequences at the telomeric Type-II loci suggested duplication of an ancestral insertion. To reconstruct this history, we compared paired LTRs and host flanks at 4q35.2, 15p13a, 21p13 and 22p13. The two LTRs within each reference provirus differed at 52– 55 of 993–996 jointly called positions (5.22–5.54%). The tree separated the 5′ LTRs from the 3′ LTRs across all four loci (100% bootstrap support, Figure S2A). This indicates that their divergence preceded duplication. Applying the published pairwise LTR clock of 0.24–0.45% divergence per million years (2) gives approximately 12–23 million years for the ancestral insertion.

The reference provirus and upstream host sequence placed 21p13 and 22p13 closest together. They differed at 7 of 7,275 proviral positions and 5 of 4,997 upstream positions. Twelve proviral sites and 25 upstream sites supported this grouping against 4q35.2 and 15p13a, recovered in all 2,000 bootstrap replicates for each region (Figure S2B, Table S9). The downstream flank instead grouped 15p13a and 21p13. These loci differed at 7 of 4,992 downstream positions, compared with 20 differences between 21p13 and 22p13. Three informative sites supported the alternative grouping, recovered in 93.8% of bootstrap replicates (Figure S2C). This change across the duplicated segment is consistent with subsequent sequence exchange. Exchange among the short arms of heterologous acrocentric chromosomes has also been observed in human pangenome assemblies (42).

Population sampling revealed further sharing among these loci. We compared 426 assigned copies from 197 donors at the four loci and 4p16.3a. Across 6,652 well-covered positions, average divergence between the telomeric loci 15p13a, 21p13 and 22p13 was 0.253–0.284%, similar to within-locus diversity of 0.229–0.273%. None of these positions was fixed for different nucleotides between any pair of these three loci in the sampled copies. Seventeen nucleotide profiles were shared between loci across the five-locus family at the 4,797 positions called in every copy (Table S9). The close similarity of these proviruses therefore reflects redistribution and exchange of an older insertion rather than recent viral integration alone. Their shared nucleotide profiles also make it impossible to infer chromosomal position from some viral sequences alone. Host flanks are needed to distinguish those copies.

**Figure 3:**
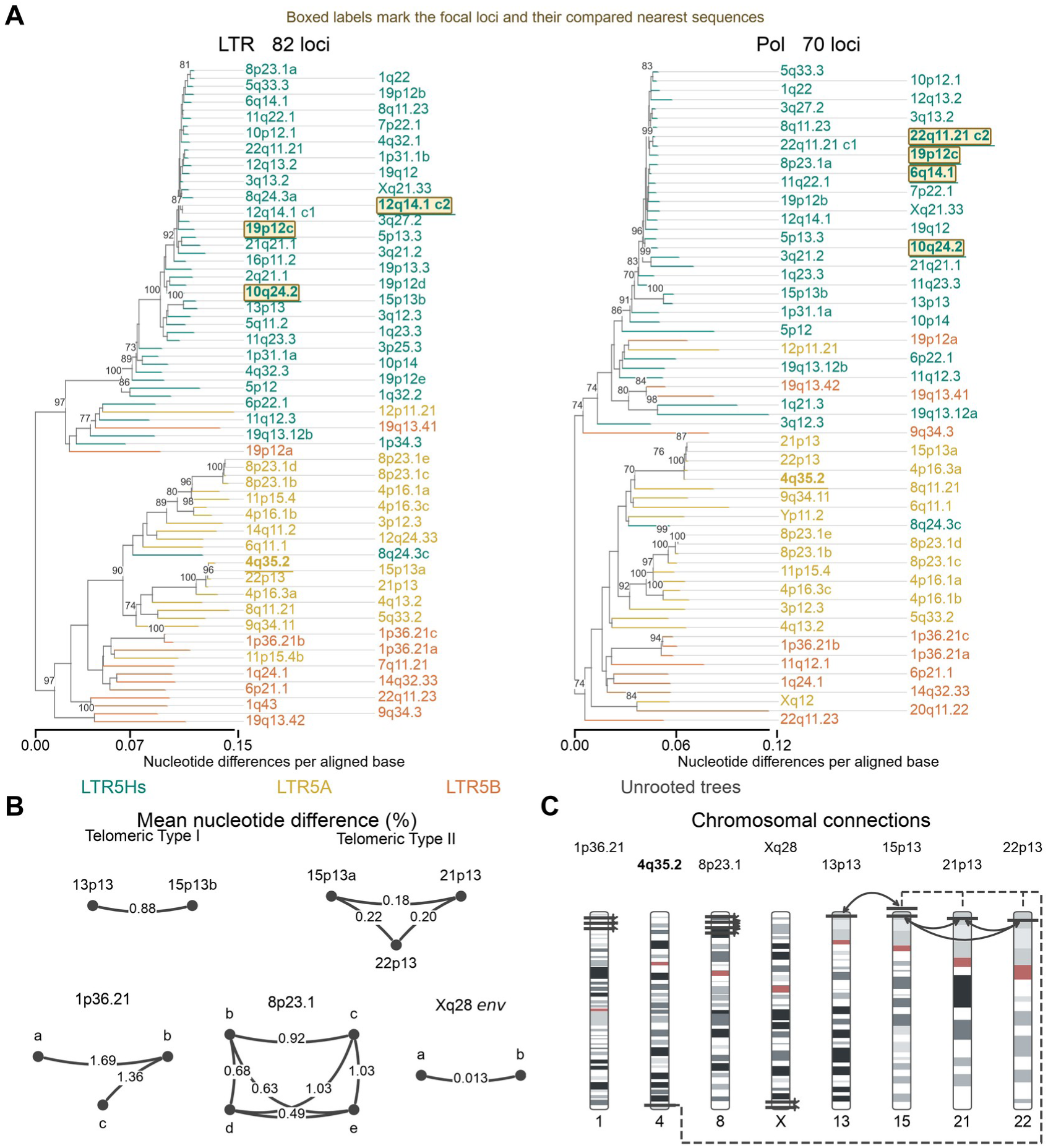
Regional sequence relationships among HML-2 loci. **(A)** Unrooted neighbor-joining trees. Branch lengths show substitutions per aligned base. Numbers show bootstrap support ≥70%. Colors identify LTR subfamilies. Boxes mark loci discussed in the text. The 4q35.2 labels are underlined. c1 and c2 denote frequency-ranked clusters. **(B)** Loci sharing an identical gene-region sequence. Labels give average pairwise nucleotide differences (APD, %) among sampled copies. The Xq28 comparison uses env only. **(C)** Chromosomal locations of connected loci. Solid lines denote exact sequence sharing. The dashed line denotes the viral and host-flank relationship between 4q35.2 and telomeric Type-II loci. Chromosomes are scaled separately. Alt text: LTR and Pol trees place some loci differently. A network shows sequence-sharing links and their mean nucleotide differences, and a chromosome map locates the related groups.

### Individuals carry different combinations of HML-2 coding sequences

We annotated each resolved copy to determine which viral genes were retained together. Under the combined ORF screen, we found that Np9-only combinations were most common in Type I, and Rec-only combinations in Type-II proviruses. Other combinations included Gag, Pro, and Pol on the same provirus (Figure 4A–B). Table S10 gives the corresponding frequencies at each locus.

Across 292 donors, mean copy counts passing the combined ORF screen were 17.1 for Gag, 9.1 for Gag–Pro, 4.1 for Gag–Pro–Pol, 8.0 for Type-II Env and theoretical Type-I Env combined, 24.4 for Np9, and 34.2 for Rec (Figure 4C). Frequencies also differed among superpopulations. Figure 4D shows the largest observed range for each annotation. Substitution and frameshift maps locate the underlying sequence changes along each protein (Figure S9A–B).

At 7p22.1, the Y195C substitution replaces the conserved tyrosine in the reverse-transcriptase YIDD motif with cysteine. This change was previously identified in HERV-K(C7)/HML-2.HOM and linked to loss of reverse-transcriptase activity (43,44). We asked whether any donor carried a provirus that retained all four canonical ORFs without this known replication barrier. Seven donors did. Another 259 met the four-intact-ORF criterion only with Y195C. The remaining 26 met neither criterion (Table S10). Reus et al. previously identified an intact YXDD motif in HML-2.HOM polymerase sequences from two human DNA samples (45).

Across all loci and donor haplotypes, we assigned each retained proviral copy to its longest consecutive Gag–Pro–Pol annotation combination. This gave 2,336 Gag-only, 1,460 Gag–Pro, and 1,196 Gag–Pro–Pol copies. Ten Type-I copies at 1q22 also passed the screen for theoretical Type-I Env (Figure S10A). An independent scan for uninterrupted reading frames examined possible products distinct from the canonical polyproteins (Figure S10B, Supplemental Results).

**Figure 4:**
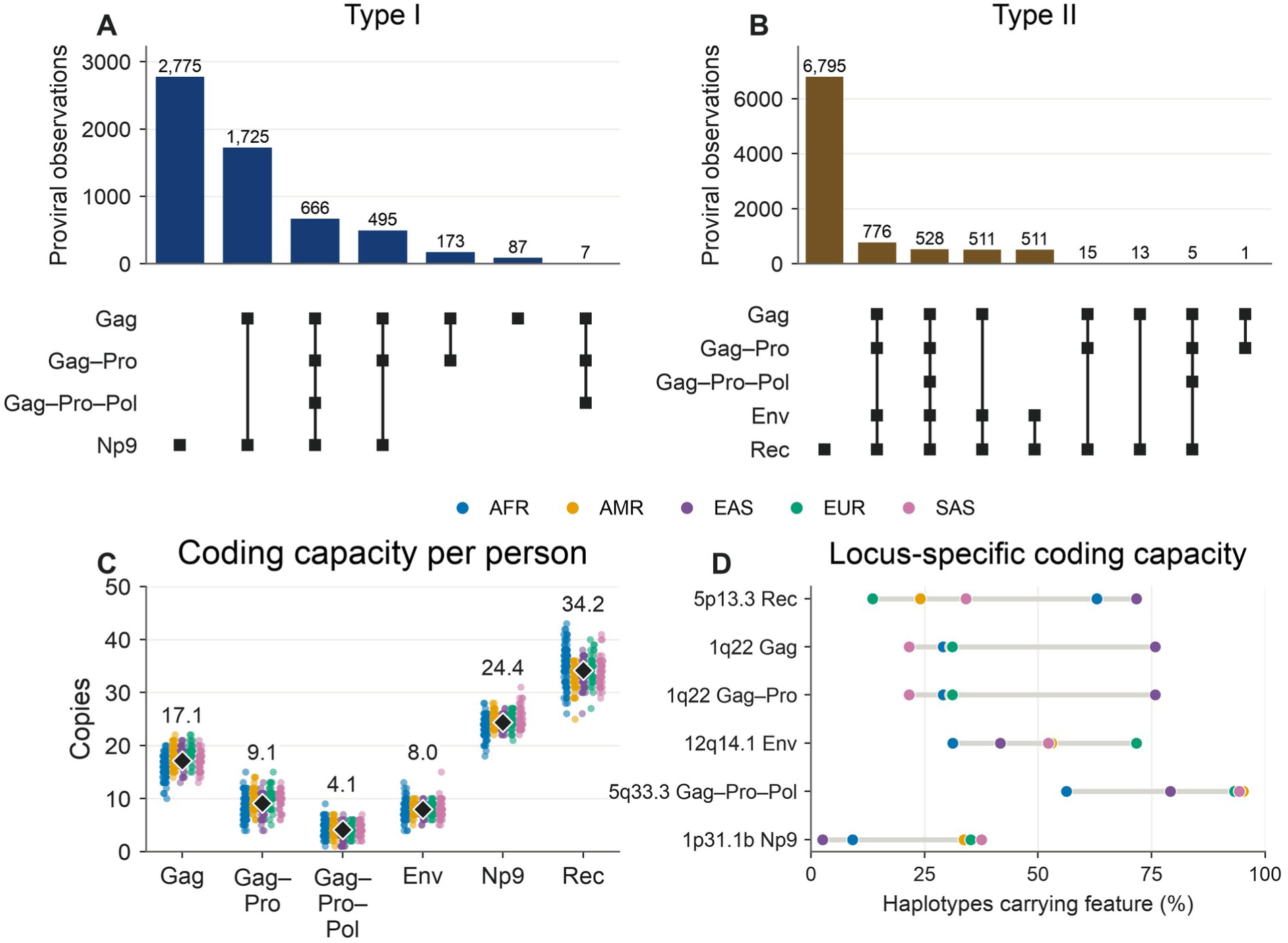
ORF annotations within proviruses and across individuals. (A–B) Combinations of the displayed ORF annotations in Type-I and Type-II proviruses. (C) Combined-screen copy counts per person. Env pools Type-II Env and theoretical N-terminally truncated Type-I Env, whose translation has not been demonstrated. Diamonds show means across 292 donors. (D) Frequencies at loci showing the largest superpopulation differences. Alt text: Plots compare ORF combinations by provirus type, copy counts per donor and population frequencies. Np9-only combinations predominate in Type I and Rec-only combinations in Type II.

### Type-I deletion variation and propagation models

The HML-2 Type-I deletion removes part of the integrase C-terminal domain, the Env signal peptide, and the splice donor used to generate rec. Alternative splicing can instead generate np9 (2). To understand how this deletion became widespread despite its effects on viral coding and splicing, we compared Type-I and Type-II sequences and tested whether both types occurred at the same insertion site. The sequence comparison used the 1,001 bp flanking the deletion and six windows elsewhere in the provirus (Figure 5A).

All directly typed proviruses assigned to LTR5A or LTR5B were Type II. LTR5Hs contained both Type-I and Type-II proviruses (Figure 5B). We found that all 15 human Type-I representatives carried thymine (T) at both zero-based KCON positions 6331 and 6492 immediately upstream of Δ292. The common deletion boundary and these linked flanking variants support a shared ancestral segment. We call this deletion-linked segment the Type-I cassette. Earlier phylogenetic analysis likewise supported a single origin of Δ292 and dispersal of the deletion-bearing region by recombination or gene conversion (46). Among these representatives, mean pairwise divergence was 2.73% in the 1,001 bases flanking the deletion, compared with 2.97–6.44% in six 1-kb windows elsewhere in the provirus.

Divergence in these flanks was 9.98% among all 29 callable Type-II representatives, but 3.68% after restriction to the 14 callable LTR5Hs Type-II representatives (Figure 5C, Table S11). The adjacent upstream window had a similar Type-I divergence of 2.97%. Across 20 loci with at least one Type-I allele, all 9,733 cassette-callable haplotype-locus records were Type I and none were Type II. These included the single 8q11.23 provirus (Figure S11, Table S12). Relative to the rest-of-provirus comparison, the nearest Type-II neighbor changed for 13/15 representatives in the concatenated cassette-flank window (KCON 6000–6501 and 6793–7293) and 11–14/15 in windows B1–B6 (Figure S12A). Δ292 removes KCON positions 6501–6792 (all coordinates zero-based). The linked T/T pair occurred in 25/26 nonhuman primate HML-2 sequence clusters carrying Δ292 and 4/57 without it (Figure S12B). Figure S12C compares the number of ancestral integrations carrying each deletion.

Frequency logos and the complete human and nonhuman primate alignments show the conserved sites and surrounding variation (Figure S13, Table S11). Of note, the fixed window used to compare sequence divergence does not define the cassette’s biological endpoints.

Among the 19 well-sampled loci with at least one Type-I allele, no cassette-callable chromosome carried a Type-II allele. We tested whether recurrent neutral conversion after integration could account for this absence of mixed alleles. The central neutral-conversion scenario predicted 6.8 mixed loci among the 16 age-annotated Type-I-positive loci, compared with none observed. Conditional on Type-I detection at these loci, the upper-bound probability of no mixed samples was 5.47 × 10⁻⁶. Doubling the older age bounds increased this probability to 3.48 × 10⁻⁵. The largest bound across all 54 rate, population-size, and age scenarios was 0.0317 (Table S13).

To understand why Δ292 is so widespread, we compared its distribution with six other large deletions detected in orangutan, siamang, and macaque HML-2 sequences. These deletions ranged from 74 to 2,255 bp and affected the LTR and leader, gag, pol, or the pol-env boundary. Each was observed at one insertion. By comparison, 26 Δ292-bearing nonhuman-primate sequences represented 16 resolved ancestral integrations and one possible additional integration. We compared the host DNA on both sides of each provirus to recognize the same inherited insertion across species and count it once (Figure S12C). We tested whether this unequal distribution required a Δ292-specific propagation advantage after allowing cumulative contributions to differ among deletion classes. For the conservative Δ292 count of 16, the Bayes factor for an added advantage fell from 2.46 × 10⁵ to 26.9 and 1.51 as the permitted variation increased (Figure 5D). Evidence for an added effect therefore depended strongly on the assumed variation in cumulative contributions.

This conclusion also held under broader assumptions about relative detection probabilities and inclusion of the possible additional Δ292-bearing integration (Figure S14A). In the model without a deletion-specific effect, greater allowed variation increased the median cumulative contribution assigned to the Δ292 class from 1.63 to 14.8 times the mean contribution assigned to the other six deletion classes (Figure S14B).

**Figure 5:**
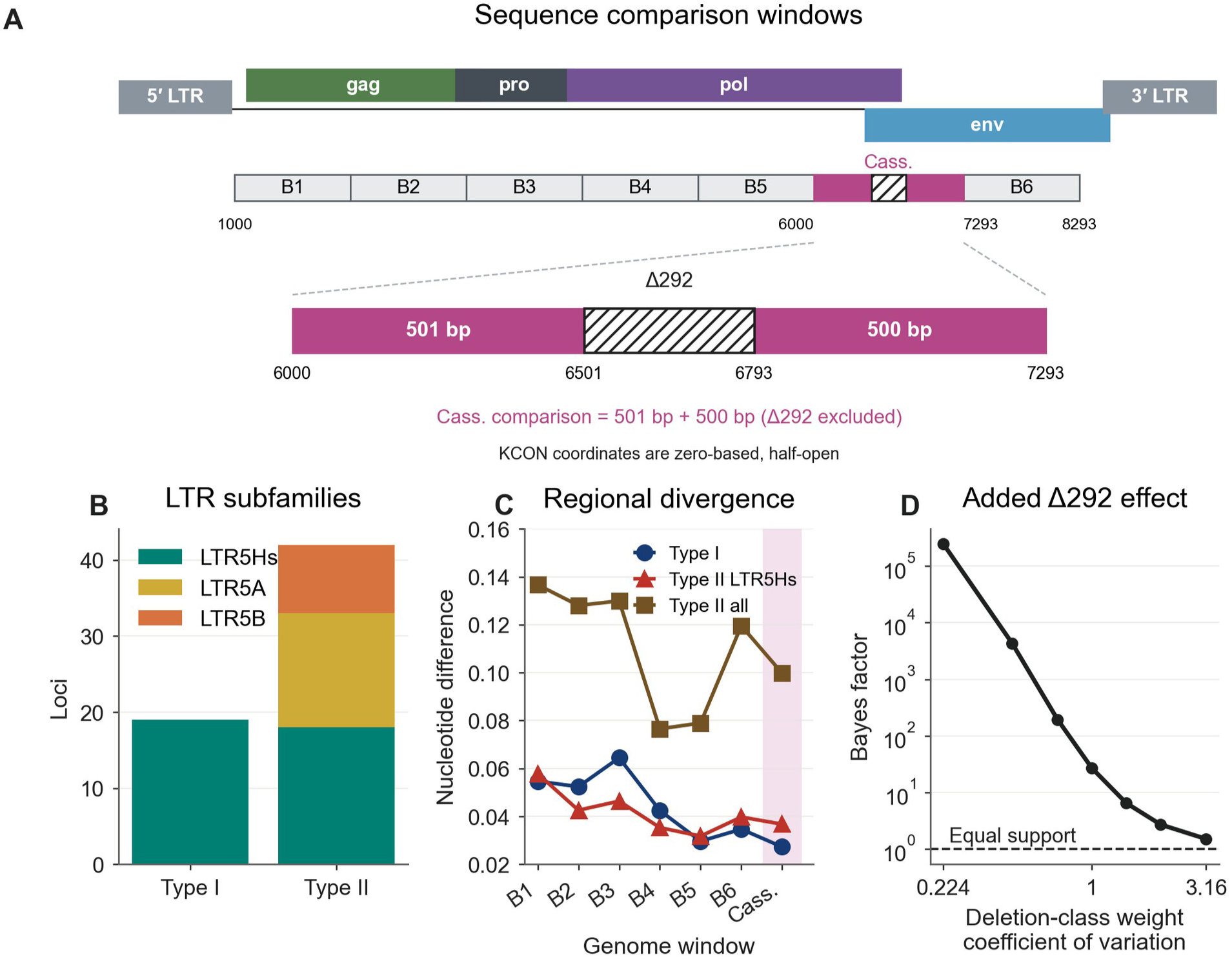
Sequence conservation and models of Type-I propagation. **(A)** KCON comparison windows and Δ292. Magenta segments form the cassette window. Coordinates are zero-based and half-open. **(B)** LTR subfamilies by element type. **(C)** Mean pairwise nucleotide differences in six 1-kb windows and the cassette flanks (Cass.). The y-axis shows the proportion of differing nucleotides. Counts per window are 15 Type-I, 29–30 Type-II, and 14–15 LTR5Hs Type-II representatives (Table S11). **(D)** Evidence for an added Δ292 propagation advantage under increasing variation in cumulative contributions among deletion classes. The dashed line marks equal model evidence. Alt text: A schematic marks the Type-I deletion. Plots compare LTR families and regional divergence, and show that evidence for a deletion-specific advantage falls as unequal cumulative contributions are allowed.

### A rare provirus retains internal coding sequence at a reference solo-LTR locus

A solo-LTR in the reference genome can conceal a full provirus carried by other individuals. Short reads spanning the shared host–LTR junctions cannot distinguish those structures. Therefore, we searched the long-read assemblies for retained internal sequence at reference solo-LTR sites and identified a two-LTR provirus at 8q11.23 in one of 584 haplotypes. The other 583 carried a solo-LTR with the same CACAC TSDs (Figure 6A). Pro and theoretical Type-I Env passed the combined ORF screen. Gag and Np9 contained premature stops, and Pol had a premature stop and a frameshift at residue 874. The two 968-bp LTRs were identical.

We recovered sequences for all 583 solo-LTR observations. The most common sequence occurred 547 times (93.8%), and each of 13 rarer sequences differed from it by one nucleotide. These sequences formed a star-like network across superpopulations (Figure 6B). Mean pairwise nucleotide diversity was 0.000127 per base across the 968-bp LTR, equivalent to 0.123 differences per sequence pair. Among twelve comparison loci, median diversity was 0.00102 per base. Two loci, 3q13.2 and 12q14.1, had lower diversity than 8q11.23 (Figure S15A, Table S4).

Both host–LTR junctions were supported in the Vindija, Altai, and Chagyrskaya Neanderthal genomes and the Denisovan genome, establishing that the insertion site was already occupied in archaic hominins (Table S14). Importantly, however, these junction reads do not distinguish a solo-LTR from a full provirus. The corresponding chimpanzee and gorilla assembly sites contained an empty pre-integration site matching the flanking sequences of this provirus in hominids.

The identical paired LTRs and low solo-LTR diversity would suggest a recent origin under a simple substitution clock (Figure S15B). However, the insertion is present in all sampled modern haplotypes and in archaic hominins, supporting an older history.

**Figure 6:**
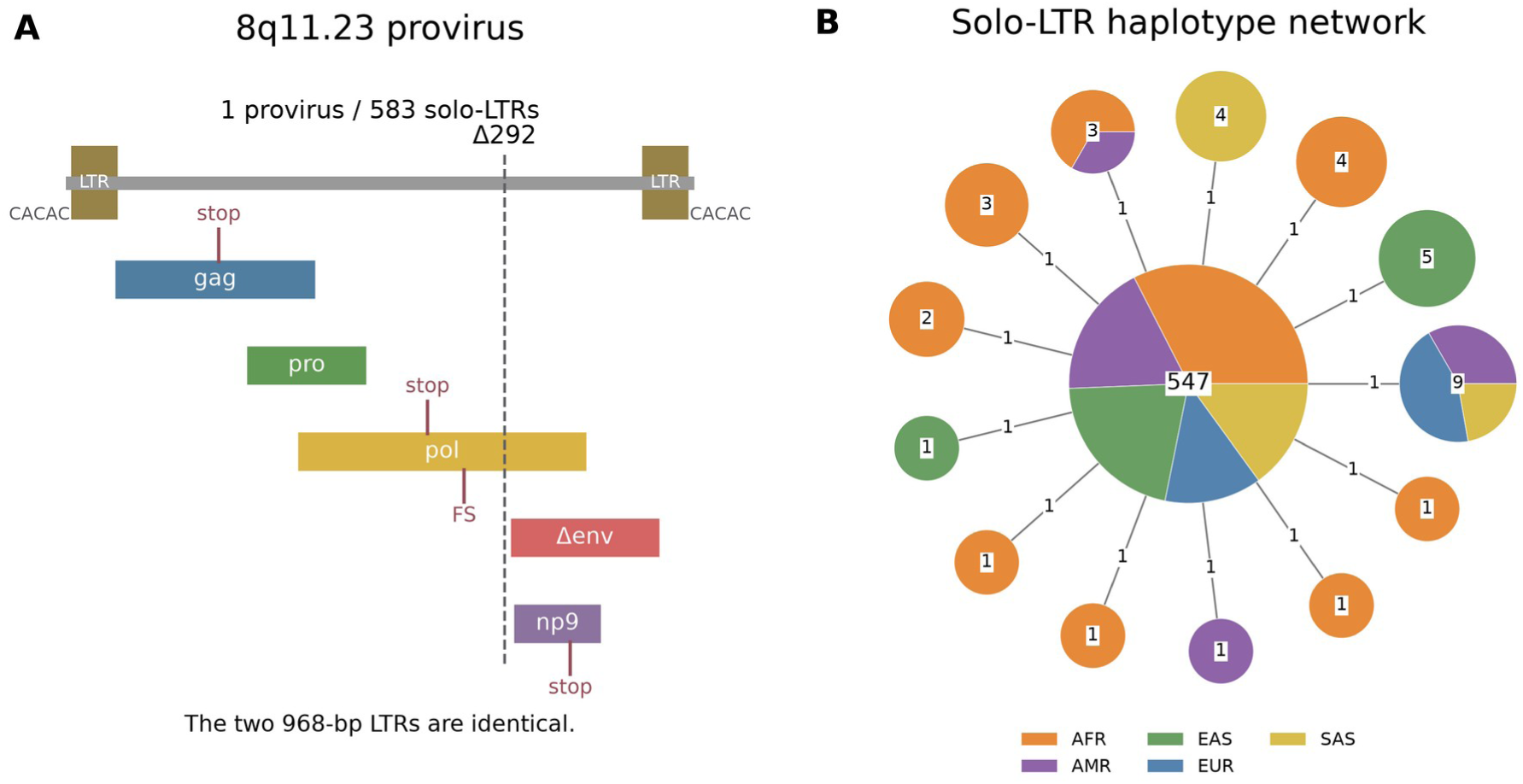
Structure and variation at 8q11.23. **(A)** Sequence-predicted proviral structure. Labels mark disruptive variants, Δ292 and the shared CACAC target-site duplication. FS denotes a frameshift. Δenv denotes theoretical Type-I Env, an N-terminally truncated product whose translation has not been demonstrated. **(B)** Network of 14 solo-LTR sequence haplotypes from 583 observations. Labels give counts. Colors show superpopulation composition. Each spoke represents one nucleotide substitution. Alt text: The rare full provirus at 8q11.23 has identical LTRs and disruptive coding variants. A star-shaped network shows one predominant solo-LTR sequence and thirteen one-base variants.

### Structural alleles associate with gene expression, growth and anti-CD20 response

Because we found that HML-2 structural variants could disable viral genes and alter the regulatory sequences, we next explored whether these variants may be linked to functionally important phenotypes in health and disease. We linked these alleles to expression, growth, and treatment-response measurements from the same donors’ lymphoblastoid cell lines to examine their possible functional consequences. The 1p31.1b internal fragment was associated with higher SLC44A5 expression in two nonoverlapping cohorts, MAGE (39 donors) and GEUVADIS (28 donors) (Figure 7A, Table S15). The MAGE association did not pass discovery-wide correction (q=0.127 across 1,289,856 tests). GEUVADIS passed correction across 30 follow-up tests (q=0.0184). Sensitivity estimates using HC3 robust standard errors had P values of 4.90 × 10⁻⁵ and 0.00123, respectively. Supplemental Methods distinguish the discovery and follow-up testing families.

At 6q14.1, solo-LTR carriers grew more slowly than provirus carriers in all four represented populations (Figure 7B). The drug-free growth estimate combined repeated measurements while accounting for experimental variation. Among 34 donors, seven carried the solo-LTR. The superpopulation- and sex-adjusted contrast had P=0.000201 with pedigree-clustered standard errors and q=0.0174 across the 174 estimable tests in the 237-model screen.

At 12q13.2, provirus carriers had higher relative viability after ofatumumab plus human serum than carriers of a solo-LTR but no provirus (Figure 7C). The serum analysis included 49 donors: 33 provirus carriers, 11 solo-LTR carriers without a provirus, and five noncarriers modeled separately. Ofatumumab had regression q=0.0146 and population-stratified permutation q=0.00933. The obinutuzumab association was less robust. Its regression q value was 0.0247, but the permutation q value was 0.0893. The rituximab interval included zero (regression q=0.461), as did the two controls without added human serum.

These associations identify specific structural contrasts for functional experiments: presence or absence of the 1p31.1b internal fragment, and retention or loss of the proviral interior at 6q14.1 and 12q13.2. Recreating these states in matched cell lines could test whether the viral sequence contributes to the phenotype. We report Gag-presence and copy-number comparisons that did not show associations in Supplemental Results and Figure S16.

**Figure 7:**
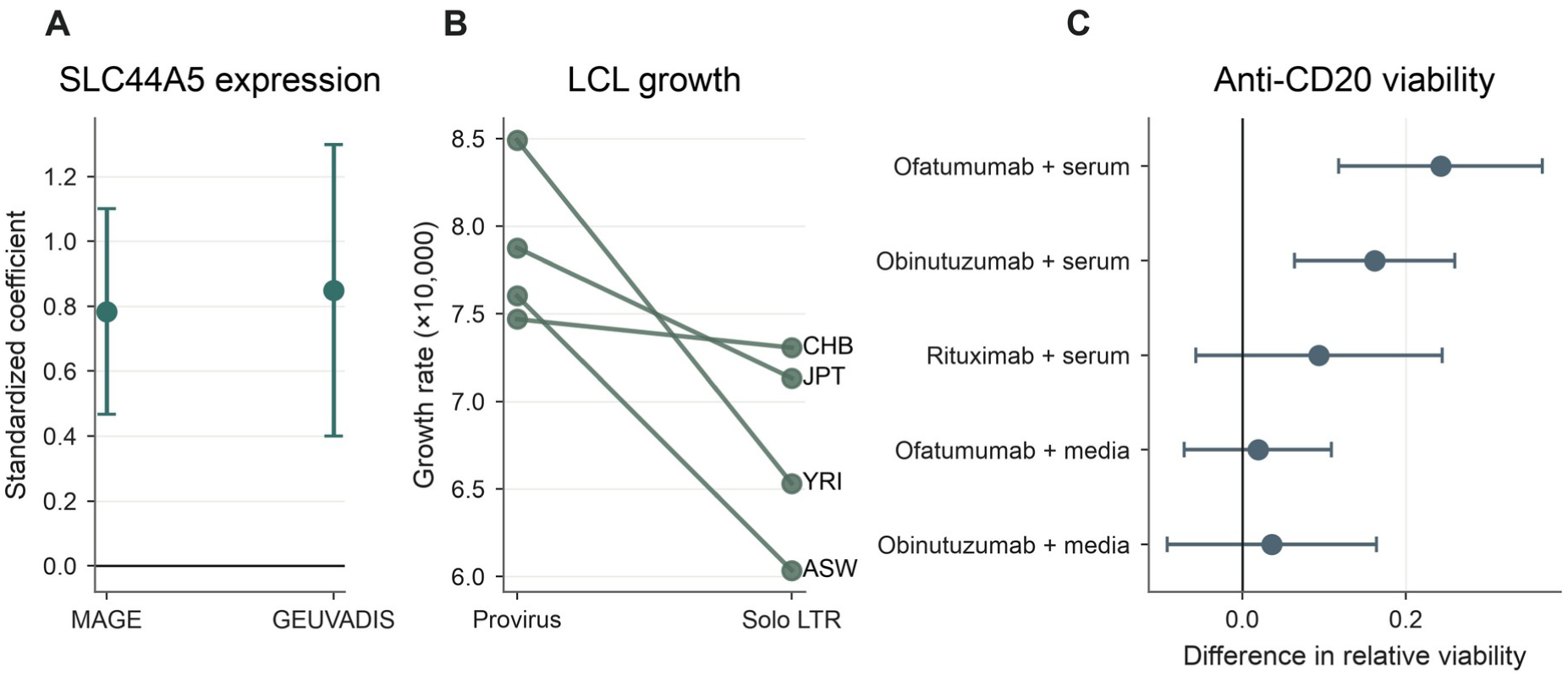
HML-2 structural states and cellular phenotypes. **(A)** Standardized regression coefficients for SLC44A5 expression versus 1p31.1b fragment presence in MAGE and GEUVADIS. **(B)** Mean LCL growth by 6q14.1 state and population. **(C)** Adjusted differences in relative viability for 12q13.2 provirus carriers versus solo-LTR carriers without a provirus. Intervals in A and C are normal-approximation 95% confidence intervals. Alt text: Regression and group-comparison plots show associations of 1p31.1b with SLC44A5 expression, 6q14.1 with cell growth, and 12q13.2 with anti-CD20 response. Uncertainty intervals are shown.

### HML-2 loci differ in chromatin accessibility

Retained ORFs can contribute to expression only if the relevant sequences are accessible to transcriptional machinery. Therefore, we used Fiber-seq to measure accessibility at 21 HML-2 loci in lymphoblastoid cell lines from 39 donors with matched genome assemblies. Accessibility was the fraction of total read coverage assigned to FIREs across the assayed regulatory peaks.

Each locus had 26–77 assayed haplotypes. Median accessibility ranged from 0 to 33.8%, highest at 8p23.1c (Figure S17). Other assayed 8p23.1 loci had medians of 30.8–32.8%. At 1q22, the median across 76 haplotypes was 7.5% (interquartile range 4.9–10.8%). Matched RNA measurements could test whether these differences predict transcription of the retained viral sequences.

## Discussion

Our study shows that the commonly studied HML-2 clade of endogenous retroviruses is far more structurally polymorphic and individually variable than previously appreciated. We uncovered a previously undescribed full-length provirus, documented tandem arrays as a previously unrecognized class of HML-2 variation, and found extensive variation at loci previously thought to be fixed. Viral genes disrupted in reference genomes remained intact in other individuals (Table S17). This hidden heterogeneity changes which viral proteins a person can express and how much of each. It expands the mechanisms by which HML-2 may affect health and disease. The associations between newly identified structural alleles and cellular phenotypes further show why HML-2 expression and function must be studied on a genome-by-genome basis rather than assumed to match standard genomic references.

Tandem arrays have previously been described only at 7p22.1 and 6q14.1 (40,41). These structures occur at 13 loci in our panel and constitute a widespread class of HML-2 structural variation. Individual array members also differ in the coding sequences they carry.

The population data also reveal how this diversity can arise after the original infection. Unequal homologous recombination between misaligned LTRs on separate chromatids can produce a solo-LTR and a tandem duplication as reciprocal products. Crossover between the 5′ LTR of one provirus and the 3′ LTR of the other leaves a solo-LTR on one chromatid and two proviral copies separated by a shared LTR on the other (40). The shared-LTR structures in HML-2 arrays support this mechanism. Tandem arrays remain substrates for further recombination and can contract to smaller arrays or solo-LTRs. Solo-LTR formation permanently removes the internal viral genes from that copy. The remaining LTR cannot regenerate those genes through the reverse rearrangement. This asymmetry may explain the abundance of solo-LTRs and the rarity of most tandem arrays.

We also show that short-read-based methods are inadequate for rigorous examination of HML-2 transcriptional or genotypic associations with health or disease. Our matched variant comparison shows that an absent or inadequate short-read call cannot be treated as the reference allele. Many ORF-associated variants remained unresolved, even though the resolved calls generally agreed with the assemblies (Table 2). Whole-ORF status requires sequence or coverage evidence across the coding region. HML-2 expression studies therefore need to account for the structural and coding variation among individuals.

Sequence comparisons also suggest exchange between elements at different loci. The LTR and Pol sequences of 19p12c and 10q24.2 have different closest relatives, although bootstrap support for those particular relatives is limited (Figure 3A). Shared complete gene sequences at separately assigned loci could reflect duplication or later exchange (Figure 3B). Differences between the two flanking TSDs at five loci also identify possible changes at insertion boundaries. Mutation, exchange, and sequence or boundary error can produce these differences. Matching TSDs do not exclude recombination confined to the viral sequence.

The spread of Δ292 points to a different setting for recombination, during viral replication before integration. Its common boundary and linked flanking variants support shared cassette ancestry across HML-2 loci (Figure 5, Figure S12). If Type-I and Type-II RNAs were copackaged, template switching during reverse transcription could transfer the deletion and adjacent sequence onto different viral backgrounds. Integration of the recombinant DNA would then place the cassette at a new genomic site. After integration, conversion would instead copy it into an occupied site and could leave both types segregating there. The absence of within-locus Type-I/II polymorphism strongly disfavors repeated neutral conversion. Together, these observations support recombination during reverse transcription, as Subramanian et al. (2) proposed.

This mechanism explains how Δ292 could move between viral backgrounds, but raises the question of why it spread so much farther than comparable deletions. Differences in promoter activity or RNA packaging could affect source output (47,48). The deletion could also change RNA allocation or the cost of producing viral products. Type-I propagation depends on other viral genomes supplying the functions disrupted by Δ292. These roles could coexist. Type-I genomes may exploit Type-II HML-2 replication functions to spread and compete with Type II for the production of new proviruses. Such ‘piggybacking’ has been proposed for the barley retroelement BARE-2 (49). We hypothesize that this competition could also reduce intact Type-II integration and provide a last line of defense after infection had already occurred. Our analysis cannot distinguish an advantage caused by Δ292 from linked viral traits or historical opportunities for propagation, all of which could contribute to the Type-I deletion’s prevalence.

We also discovered a previously undescribed full-length HML-2 provirus at 8q11.23. Fascinatingly, the insertion is fixed in humans but persists almost entirely as a solo-LTR of remarkably low sequence diversity. The one full-length allele identified has identical LTRs. One possible explanation is that the internal viral genes were initially retained through exaptation.

This could have allowed the provirus to spread to fixation before those genes became maladaptive. For example, a viral product could have acted as a restriction factor against an exogenous virus, then become harmful after that virus disappeared. Selection against the proviral interior would then favor solo-LTR alleles and reduce variation at the locus. Recombination becomes less frequent as the LTRs diverge (50), so this process could preferentially remove interiors from proviruses that retained identical ancestral LTRs.

We also show that the coding-potential diversity we identify here may have meaningful biological consequences. Our functional screens connect this evolutionary variation to present-day cellular traits. The association between the 1p31.1b fragment and SLC44A5 expression in two independent cohorts in particular identifies a candidate for testing nearby gene regulation. The 6q14.1 growth and 12q13.2 anti-CD20 associations distinguish a provirus from a solo-LTR (Figure 7). Recreating these states in matched cell lines would separate element effects from linked host variants. Our Fiber-seq analysis adds a complementary measure of chromatin accessibility at individual loci and haplotypes (Figure S17), but does not directly measure transcription. Matching these measurements to RNA expression would help determine whether the retained sequences are transcribed. Our functional analyses also use relatively small matched subsets, so the associations we have identified will need to be tested in larger cohorts.

Long-read pangenomes reveal extensive structural and coding variation that changes which HML-2 products each person can express. This previously hidden variability expands the mechanisms by which HML-2 may affect health and disease and establishes the need to study endogenous retroviruses on a genome-by-genome basis.

## Data Availability

Code and derived data (v0.4.3) are archived at https://doi.org/10.5281/zenodo.22907293. Analysis code is also available at https://github.com/dmworstell/hml2-pangenome. The input files and implementation for the ORF-associated variant comparison accompany Table S3 in Supplementary Data. Oxford Nanopore reads were obtained from the public human-pangenomics archive using the HPRC data_ont_pre_release.index.csv source index. The matching sample IDs, file locations, SRA accessions and BioProjects (PRJNA701308 and PRJNA731524 among them) are provided in Table S2 supporting data. Long-read assemblies and pangenome graphs were obtained from the Human Pangenome Reference Consortium (HPRC releases 1 and 2, https://humanpangenome.org/data/, and Release 2 assemblies and versioned indices at https://github.com/human-pangenomics/hprc_intermediate_assembly) and the Human Genome Structural Variation Consortium (HGSVC3, https://www.hgsvc.org). The short-read comparison used high-coverage 1000 Genomes Project VCF data distributed through the International Genome Sample Resource (https://www.internationalgenome.org) (23). MAGE expression matrices and covariates are archived at https://doi.org/10.5281/zenodo.10535719.

Source RNA-seq reads are under BioProject PRJNA851328. The GEUVADIS expression matrix (GEUVADIS_gene_RPKM_50FN_resk10.txt.gz) and sample metadata are available under ArrayExpress/BioStudies accession E-GEUV-1. EBV phenotypes were obtained from the supplementary datasets of Mandage et al. (https://doi.org/10.1371/journal.pone.0179446.s003) and Houldcroft et al. (https://doi.org/10.1371/journal.pone.0108384.s003), and growth phenotypes from Im et al. (https://doi.org/10.1371/journal.pgen.1002525.s001). Anti-CD20 phenotypes were obtained from the supplementary data accompanying Small et al. (https://doi.org/10.3390/cells12121574).

## Acknowledgements

We carried out all analyses at Tufts University, Graduate School of Biomedical Sciences. We thank Michael P. Rist for his contributions and all current and former members of the John Coffin Laboratory for suggestions and revisions. We created the schematic in Figure 1A with BioRender. OpenAI ChatGPT/Codex, Google Gemini, and Anthropic Claude assisted with code development. ChatGPT/Codex also assisted with manuscript edits, schematic figures, and the graphical abstract, under D.A.M.-W.’s direction. D.A.M.-W. reviewed and revised all generated material.

## Author Contributions

D.A.M.-W. and J.M.C. conceived and designed the study. D.A.M.-W. developed the analysis pipelines, performed the computational analyses, and wrote the manuscript. M.F. contributed to experimental validation. A.M. and A.B.S. contributed Fiber-seq data and chromatin accessibility analysis. A.M. and M.R.V. contributed to generating the Fiber-seq data table. J.M.C. supervised the study and edited the manuscript.

## References

1. Lander, E.S., Linton, L.M., Birren, B. et al. (2001) Initial sequencing and analysis of the human genome. Nature, 409, 860–921. 10.1038/35057062

2. Subramanian, R.P., Wildschutte, J.H., Russo, C. et al. (2011) Identification, characterization, and comparative genomic distribution of the HERV-K (HML-2) group of human endogenous retroviruses. Retrovirology, 8, 90. 10.1186/1742-4690-8-90

3. Garcia-Montojo, M., Doucet-O’Hare, T., Henderson, L. et al. (2018) Human endogenous retrovirus-K (HML-2): a comprehensive review. Crit Rev Microbiol, 44, 715–738. 10.1080/1040841x.2018.1501345

4. Grow, E.J., Flynn, R.A., Chavez, S.L. et al. (2015) Intrinsic retroviral reactivation in human preimplantation embryos and pluripotent cells. Nature, 522, 221–225. 10.1038/nature14308

5. Liu, X., Liu, Z., Wu, Z. et al. (2023) Resurrection of endogenous retroviruses during aging reinforces senescence. Cell, 186, 287–304.e26. 10.1016/j.cell.2022.12.017

6. Li, W., Lee, M.-H., Henderson, L. et al. (2015) Human endogenous retrovirus-K contributes to motor neuron disease. Sci Transl Med, 7, 307ra153. 10.1126/scitranslmed.aac8201

7. Gleason, C., Terry, S.N., Hernandez, M.M. et al. (2025) An integrated approach for the accurate detection of HERV-K HML-2 transcription and protein synthesis. Nucleic Acids Res, 53, gkaf011. 10.1093/nar/gkaf011

8. Murimi-Worstell, D.A., Murimi-Worstell, I.B., Roy, F.M. et al. (2025) In-depth analysis of endogenous retrovirus expression in glioblastoma. Mob DNA, 16, 29. 10.1186/s13100-025-00365-w

9. Bendall, M.L., de Mulder, M., Iñiguez, L.P., et al. (2019) Telescope: Characterization of the retrotranscriptome by accurate estimation of transposable element expression. PLoS Comput Biol, 15, e1006453. 10.1371/journal.pcbi.1006453

10. Macfarlane, C.M. and Badge, R.M. (2015) Genome-wide amplification of proviral sequences reveals new polymorphic HERV-K(HML-2) proviruses in humans and chimpanzees that are absent from genome assemblies. Retrovirology, 12, 35. 10.1186/s12977-015-0162-8

11. Wildschutte, J.H., Williams, Z.H., Montesion, M. et al. (2016) Discovery of unfixed endogenous retrovirus insertions in diverse human populations. Proc Natl Acad Sci U S A, 113, E2326–34. 10.1073/pnas.1602336113

12. Liao, W.-W., Asri, M., Ebler, J. et al. (2023) A draft human pangenome reference. Nature, 617, 312–324. 10.1038/s41586-023-05896-x

13. Lucas, J.K., Hebbar, P., Liao, W.-W. et al. (2026) HPRC2: A human pangenome reference with near-complete coverage of common genetic variation. bioRxiv [preprint], 2026.07.21.739710v1. 10.64898/2026.07.21.739710

14. Logsdon, G.A., Ebert, P., Audano, P.A. et al. (2025) Complex genetic variation in nearly complete human genomes. Nature, 644, 430–441. 10.1038/s41586-025-09140-6

15. Nurk, S., Koren, S., Rhie, A. et al. (2022) The complete sequence of a human genome. Science, 376, 44–53. 10.1126/science.abj6987

16. Guarracino, A., Heumos, S., Nahnsen, S. et al. (2022) ODGI: understanding pangenome graphs. Bioinformatics, 38, 3319–3326. 10.1093/bioinformatics/btac308

17. Li, H. (2018) Minimap2: pairwise alignment for nucleotide sequences. Bioinformatics, 34, 3094–3100. 10.1093/bioinformatics/bty191

18. Danecek, P., Bonfield, J.K., Liddle, J. et al. (2021) Twelve years of SAMtools and BCFtools. Gigascience, 10, giab008. 10.1093/gigascience/giab008

19. Lee, Y.N. and Bieniasz, P.D. (2007) Reconstitution of an infectious human endogenous retrovirus. PLoS Pathog, 3, e10. 10.1371/journal.ppat.0030010

20. Quinlan, A.R. and Hall, I.M. (2010) BEDTools: a flexible suite of utilities for comparing genomic features. Bioinformatics, 26, 841–842. 10.1093/bioinformatics/btq033

21. Katoh, K. and Standley, D.M. (2013) MAFFT multiple sequence alignment software version 7: improvements in performance and usability. Mol Biol Evol, 30, 772–780. 10.1093/molbev/mst010

22. Cock, P.J.A., Antao, T., Chang, J.T. et al. (2009) Biopython: freely available Python tools for computational molecular biology and bioinformatics. Bioinformatics, 25, 1422– 1423. 10.1093/bioinformatics/btp163

23. Byrska-Bishop, M., Evani, U.S., Zhao, X. et al. (2022) High-coverage whole-genome sequencing of the expanded 1000 Genomes Project cohort including 602 trios. Cell, 185, 3426–3440.e19. 10.1016/j.cell.2022.08.004

24. Ebert, P., Audano, P.A., Zhu, Q. et al. (2021) Haplotype-resolved diverse human genomes and integrated analysis of structural variation. Science, 372, eabf7117. 10.1126/science.abf7117

25. Ebler, J., Ebert, P., Clarke, W.E. et al. (2022) Pangenome-based genome inference allows efficient and accurate genotyping across a wide spectrum of variant classes. Nat Genet, 54, 518–525. 10.1038/s41588-022-01043-w

26. Saitou, N. and Nei, M. (1987) The neighbor-joining method: a new method for reconstructing phylogenetic trees. Mol Biol Evol, 4, 406–425. 10.1093/oxfordjournals.molbev.a040454

27. Stergachis, A.B., Debo, B.M., Haugen, E. et al. (2020) Single-molecule regulatory architectures captured by chromatin fiber sequencing. Science, 368, 1449–1454. 10.1126/science.aaz1646

28. Vollger, M.R., Swanson, E.G., Neph, S.J. et al. (2026) Somatic epimutations cap genetic determinism in the human diploid chromatin epigenome. bioRxiv [preprint], 2024.06.14.599122v3. 10.1101/2024.06.14.599122

29. Mandage, R., Telford, M., Rodríguez, J.A. et al. (2017) Genetic factors affecting EBV copy number in lymphoblastoid cell lines derived from the 1000 Genome Project samples. PLoS One, 12, e0179446. 10.1371/journal.pone.0179446

30. Houldcroft, C.J., Petrova, V., Liu, J.Z. et al. (2014) Host genetic variants and gene expression patterns associated with Epstein-Barr virus copy number in lymphoblastoid cell lines. PLoS One, 9, e108384. 10.1371/journal.pone.0108384

31. Im, H.K., Gamazon, E.R., Stark, A.L. et al. (2012) Mixed effects modeling of proliferation rates in cell-based models: consequence for pharmacogenomics and cancer. PLoS Genet, 8, e1002525. 10.1371/journal.pgen.1002525

32. Taylor, D.J., Chhetri, S.B., Tassia, M.G. et al. (2024) Sources of gene expression variation in a globally diverse human cohort. Nature, 632, 122–130. 10.1038/s41586-024-07708-2

33. Lappalainen, T., Sammeth, M., Friedländer, M.R. et al. (2013) Transcriptome and genome sequencing uncovers functional variation in humans. Nature, 501, 506–511. 10.1038/nature12531

34. Small, G.W., Akhtari, F.S., Green, A.J. et al. (2023) Pharmacogenomic Analyses Implicate B Cell Developmental Status and MKL1 as Determinants of Sensitivity toward Anti-CD20 Monoclonal Antibody Therapy. Cells, 12, 1574. 10.3390/cells12121574

35. Benjamini, Y. and Hochberg, Y. (1995) Controlling the False Discovery Rate: A Practical and Powerful Approach to Multiple Testing. Journal of the Royal Statistical Society. Series B (Methodological), 57, 289–300. 10.1111/j.2517-6161.1995.tb02031.x

36. Prufer, K., de Filippo, C., Grote, S., et al. (2017) A high-coverage Neandertal genome from Vindija Cave in Croatia. Science, 358, 655–658. 10.1126/science.aao1887

37. Prufer, K., Racimo, F., Patterson, N. et al. (2014) The complete genome sequence of a Neanderthal from the Altai Mountains. Nature, 505, 43–9. 10.1038/nature12886

38. Mafessoni, F., Grote, S., de Filippo, C., et al. (2020) A high-coverage Neandertal genome from Chagyrskaya Cave. Proc Natl Acad Sci U S A, 117, 15132–15136. 10.1073/pnas.2004944117

39. Meyer, M., Kircher, M., Gansauge, M.-T. et al. (2012) A high-coverage genome sequence from an archaic Denisovan individual. Science, 338, 222–226. 10.1126/science.1224344

40. Hughes, J.F. and Coffin, J.M. (2004) Human endogenous retrovirus K solo-LTR formation and insertional polymorphisms: implications for human and viral evolution. Proc Natl Acad Sci U S A, 101, 1668–1672. 10.1073/pnas.0307885100

41. Pasternack, N., Paulsen, O. and Nath, A. (2025) Characterization of novel human endogenous retrovirus structures on chromosomes 6 and 7. Front Genet, 16, 1498978. 10.3389/fgene.2025.1498978

42. Guarracino, A., Buonaiuto, S., de Lima, L.G., et al. (2023) Recombination between heterologous human acrocentric chromosomes. Nature, 617, 335–343. 10.1038/s41586-023-05976-y

43. Tönjes, R.R., Czauderna, F. and Kurth, R. (1999) Genome-Wide Screening, Cloning, Chromosomal Assignment, and Expression of Full-Length Human Endogenous Retrovirus Type K. J Virol, 73, 9187–9195. 10.1128/JVI.73.11.9187-9195.1999

44. Berkhout, B., Jebbink, M. and Zsíros, J. (1999) Identification of an Active Reverse Transcriptase Enzyme Encoded by a Human Endogenous HERV-K Retrovirus. J Virol, 73, 2365–2375. 10.1128/JVI.73.3.2365-2375.1999

45. Reus, K., Mayer, J., Sauter, M. et al. (2001) Genomic Organization of the Human Endogenous Retrovirus HERV-K(HML-2.HOM) (ERVK6) on Chromosome 7. Genomics, 72, 314–320. 10.1006/geno.2000.6488

46. Belshaw, R., Pereira, V., Katzourakis, A. et al. (2004) Long-term reinfection of the human genome by endogenous retroviruses. Proc Natl Acad Sci U S A, 101, 4894–4899. 10.1073/pnas.0307800101

47. Bhardwaj, N., Montesion, M., Roy, F. et al. (2015) Differential expression of HERV-K (HML-2) proviruses in cells and virions of the teratocarcinoma cell line Tera-1. Viruses, 7, 939–968. 10.3390/v7030939

48. Montesion, M., Williams, Z.H., Subramanian, R.P. et al. (2018) Promoter expression of HERV-K (HML-2) provirus-derived sequences is related to LTR sequence variation and polymorphic transcription factor binding sites. Retrovirology, 15, 57. 10.1186/s12977-018-0441-2

49. Tanskanen, J.A., Sabot, F., Vicient, C. and Schulman, A.H. (2007) Life without GAG: The BARE-2 retrotransposon as a parasite’s parasite. Gene, 390, 166–174. 10.1016/j.gene.2006.09.009

50. Belshaw, R., Watson, J., Katzourakis, A. et al. (2007) Rate of recombinational deletion among human endogenous retroviruses. J Virol, 81, 9437–9442. 10.1128/jvi.02216-06

