## Supplementary information for "Structural polymorphism and population-variable coding capacity of HERV-K(HML-2) in human pangenomes"

### Supplemental Methods

#### Reference validation of graph-extracted elements

We checked graph-extracted elements against the reference assembly used for their locus coordinates (T2T-CHM13 or GRCh38). The reference search window extended 6 Mb on either side of the locus, clipped to chromosome bounds. We selected the best alignment by matching base count. An off-locus alignment of at least 2 kb and at least 97% identity triggered rejection when its midpoint lay outside the locus interval expanded by 5 kb. When the body did not localize the element, the check used 300-bp outer host-flank probes and alignments of at least 150 bp. An off-locus best flank hit also triggered rejection. Missing or inconclusive reference-alignment evidence did not itself trigger rejection. The flank-anchored degraded loci 8p22 and 17p13.1 were exempted from this check.

#### Structural classification

We identified HML-2 boundaries using minimap2 asm20 alignments to KCON with --score-N=0 and --secondary=no. KCON-hit filtering allowed MAPQ 0. Structural-boundary detection retained alignment blocks of at least 250 bp. Multiple traversals of KCON positions 1500–7499 (zero-based) were detected using blocks of at least 700 bp and reference coverage depth of at least two. A gap of more than 3,000 bp between HML-2 blocks separated independent copies from a contiguous array. LTR-chain splitting required LTR alignments longer than 750 bp and MAPQ>40. For non-array internal-bearing sequences, we defined a provirus as coverage of at least 80% of the type-matched KCON reference. Lower coverage defined a fragment. Coverage was the fraction of reference positions aligned to non-gap sample bases.

We extracted graph subgraphs with odgi extract -c 10 --optimize, sorted with -p Y, groomed, pruned with -c 1, chopped to 8-bp nodes, and sorted with -O -p cb before path extraction.

We verified extracted sequences against their source FASTAs and native pangenome paths using donor, haplotype, and extraction interval, and recorded source-file checksums. For graph-derived sequences, we aligned continuous native paths to T2T-CHM13 and GRCh38 using minimap2 asm20 and base-resolved CIGAR output. A candidate required a single physical interval supported by a primary alignment with MAPQ at least 20, at least 95% identity across paired alignment bases, at least 4,500 aligned bases in each 5-kb host flank, and at least 450 aligned bases in each adjacent 500-bp flank. Structural insertion and deletion columns did not enter the nucleotide-identity denominator. Intervals crossing an alignment skip were rejected. We analyzed accepted sequences with the same structural and ORF caller. We checked overlapping source intervals for repeated observations before counting them as separate copies.

The search for nonreference proviral sequence retained HML-2 sequences longer than 5,000 bp without a corresponding T2T proviral structure. Candidates were compared with known occupied sites to distinguish a newly identified structural allele from a new insertion site.

#### Structural-state denominators and locus assignment

The sex-chromosome totals comprised two X copies from each of 144 female donors and one X and one Y from each of 144 male donors. Four donors with unknown sex were excluded from the sex-chromosome totals. We assigned X/Y assembly partitions using sample sex metadata and sequence-supported chromosome assignments. A call required a retained provirus, solo-LTR, fragment, or multi-copy observation assigned to that locus, or an explicit noncarrier call. Figure 1 and Tables S1/S5 report called and total chromosome counts separately. A missing row or a record moved to a different physical locus did not create a noncarrier call.

Direct alignment to the published HML-11 sequence panel (1) gave 98.44% identity over 95.19% of the 8p22 query and 99.18% identity over the full 17p13.1 query.

#### ORF integrity scoring

ORF coordinates refer to KCON (2) and its MAFFT alignment (3). We reconstructed rec from KCON bases 6451–6711 and 8411–8467 (1-based, inclusive), including its terminal stop codon. We applied start-codon checks to gag, env, np9, and rec. Type-I Env annotations describe the remaining env reading frame after Δ292 relative to the type-matched reference. A passing call does not restore the deleted Env signal peptide or demonstrate translation of the theoretical N-terminally truncated product. Each gag, pro, and pol region was scored in its own annotated frame, so the canonical programmed frame transitions were not treated as disruptive sequence mutations. We classified a premature stop before the final translated residue as Nonsense or Frameshift according to the position of the first frameshift. Intact calls had no additional frameshift or premature stop and met the applicable start-codon and length criteria. The combined screen also included Intact_FS_End calls, the stop-free altered-frame candidates defined in the main Methods. Fragment_Intact calls were excluded. For Figure S4A, we grouped proviral copies by locus and haplotype, ordered them by recorded part number, and required unique, contiguous positions. Intact and Intact_FS_End were the two passing categories present among eligible copies. Missing and Undetermined annotations were excluded from the gene-position denominator. Figure S4B uses array-bearing haplotypes as its denominator.

For each locus, variant counts included retained Provirus and Provirus_from_Multi records from public donors. We excluded reference sequences and rejected assembly artifacts and duplicate locus labels. Amino-acid positions were parsed from explicit substitution annotations, and frameshift starts from Frameshift_at annotations. We counted each copy once per variant. The denominator was the number of retained proviral copies at the named locus. In-frame indel annotations were retained in the source table but were not classified as amino-acid substitutions (Figure S9).

For the ORF-combination and longest-reading-frame analyses in Figure S10, we used retained proviral copies from public donors. We excluded reference sequences, assembly artifacts, duplicate records for the same assembled interval, and HML-11 elements. For Figure S10A, we pooled Intact and Intact_FS_End calls and classified each copy by its longest consecutive combination of passing Gag, Pro, and Pol annotations. Type-I copies passing all three genes and the theoretical N-terminally truncated Env annotation formed a fourth category. Translation of that Env product has not been demonstrated. For the independent reading-frame scan in Figure S10B, we aligned retained source sequences to the type-specific KCON reference using the same ORF caller. We removed alignment gaps and retained nucleotide insertions and deletions in each copy.

In each of the three forward frames, we followed ATG-starting segments to an in-frame stop, an ambiguous codon, or the sequence end. We resolved ties by retaining the first frame and then the first ATG encountered. We mapped each reading frame to the KCON gene intervals to identify the region containing its start and the regions it extended through. Where gene intervals overlapped, we assigned each shared base to the upstream gene so that it was counted in only one region.

### Target-site duplications

We aligned the terminal LTRs in each retained HML-2 sequence and recorded the immediately adjacent host sequence. We reported reverse-strand calls in insertion orientation. We joined retained boundary measurements to the included manuscript records by exact locus and record identity. For reassigned records, we used the original locus label. We excluded single-junction observations, unavailable calls, and records without an exact match from paired comparisons. The retained caller compared the host suffix at the 5′ junction with the host prefix at the 3′ junction and selected the longest of 6, 5, or 4 bases with no more than two differences. Because that rule can include bases outside a shorter TSD, we evaluated all three lengths before interpreting a difference. We classified a difference as a candidate TSD difference only if the sequences remained unequal at every length. For comparisons longer than the retained sequence, known overlapping bases provided a lower bound on the number of differences. At 8p23.1a, the retained six-base windows differed at two positions, but the five-base candidates matched exactly. We kept these 42 observations separate. The analyzed catalog contained 12,095 paired calls, 8,582 single-junction calls, 38,114 unavailable calls, and 865 records without an exact retained boundary match. Source tables preserve the sequences, comparison lengths, admission states, and record identities.

### Multi-copy resolution and coverage validation

We divided tandem arrays into individual copies using the ordered LTR boundaries. We retained copies on separate contigs or separated by host sequence and recorded their individual coordinates and flanks. Oxford Nanopore reads were aligned with minimap2 using the map-ont preset (4). We used samtools depth (5) to measure median coverage across each HML-2 copy and its adjacent host sequence. For the within-region depth check, we divided median proviral depth by half the median local-flank depth and used a cutoff of 0.6. This check assessed support for a provirus within its assembled region. The additional-copy classification below instead compared local-flank depth with the donor's baseline across regions. Regions with zero flank coverage or <0.4 times the sample median of positive regional flank medians were excluded from depth ratios, but retained in NucFreq summaries. Sites were counted as mixed-base calls when the second-most abundant base had ≥3 reads and represented ≥15% of the top-two base-count sum. Regional fractions divided qualifying sites by positions with nonzero top-two depth. No additional depth cutoff was applied. We assessed NucFreq positions in the interior interval obtained by excluding 10 kb from each end of the depth track. For tracks of 20 kb or less, we used the full span. Regional fractions were screened against 0.05.

We calculated array-spanning read identity from NM and aligned length. At 14q11.2, the three-record state includes two tandem proviruses and a separate fragment. The fragment contributes to total structural copy count but is excluded from the within-array comparisons in Figures 2C–F.

For candidate additional copies, the depth model used the ratio of local-flank median depth to the donor's diploid flank-depth baseline, capped at 1. We compared Gaussian likelihoods centered at 0 for an unsupported assembled copy and 1 for a supported copy. Both states used the same empirical scale—the larger likelihood determined exclusion or retention in the population summaries. The two likelihoods were equal at a ratio of 0.5. Their normalized values remain in Table S2. We retained nine structurally authenticated segmental duplications independently of this depth classification. The 34 depth-classified artifacts and one additional 7p22.1 record crossing an assembly gap were excluded from population, copy-number, and ORF summaries. The records remained in the output table. Seventy-eight records that duplicated the same assembled interval under different locus labels were also retained in the table but counted only once.

The GRCh38 8q24.3b and 8q24.3c flanks map to the same CHM13 interval. Validation compared the elements and 5-kb, 80–100-kb, and 0.5–1-Mb host flanks in HPRC assemblies, and inspected both ends of orphan contigs carrying either label. We retained the 8q24.3b alias rows in the table but excluded them from counts.

**Nucleotide differences within tandem arrays**

We extracted each retained array unit from its source FASTA using its catalog copy interval and strand. We derived copy numbers and copy order from the retained structural catalog. We retained every nucleotide and any insertions absent from KCON. We collapsed exact duplicate sequences for alignment, then restored their original copy and haplotype identities. We aligned each locus with Type-II KCON using MAFFT --auto. KCON supplied coordinates, not an ancestral sequence. For internal comparisons, we used KCON positions 969–8504. This excluded the LTRs shared by adjacent units. We kept insertions within this interval in the alignment.

For each pair of copies in an array, we counted substitutions at positions called A, C, G, or T in both copies. We used the definition of nucleotide divergence from the main Methods. We counted contiguous internal gap differences separately and recorded their lengths and sequences. We did not count terminal differences in sequence extent as indels. These are observed gap tracts, not inferred numbers of independent mutational events. Gap placement within a repeat can depend on the alignment. Array-level diversity was the mean of all pairwise nucleotide divergences within that array. Locus-level means gave each array equal weight. We also retained the nucleotide patterns in copy order and the frequency of each variable site or gap among arrays. We did not treat repeated observations of the same variant across haplotypes as independent mutations.

For arrays with at least three copies, we compared every pair and retained all ties for the smallest nucleotide divergence. Shared nucleotide patterns were evaluated alongside physical copy order. We also calculated conditional divergence times as $t = k/(Lr)$, where $k$ is the substitution count, $L$ is the number of jointly called bases, and $r$ is the pairwise substitution rate. We used $r = 0.0024$–$0.0045$ differences per site per million years from the published LTR clock (1) for rate-sensitivity calculations. Applying this range to internal sequence assumes neutral evolution at those rates. Because $r$ is already a pairwise rate, we did not multiply it by two. We obtained exact Poisson 95% count intervals at each rate. For zero differences, we reported the one-sided 95% upper limit, $-\ln(0.05)/(Lr)$, instead of assigning age zero. We did not pool pairwise comparisons sharing copies as independent observations. These estimates describe the time required to accumulate the observed divergence under the model. They identify a duplication time only if copying began with identical sequence and later exchange did not reset the differences.

**Short-read comparison**

For the ORF-associated variant comparison, we retained the 282 donors and 83 loci shared by the long-read and short-read catalog panels. Targets were derived from the retained long-read sequences, independently of whether the short-read VCF contained a matching record. We bound each native sequence to its donor, haplotype, locus, and source record. Of 20,096 retained provirus and copy records, 19,031 had authenticated native sequences; 1,065 lacked an exact retained sequence binding and were excluded from target discovery. We did not substitute parent tandem-array windows for individual copy records. Four loci lacked a mapped hg38 element interval. Table S3 includes source bindings, exclusions, hashes, and coordinate checks.

We recovered native single-nucleotide and indel genotypes from the 20201028_3202_raw_GT_with_annot high-coverage 1000 Genomes callset for the 79 mapped loci. We verified the genomic coordinates and orientation of each retained hg38 interval against every VCF REF allele contained within it. All 34,100 checked REF alleles matched. We aligned the type-appropriate KCON reference to hg38 to annotate each coding frame, then aligned native long-read sequences to that locus using minimap2 with the asm20 preset. Target discovery required one primary alignment, mapping quality of at least 20, and at least 90% aligned nucleotide identity. We never filled in unaligned reference sequence.

The target set included single-nucleotide changes between a premature stop and a non-stop codon, and indels shorter than 50 bp whose length change relative to hg38 was not divisible by three. Stop-codon targets required all three long-read bases to map contiguously to the canonical codon and exactly one difference from the hg38 codon. Terminal stop codons and codons containing multiple substitutions or gaps were outside this target definition. We normalized indels against the genomic-forward hg38 sequence before annotation. We excluded repeat-equivalent placements that crossed a coding boundary. A frame-changing indel can either disrupt or restore a reading frame, depending on the surrounding sequence. Each gene retained its own canonical reading frame. Type-I Env refers only to the theoretical N-terminally truncated reading frame; translation of this product has not been demonstrated.

We compared normalized alleles as carrier states and did not infer phase or allele dosage. We supported a target when a complete short-read genotype included its alternate allele and met depth ≥10, genotype quality ≥20, and site/sample filter criteria. We supported an explicit all-reference genotype that met the same criteria as a supported reference call. We reported supported alternate alleles separately. Absent records, missing or partial genotypes, inadequate or unavailable quality, failed filters, and ambiguous representations remained unresolved. Equivalent indels were normalized before matching. Conflicting duplicate records or overlapping complex alleles that could not establish the target allele remained unresolved. hg38 bases never replaced missing records or alleles.

The final comparison contained 564 distinct alleles at 36 loci and 20,500 donor–locus–variant pairs across all 282 donors. We counted a variant once per donor and locus even if multiple haplotypes or copies carried it. Gene-specific rows can overlap because the viral genes overlap; overall totals deduplicate these pairs. Recovery uses all tested pairs as its denominator. Concordance among callable pairs uses only supported alternate, supported reference, and supported other-alternate calls. Unresolved calls are excluded from that latter denominator and are reported in full. The data package supplies every target, its long-read evidence, the native short-read genotype and quality fields, the classification, per-locus and per-gene counts, and the executable comparison. The 1q22 example is restricted to 272 long-read carriers of the stop-removing allele; it is not a complete Gag-ORF classification.

To quantify missing evidence for intact coding sequences, we joined each targeted variant to the exact long-read copy and its gene annotation. This analysis required the catalog status Intact and excluded the broader altered-frame category. Of 13,264 intact copy–gene observations, 2,330 carried at least one targeted variant and entered this assessment; the other 10,934 remained untested by this selected variant panel. We recorded whether any target in that copy and gene was unresolved or discordant, or whether all targeted alternate alleles had supported carrier calls. The unresolved fraction was 2,166/2,330 (93.0%). Grouping by donor, locus, and gene gave 1,460/1,565 (93.3%) groups containing at least one unresolved variant in an intact copy. These counts do not imply that other copies were non-intact. Even complete support for the targeted alleles does not establish coverage of every ORF base or phase across the gene. The copy-level joins and separate untested inventory accompany Table S3.

We examined the four 1000 Genomes callsets described in the main Methods (6-8). For the Illumina ensemble comparison in Figure S1, we retained structural-variant records with a direction and reference state compatible with the proposed HML-2 change. We recorded whether the assembly contradicted an alternate call and whether the structural state disagreed with the assembly. We averaged errors within each donor–locus combination before calculating the locus-level fraction. Each panel shows the 15 loci with the highest disagreement among those with at least ten matched donor–locus combinations.

**Phylogenetic and recombination analyses**

Representative proviral sequences were aligned with MAFFT (3). For the neighbor-joining trees (9) in Figures S7A–B and S8A–C, we counted repeated labels for the same assembled interval once.

For the main LTR and Pol overview, we pruned these trees to retain the most frequent exact-sequence cluster at each locus, plus the nearest other-locus clusters used in the 19p12c and 10q24.2 comparisons. Figure S8A and S8C retain both frequency-ranked clusters where available.

We linked each retained KCON alignment record to its original extraction locus, donor, and haplotype, then assigned it to the physical locus established from its source sequence. Copy-suffixed records, multiple-source slots, and telomeric records required a unique full-query alignment to one current source FASTA with minimap2 asm20. These alignments had no mismatches, query insertions, or clipping. We retained source insertions omitted by the KCON projection in the alignment record. We projected source sequences to KCON using the same annotation caller. We lifted Type-I projections to the Type-II reference grid across the verified 292-base deletion. Each source copy contributed once. If a source had multiple projections, we chose the one with the greatest A/C/G/T coverage. The LTR alignment used KCON positions 8504–9471, or positions 0–967 if the 3′ LTR did not contain at least 600 A/C/G/T bases. We defined gene intervals as gag [1111, 3112), pro [2913, 3918), pol [3878, 6749), and env [6450, 8550).

Coordinates were zero-based and half-open. Each region required at least 50% A/C/G/T coverage. We retained up to the two most frequent exact aligned sequence clusters per locus and region. We used the pairwise nucleotide-divergence calculation defined in the main Methods and inferred neighbor-joining trees with Biopython. Negative branch lengths were set to zero. Locus counts include all retained observations meeting the region threshold, not only the two displayed clusters. We pruned the main-figure overview trees from the full trees without refitting their distances. For LTR–Pol comparisons, we restricted candidate neighbors to the 64 loci present in both trees. We compared the modal focal cluster with every retained cluster at other common loci using summed branch lengths along the connecting path. The five-region comparison used the 46 loci shared by all five trees. We selected modal clusters separately in each region. For the 4q35.2 comparison, we counted nucleotide differences at aligned positions where both selected Pol sequences had A, C, G, or T. We excluded gaps and ambiguous bases. We also inspected the retained pairwise minimap2 reference alignments for a continuous alignment across the element and both 5-kb host flanks (Table S9). The chromosome diagram includes this broader relationship separately from exact gene-sequence sharing.

Telomeric assignment required a primary host-linked alignment with MAPQ at least 20, at least 80% of the source element aligned, and at least 50% of its bases within the named reference element. Exactly one candidate reaching at least half of the 80–100-kb chromosome-interior window established a long-flank assignment. If no such unique long anchor was present, assignment required exactly one candidate across the full comparison and alignment of at least half of each 5-kb host flank. Competing long anchors and local chromosome-4 matches did not establish a named telomeric assignment. Source and target had to belong to the same Type-I or Type-II family. We verified exact source subspans against the native assembly or graph sequence before projecting host alignments. Records without a unique named assignment remained in their corresponding copy group.

We reused current source-to-locus assignments and counted each assembled source once. Complete reference windows included 15p13a, 21p13, and 22p13 from CHM13, and 4q35.2 from GRCh38. After orienting all sequences consistently, we aligned the first and last 1,200 bases of 15p13a using local alignment (match 2, mismatch −3, gap opening −10, extension −0.5). Homologous LTR regions occupied zero-based, half-open intervals [6,1000) and [6286,7283) in the 7,283-base reference segment.

We aligned genomic sources using minimap2 asm20, CIGAR output, explicit match/mismatch operations, and no secondary alignments. We required one primary alignment and ≥4,000 mapped reference positions for inclusion. Projected nucleotide sites required A, C, G, or T in ≥90% of sources at every locus. This retained 6,652 positions. Between-locus divergence was the mean probability of a nucleotide difference between random draws from the two loci. Within-locus diversity used the n/(n−1) correction at each site. Shared nucleotide profiles used the 4,797 positions called in all 426 sources. We excluded indels and unobserved positions.

We extracted mapped LTRs with their insertions and aligned them with MAFFT --auto. Each LTR required ≥500 mapped bases and each pair ≥500 jointly called bases. Four reference copies and 26 population copies passed. The population copies comprised 24 at 4q35.2 and one each at 15p13a and 22p13. Figure S2A used 992 positions called in all eight reference LTRs, the fraction of these positions that differed between each pair, neighbor joining, and 1,000 column-bootstrap replicates. Midpoint rooting was for display.

Reference flank alignments used the provirus plus 8 kb on each side. A single primary alignment had to span the provirus and both adjacent 5 kb flanks at mapping quality ≥50. All four references passed. Upstream, proviral, and downstream comparisons used 4,997, 7,275, and 4,992 positions called in all four references. For each of the three unrooted quartet splits, we summed the two within-pair mismatch counts, selected the minimum, and retained ties. We resampled columns 2,000 times. Figures S2B and S2C show neighbor-joining trees oriented from 4q35.2 without asserting a root. All bootstraps used seed 20260918.

We calculated LTR ages as the nucleotide mismatch fraction divided by 0.0024 or 0.0045 per million years. These are pairwise rates from Subramanian et al. (1), and we did not apply an additional factor of two. Indels were not counted as substitutions. Note that the range reflects alternative clock rates, not a confidence interval.

For the nucleotide-sharing network, we linked each donor sequence to its source FASTA and assigned a genomic locus. We counted identical sample, haplotype, and assembled-interval records once. Sequences were aligned to Type-II KCON with minimap2 asm20, --score-N=0, --secondary=no, and CIGAR output. The analyzed intervals were gag 1111–3111, pro 2913–3917, pol 3878–6748, and env 6450–8549, all zero-based inclusive. Both boundary bases had to occur in the same alignment. We retained the intervening sample DNA and preserved its insertions and deletions. We excluded sequences with ambiguous bases and gene regions yielding multiple distinct copy sequences. The locus-level network excludes Type-I telomeric, Type-II telomeric, and unresolved 8p23.1 pools. Each edge counts identical gene-region DNA sequences observed at two distinct assigned loci, deduplicated by gene and nucleotide sequence. A single assembled interval cannot contribute evidence to both endpoints of an edge. We show only loci that share at least one complete gene sequence. We recovered source FASTA files for 6,375 of 6,378 selected records. We excluded the three records without an unambiguous source from this sequence comparison.

For the APD labels in Figure 3B, we compared all available copies at each connected pair of loci, separately for each gene. We required gene regions to have at least 50% A/C/G/T coverage of the KCON interval. We calculated divergence at positions called in both sequences and gave each gene-to-gene comparison between two copies equal weight in the overall mean. Thus, genes represented by more copy pairs contributed more values to that mean. Table S9 also reports each gene separately.

At Xq28a and Xq28b, the sequence-sharing dataset contained env sequences from 214 and 157 copies, respectively. We aligned the 17 distinct env sequences with MAFFT and weighted each by its observed copy count. The alignment contained 2,110 bases and no gaps. The 33,598 between-locus copy comparisons had a mean divergence of 0.0134%.

Within-array sequence comparisons in Figures 2C–F used the gene-specific protein-changing variant profiles and ORF classifications recorded for each copy. A tandem array was variable for a gene if its members had different profiles or ORF classifications. These profile comparisons are distinct from the exact nucleotide-sequence comparison in Figure 3B.

**Chromatin accessibility analysis**

The bonFIRE consensus peaks and Fiber-seq FIRE calls came from lymphoblastoid cell lines from 39 donors with matched genome assemblies (10-12).

For Fiber-seq, we analyzed 21 HML-2 consensus peak files generated with bonFIRE. We used primary sequencing runs (is_primary_sample=true) and retained peaks with total coverage ≥10 reads. We summed FIRE and total coverage across assayed consensus peaks within each primary sample run, donor, and haplotype, then took their ratio. Primary-run selection retained 78 haplotypes from 39 biological donors and excluded 12 alternate run–haplotype records. The dataset comprised 1,344 haplotype-locus observations from 39 donors. Per-locus counts ranged from 26–77 haplotypes and 22–39 donors. We matched assay labels to the structural catalog by CHM13v2.0 coordinates and ordered loci by median accessibility (Figure S17). The crosswalk in Supplementary Data links all 21 assay labels to their coordinates. Twenty intervals matched catalog loci. The separately assayed 14q12b interval is not present in the structural catalog. The 7p22.1 assay covers one proviral segment of the tandem array.

**Type-I deletion analyses**

We use Type-I cassette to mean the shared 292-bp deletion at the pol-env boundary together with its associated flanking sequence. For the post-integration conversion analysis, we used a panel of 58 autosomal loci with at least 59 cassette-typed chromosomes from unrelated donors per locus. Nineteen loci contained only Type-I calls and 39 contained only Type-II calls. We excluded offspring, duplicate assemblies, and reference genomes from this sample. We modeled a fixed occupied recipient population initially carrying Type II. Type I could arise at any time, and a donor was always available. Converted alleles could drift to loss or fixation, and conversion could recur during polymorphism or after loss. For Type-I frequency x, the mean change per generation was $\mu(1-x)$ and the variance was $x(1-x)/(2N_e)$. The model assumed no selection. We used 25 years per generation and effective population sizes ($N_e$) of 5,000, 10,000, and 20,000. This model conditions on occupied recipient populations.

We used the low, mean, and high interlocus conversion rates reported by Dumont and Eichler, $6.1 \times 10^{-7}$, $3.2 \times 10^{-6}$, and $1.05 \times 10^{-5}$ per duplicated site per generation (13). We also tested the independent estimate of $2.5 \times 10^{-7}$ and its bootstrap 95% interval, $0.8 \times 10^{-7}$ to $5 \times 10^{-7}$, from Harpak et al. (14). We used these estimates from human paralogs as effective cassette-transfer rates, without multiplying by deletion length or donor count.

The six youngest age-linked Type-I loci, 12q13.2, 1p31.1b, 1q22, 3q13.2, 3q27.2 and 5q33.3, were each allowed 2 million years. We assigned older loci the upper bounds from the published Subramanian et al. age estimates, matched by genomic coordinates (1). Sensitivity analyses doubled each older bound or granted complete Type-I fixation to every older locus. We granted complete Type-I fixation to the three loci without matched age estimates, 19p12c, 1p31.1a, and 1q32.2, in every scenario. These assignments give conversion the full specified time and make the joint probabilities upper bounds within this model. Table S13 includes the locus identities, age bounds, and sample sizes.

We calculated sample probabilities from the neutral diffusion moment equations by matrix exponentiation. For n typed chromosomes, $a=E[x^n]$ is the probability of sampling only Type I, $b=E[(1-x)^n]$ is the probability of sampling only Type II. Conditional on detecting Type I at a particular locus, the probability of no sampled Type-II allele is $a/(1-b)$. We assumed independent locus trajectories given the rates, ages, and $N_e$ and multiplied these conditional probabilities over the observed Type-I-positive loci. The conditioning retained the exact positive locus identities, not just their number. Expected mixed-locus counts were the sum of one minus these conditional probabilities. Chromosomes sampled at one locus shared its drift trajectory. Table S13 provides the full 54-scenario results and an independent finite-volume numerical check.

We grouped sequences missing the same viral interval into one deletion class, defined by its breakpoint coordinates and length. We compared Δ292 with six other deletion classes of at least 50 bp detected in orangutan, siamang or macaque HML-2 sequences. The comparison deletions were 74, 112, 116, 208, 868, and 2,255 bp long; each was observed at one ancestral proviral integration site. In the nonhuman-primate panel, comparisons of the host DNA on both sides of each provirus identified 16 ancestral integrations carrying Δ292 and one possible additional integration. We therefore used count vectors [16,1,1,1,1,1,1] and [17,1,1,1,1,1,1], conditioned on totals of 22 and 23 deletion-class observations. An insertion inherited from a shared ancestral integration counted once across species. The 74-bp and 2,255-bp deletions occurred in Δ292-bearing proviruses, so some insertions contributed to more than one deletion class. Input records and orthology assignments accompany Table S11. A symmetric Dirichlet prior represented relative cumulative contributions of the seven deletion classes. Each weight summarizes the sampled history and does not identify an individual source locus or its active period. We tested $\alpha = 0.1, 0.25, 0.5, 1, 2, 5$, and 20. The plotted coefficient of variation, $1/\sqrt{\alpha}$, describes the unnormalized gamma weights for the deletion classes. Both models allowed the same deletion-class variation. The additional-effect model applied a log-uniform multiplier from 1 to 50 to the Δ292 class. Both models integrated over relative detection odds of 0.67–1.5 or 0.25–4. We used deterministic Gauss–Legendre quadrature with 192 probability nodes and 80 prior nodes to integrate the multinomial likelihood. We used ratios of marginal evidence to estimate Bayes factors.

**Type-I sequence comparisons**

We compared six backbone windows and the concatenated deletion flanks in the retained MAFFT alignment of 45 human locus representatives. Using zero-based, half-open Type-II KCON coordinates, B1–B5 were 1000–2000, 2000–3000, 3000–4000, 4000–5000 and 5000–6000. B6 was 7293–8293. The fixed comparison window joined 6000–6501 and 6793–7293 for a total of 1,001 bases. It excluded Δ292 at [6501, 6793). We used the nucleotide-divergence definition in the main Methods and averaged the proportions over eligible locus pairs. Each sequence required A/C/G/T calls at 90% of a window, and each pair required joint calls at 80%. Every retained pair met the latter threshold. The fixed window excluded the Type-II representative at 10q24.2, which had only 745 called positions. We recorded nearest-neighbor changes relative to the rest of the provirus and performed leave-one-out checks. We grouped other naturally occurring internal deletions by breakpoint and length. Type-I calls required a single junction directly joining callable KCON coordinates 6500 and 6793, with no mapped bases within the deleted interval [6501, 6793) in zero-based coordinates. Type-II calls required at least 278 of the 292 interval positions to contain callable A, C, G, or T bases. We also required at least 15 callable positions in each of three 20-base windows at the beginning, midpoint, and end of the interval. All informative alignment passes within a record had to agree. We joined the retained direct calls to the current catalog by locus, donor, haplotype, cohort, and contig, and counted each donor-haplotype-locus combination once.

The representative-selection procedure prioritized HPRC records. It sorted identifiers lexically and selected the first readable provirus FASTA. It contained 45 of the 64 eligible catalog loci. Eighteen excluded loci lacked a population provirus row, and the historical 8p22 comparator lacked a readable source FASTA. The retained panel contained 15 Type-I and 30 Type-II representatives. The latter comprised 15 LTR5Hs, 11 LTR5A and four LTR5B loci. Table S11 records the selected identifiers, source hashes, and subfamily assignments.

We ranked sampled representatives by their mean cassette divergence from the other Type-I representatives. Site-frequency logos used all available A/C/G/T calls at each position, so their per-position denominator can differ from the whole-window denominator. Letter height represents nucleotide frequency. Table S11 includes base, gap, and ambiguous-call counts. Figure S13 uses one-based inclusive display coordinates. Its linked sites 6332 and 6493 correspond to zero-based positions 6331 and 6492. The Δ292 interval is 6502–6793 in one-based coordinates.

The nonhuman primate panel contains seven bonobo, nine chimpanzee, 19 gorilla, 12 macaque, nine orangutan, and 27 siamang clusters. It includes 26 canonical deletion clusters, 55 retained Type-II clusters, and two alternative-deletion clusters. Cross-species orthologs are separate sequence observations in this alignment. The full cassette projection spans zero-based KCON [6000, 7293) and includes Δ292. We omit insertions relative to KCON and retain projected gaps. One gorilla cluster classified as Type I has a one-column boundary shift in the projection. The aligned FASTA files in Table S11 supporting data preserve that observed alignment.

For the uncertainty columns in Table S11, we resampled callable loci with replacement 2,000 times and recalculated the mean pairwise divergence. Repeated draws of the same locus contributed zero self-distance. The table gives percentile intervals. The main figure shows observed means.

#### Functional association analyses

We screened cellular growth and EBV DNA abundance using superpopulation-adjusted linear models and pedigree-clustered standard errors. Sensitivity analyses used population fixed effects. For the 1q22 Gag comparisons in Figure S16B, the exposure was presence of at least one copy passing the combined ORF screen. The 7p22.1 comparison used diploid copy number. These three comparisons used HC3 heteroskedasticity-robust standard errors. Mandage et al. (15) and Houldcroft et al. (16) EBV outcomes were log2-transformed. Im et al. (17) divided intrinsic growth by 10,000. Burden models used the same HML-2 inclusion criteria as the ORF analysis. The screen contained 237 exposure–outcome combinations. Benjamini–Hochberg correction was applied to the 174 nonmissing P values from 72 Mandage, 54 Houldcroft, and 48 Im tests. The other 63 models had fewer than five donors in one comparison group (44 models), fewer than 30 donors (18 models), or an invariant exposure (one model). Their P and q values remained missing. We retained identical exposure vectors as separate rows when they represented different exposure–outcome tests. The Type-I/II burden analysis comprised 17 exposures across three outcomes, or 51 fits. Results are supplied in Supplementary Data.

For 1p31.1b, we compared internal-fragment presence with SLC44A5 expression in MAGE (18) and GEUVADIS (19). The 39-donor MAGE model included sex, genotype principal components 1–5, and probabilistic estimation of expression residuals (PEER) factors 1–5. We normalized expression by the trimmed mean of M-values (TMM) and inverse-normal transformed it. The donor-disjoint GEUVADIS subset included 28 donors after removing five donors overlapping with MAGE. It used PEER-residualized expression with population and sex covariates. We scaled both displayed coefficients and their standard errors by SD(exposure)/SD(expression). We used HC3 robust standard errors. The MAGE discovery screen considered 1,409 candidate exposures. Complete-case filtering required at least 16 donors, at least three donors per binary state or three distinct values for a nonbinary exposure, a full-rank design, and positive residual degrees of freedom. This retained 146 candidate aliases. We counted identical donor sets, exposure vectors, and nonconstant adjustment vectors once. We tested the resulting 64 exposure models against 20,154 genes. Benjamini–Hochberg correction covered all 1,289,856 ordinary least-squares P values. The separately reported HC3 P value for the MAGE SLC44A5 sensitivity analysis was not included in that correction. GEUVADIS follow-up correction included 30 estimable population-adjusted tests. The prespecified replication and superpopulation-adjusted sensitivity models were not part of this secondary family. Tables S15e–h enumerate the discovery family, candidate eligibility and aliases, all 98 GEUVADIS follow-up rows, and the MAGE HC3 sensitivity models.

For 12q13.2, we used the anti-CD20 viability data of Small et al. (20). The 49-donor serum cohort contained 33 provirus carriers, 11 solo-LTR carriers without a provirus, and five noncarriers. A three-class structural model estimated the provirus-versus-solo-LTR contrast while modeling the noncarrier group separately. Models included sex and population fixed effects and used HC3 standard errors. The media outcomes had 48 and 45 evaluable donors. Sensitivity analyses used 20,000 Freedman–Lane residual permutations within population. We applied a Benjamini–Hochberg correction (21) across 40 adjusted state/outcome models. All anti-CD20 conditions came from the same cohort. The displayed intervals use the normal approximation, calculated as the coefficient plus or minus 1.96 times its robust standard error.

**Sequence and archaic evidence at 8q11.23**

At 8q11.23, structural counts used all 584 donor haplotypes. Sequence analysis included all 583 solo-LTR observations. We retrieved 461 sequences from the assembly extraction directory and 122 from the graph extraction directory using their exact retained catalog identifiers. We aligned both sets to the 968-bp LTR reference and recovered all 968 positions in every sequence. The 13 segregating sites were within the LTR. The full provirus came from the paternal haplotype of HG03098. Population groups followed IGSR, HPRC, HGSVC, and Coriell metadata (Table S4).

We examined published GRCh37 alignments from the Vindija 33.19, Altai, and Chagyrskaya Neanderthals and the Denisovan individual (22-25). We verified the LTR boundaries by sequence identity with the corresponding GRCh38 and T2T-CHM13 sequences and their flanking target-site duplications. We retrieved regional alignments with samtools 1.23.1. Junction support required mapping quality ≥25 and an uninterrupted aligned block extending at least 10 bp into both the host flank and LTR. We excluded unmapped, secondary, supplementary, duplicate, and QC-failed alignments. We counted reads separately at each junction and repeated the checks using 20-bp and 30-bp anchors. We verified ape empty sites by aligning both human host flanks and checking for a single shared target-site sequence without an intervening LTR (Table S14).

**Solo-LTR diversity and expected differences at 8q11.23**

We used the retained catalog and the retained solo-LTR sequences from the same public donors used in Figure 6. Reference genomes were excluded. Each donor-haplotype sequence contributed once at a locus. Available extractions could include host sequence and differ in total length, so we mapped each unique sequence to the 968-bp HML-2 LTR reference with minimap2 asm20 and retained primary alignments. We required at least 800 aligned A/C/G/T bases per sequence and at least 800 positions callable in every retained sequence at a locus. Thirteen loci met these criteria and had at least 20 sequences each. They contained 938–968 common callable LTR positions. The final counts ranged from 22 to 583 sequences per locus. Eleven other loci had fewer than 20 available sequences. The 15q25.2 extractions did not yield qualifying LTR alignments and were excluded from this comparison. These exclusions are listed with the source data.

We calculated nucleotide diversity as the total number of differing A/C/G/T pairs divided by the total number of comparable base pairs. Gaps, insertions relative to the reference, and ambiguous bases were excluded. At 8q11.23, the 583 sequences supplied 169,653 pairs of sampled sequences and 164,224,104 comparable base pairs across 968 positions. The 20,822 nucleotide differences gave 0.122733 differences per sequence pair.

We compared the observed solo-LTR differences with the clock expectation E[D] = Lrt across divergence times, using L = 968 bases and pairwise rates r = 0.0024 and 0.0045 differences per site per million years (1). No additional factor of two was applied. For two solo-LTR alleles, t represents their pairwise coalescence time. For the two LTRs of a provirus, t represents time since insertion only if subsequent exchange has not reset their differences. The comparison shows how little sequence divergence is present at 8q11.23; it does not assign an age to its occupied insertion site.

### Supplemental Results

#### Validation of structural calls

We identified one redundant named locus. The GRCh38 coordinates assigned to HML-2 8q24.3b and its extended flanks coincided with the T2T 8q24.3c interval in 465 of 466 assemblies. The additional matching contig lacked distinct flanks. We therefore treated 8q24.3b as an alias of 8q24.3c, not a separate locus. Similar proviruses in duplicated host regions complicate assignment to individual telomeric loci. Flanking host sequence assigned 2,235 records to named chromosome arms. We retained 603 records without a unique supported assignment in Type-I or Type-II copy groups (Table S5). Of 45 suspected additional-copy records, 34 had local-flank median depth at or below 50% of the sample-wide diploid flank-depth baseline, and one HG00658 7p22.1 record crossed an assembly gap. We retained ten copies. Nine were independently supported segmental duplications (Figure S3A–B, Table S2). We also identified 78 records that represented an assembled interval already listed under another locus name and counted those intervals once. The largest fraction of mixed-base sites among 351 evaluated regions was 0.0135, below the 0.05 screening threshold (Table S2 supporting data). The extra paternal 1q22 record in HG00423 illustrates an unsupported assembly copy. Its proviral and flanking read depths were lower than those of the supported copy. We excluded the extra record and retained the supported copy (Figure S3C).

#### Target-site boundary comparisons

Among 12,095 retained paired boundary observations, 20 pairs at five loci differed at every candidate length from 4 to 6 bp (Figure S6A). At 8p23.1a, all 42 six-base comparisons were ACCTTT/CCTTTT, but the adjacent five-base sequences were identical: CCTTT/CCTTT (Figure S6B). We did not count those longer-window differences as TSD substitutions. Only one terminal junction was supported at 3q12.3, 3q21.2, 4q32.3, and 5p12, so those loci did not contribute paired comparisons.

#### Regional phylogenetic comparisons

For the most frequent 19p12c sequence, the nearest other-locus sequence by tree distance was 12q14.1 in the LTR tree and 22q11.21 in Pol. The nearest neighbor of 10q24.2 changed from 12q14.1 to 6q14.1 (Figure 3A). Both comparisons were restricted to the 64 loci represented in the LTR and Pol trees. These particular neighbors were recovered in 24.0% and 47.9% of bootstrap replicates for 19p12c and 18.4% and 51.3% for 10q24.2. These results provide limited support for the specific nearest-neighbor relationships. Gag and Pro gave two other nearest neighbors for 19p12c, 8p23.1a and 7p22.1 (Figures S7A–B). The expanded LTR tree retains up to two sequence clusters per locus (Figure S8A). The nearest Env neighbor was 10p12.1 (Figure S8B). The expanded Pol tree also retains up to two clusters per locus (Figure S8C). The Gag, Pro, and Env relationships held among the 46 loci represented in all five trees.

**Alternative reading frames**

The independent single-frame scan examined possible products distinct from canonical polyproteins. Of 5,298 copies whose longest ATG-starting ORF began in gag, 4,824 overlapped gag alone, 263 overlapped gag and pro, and 211 overlapped all three annotated regions (Figure S10B). The last group consisted of 339-amino-acid ORFs at 3p12.3. Their overlap with all three regions does not imply a full-length Gag–Pro–Pol product.

**Additional functional comparisons**

Combined-screen Gag copy number at 1q22 was zero, one, or two in 124, 113, and 55 donors, respectively (Figure S16A, Table S16). Carriage of at least one 1q22 Gag copy passing the combined screen was not associated with growth or EBV DNA abundance, nor was 7p22.1 copy number associated with EBV DNA abundance (P=0.61, 0.41 and 0.92, respectively, Figure S16B).

Type-I copy number with an np9 annotation passing the combined screen was associated with EBV DNA abundance after adjustment for total Type-I copy number ($P=8.76 \times 10^{-4}$). The q value was 0.0447 across the 51 burden models and 0.0508 across the 174 estimable tests in the 237-model screen.

**Sequence differences and relative history of tandem arrays**

We recovered 617 unit sequences from 289 of the 290 retained arrays across twelve loci. These included all 247 arrays at 7p22.1 and all nine at 1p31.1b. Source sequence was unavailable for the two-copy NA20282 h2 array at 14q11.2, so we excluded it from nucleotide comparisons. Table S8 gives the sequences, copy identities, pairwise denominators, substitutions, gap tracts, and site frequencies.

At 7p22.1, 197/247 arrays (79.8%) had internal substitution differences (Figure S5A). Internal gap differences occurred in 167/247 arrays (67.6%). All 167 arrays with gap differences differed at the one-base C tract at KCON position 3154 in pro. The mean within-array nucleotide divergence was 0.0949% (Table S8). Fifty arrays were identical internally at their jointly called positions. Other substitutions were distributed across the internal region. The most frequent substitution difference was at KCON position 3915, observed in 175/247 arrays (70.9%, Figure S5B). The per-array patterns show that much of this variation reflects recurring copy classes across haplotypes.

In three-copy 7p22.1 arrays, copies 2 and 3 differed by 0–3 substitutions, compared with 8–13 differences from copy 1. Three of the four four-copy arrays also separated copy 1 from a more similar downstream group. The fourth, HG01457 h2, had four identical internal sequences. In the six-copy HG04115 paternal array, copies 2–6 were identical across 7,536 called internal bases. Each differed from copy 1 at ten of 7,535 jointly called bases and at a one-base C gap (Figure S5C).

At 1p31.1b, the mean within-array nucleotide divergence was 0.00584%. Five arrays had no internal substitution or gap differences. HG01167 h2 and HG03009 h1 each had the same three-copy nucleotide pattern at KCON position 2298 (T/T/G). HG03470 h1 had a C/T difference at position 8300. HG03732 h2 had a two-base gap at positions 2360–2361 in copy 1. No other internal differences between copies were detected in these nine arrays (Figure S5D). The three four-copy arrays had identical internal sequences within each array.

#### Conditional timing from substitution differences

For the median 7p22.1 pairwise count of nine substitutions across 7,535 bases, the two rate endpoints gave 0.27 and 0.50 million years. This rate sensitivity range is not a confidence interval. At the faster rate, the exact Poisson 95% interval was 0.12–0.50 million years, and at the slower rate it was 0.23–0.94 million years. In the six-copy array, the ten differences between copy 1 and the downstream copies gave 0.29–0.55 million years. The zero differences among downstream copies gave one-sided 95% upper limits of 0.088 million years at the faster rate and 0.166 million years at the slower rate.

At 1p31.1b, one difference across 4,437 bases gave 0.050–0.094 million years. The corresponding exact 95% intervals were broad, 0.0013–0.28 million years at the faster rate and 0.0024–0.52 million years at the slower rate. A pair with no differences across the same length had a one-sided 95% upper limit of 0.150 or 0.281 million years at the two rate endpoints. The low counts are therefore compatible with recent sequence copying, but do not precisely date the duplication events. Gene conversion after duplication could also reduce divergence between copies.

#### Type-I cassette comparisons within LTR5Hs

Across the six backbone windows, mean divergences among LTR5Hs Type-II representatives were 5.78%, 4.26%, 4.64%, 3.53%, 3.18% and 3.97%. Type-I values were 5.47%, 5.24%, 6.44%, 4.24%, 2.97% and 3.47%. Restriction to LTR5Hs therefore removes a general separation between the two types across these windows. Cassette divergence remains lower in Type I, at 2.73% versus 3.68% (Table S11).

The nearest Type-II cassette representative was 8p23.1a for twelve of fifteen Type-I representatives. One more was equally close to 8p23.1a and 11q22.1. The other two were closest to 19q12 and 11q22.1. These assignments did not change after restriction to LTR5Hs. Mean divergence between 8p23.1a and the Type-I panel was 2.134%. Among Type-I representatives, 19p12c had the lowest mean cassette divergence from the others (1.843%); the next three values were 1.865–1.872% (Figure S13B).

The full nonhuman primate alignment reproduces the linked T/T pair at zero-based KCON 6331 and 6492 in 25 of 26 canonical deletion clusters and four of 57 clusters without Δ292. The latter group includes two clusters with other deletions. These are cluster counts, not independent insertion counts.

#### Supplementary References


1. Subramanian, R.P., Wildschutte, J.H., Russo, C. *et al.* (2011) Identification, characterization, and comparative genomic distribution of the HERV-K (HML-2) group of human endogenous retroviruses. *Retrovirology*, **8**, 90. https://doi.org/10.1186/1742-4690-8-90
2. Lee, Y.N. and Bieniasz, P.D. (2007) Reconstitution of an infectious human endogenous retrovirus. *PLoS Pathog*, **3**, e10. https://doi.org/10.1371/journal.ppat.0030010
3. Katoh, K. and Standley, D.M. (2013) MAFFT multiple sequence alignment software version 7: improvements in performance and usability. *Mol Biol Evol*, **30**, 772–780. https://doi.org/10.1093/molbev/mst010
4. Li, H. (2018) Minimap2: pairwise alignment for nucleotide sequences. *Bioinformatics*, **34**, 3094–3100. https://doi.org/10.1093/bioinformatics/bty191
5. Danecek, P., Bonfield, J.K., Liddle, J. *et al.* (2021) Twelve years of SAMtools and BCFtools. *Gigascience*, **10**, giab008. https://doi.org/10.1093/gigascience/giab008

6. Byrska-Bishop, M., Evani, U.S., Zhao, X. *et al.* (2022) High-coverage whole-genome sequencing of the expanded 1000 Genomes Project cohort including 602 trios. *Cell*, **185**, 3426–3440.e19. https://doi.org/10.1016/j.cell.2022.08.004
7. Ebert, P., Audano, P.A., Zhu, Q. *et al.* (2021) Haplotype-resolved diverse human genomes and integrated analysis of structural variation. *Science*, **372**, eabf7117. https://doi.org/10.1126/science.abf7117
8. Ebler, J., Ebert, P., Clarke, W.E. *et al.* (2022) Pangenome-based genome inference allows efficient and accurate genotyping across a wide spectrum of variant classes. *Nat Genet*, **54**, 518–525. https://doi.org/10.1038/s41588-022-01043-w
9. Saitou, N. and Nei, M. (1987) The neighbor-joining method: a new method for reconstructing phylogenetic trees. *Mol Biol Evol*, **4**, 406–425. https://doi.org/10.1093/oxfordjournals.molbev.a040454
10. Stergachis, A.B., Debo, B.M., Haugen, E. *et al.* (2020) Single-molecule regulatory architectures captured by chromatin fiber sequencing. *Science*, **368**, 1449–1454. https://doi.org/10.1126/science.aaz1646
11. Vollger, M.R., Swanson, E.G., Neph, S.J. *et al.* (2026) Somatic epimutations cap genetic determinism in the human diploid chromatin epigenome. *bioRxiv [preprint]*, 2024.06.14.599122v3. https://doi.org/10.1101/2024.06.14.599122
12. Lucas, J.K., Hebbar, P., Liao, W.-W. *et al.* (2026) HPRC2: A human pangenome reference with near-complete coverage of common genetic variation. *bioRxiv [preprint]*, 2026.07.21.739710v1. https://doi.org/10.64898/2026.07.21.739710
13. Dumont, B.L. and Eichler, E.E. (2013) Signals of Historical Interlocus Gene Conversion in Human Segmental Duplications. *PLoS One*, **8**, e75949. https://doi.org/10.1371/journal.pone.0075949
14. Harpak, A., Lan, X., Gao, Z. *et al.* (2017) Frequent nonallelic gene conversion on the human lineage and its effect on the divergence of gene duplicates. *Proc Natl Acad Sci U S A*, **114**, 12779–12784. https://doi.org/10.1073/pnas.1708151114
15. Mandage, R., Telford, M., Rodríguez, J.A. *et al.* (2017) Genetic factors affecting EBV copy number in lymphoblastoid cell lines derived from the 1000 Genome Project samples. *PLoS One*, **12**, e0179446. https://doi.org/10.1371/journal.pone.0179446
16. Houldcroft, C.J., Petrova, V., Liu, J.Z. *et al.* (2014) Host genetic variants and gene expression patterns associated with Epstein-Barr virus copy number in lymphoblastoid cell lines. *PLoS One*, **9**, e108384. https://doi.org/10.1371/journal.pone.0108384
17. Im, H.K., Gamazon, E.R., Stark, A.L. *et al.* (2012) Mixed effects modeling of proliferation rates in cell-based models: consequence for pharmacogenomics and cancer. *PLoS Genet*, **8**, e1002525. https://doi.org/10.1371/journal.pgen.1002525
18. Taylor, D.J., Chhetri, S.B., Tassia, M.G. *et al.* (2024) Sources of gene expression variation in a globally diverse human cohort. *Nature*, **632**, 122–130. https://doi.org/10.1038/s41586-024-07708-2
19. Lappalainen, T., Sammeth, M., Friedländer, M.R. *et al.* (2013) Transcriptome and genome sequencing uncovers functional variation in humans. *Nature*, **501**, 506–511. https://doi.org/10.1038/nature12531
20. Small, G.W., Akhtari, F.S., Green, A.J. *et al.* (2023) Pharmacogenomic Analyses Implicate B Cell Developmental Status and MKL1 as Determinants of Sensitivity toward Anti-CD20 Monoclonal Antibody Therapy. *Cells*, **12**, 1574. https://doi.org/10.3390/cells12121574
21. Benjamini, Y. and Hochberg, Y. (1995) Controlling the False Discovery Rate: A Practical and Powerful Approach to Multiple Testing. *Journal of the Royal Statistical Society. Series B (Methodological)*, **57**, 289–300. https://doi.org/10.1111/j.2517-6161.1995.tb02031.x
22. Prufer, K., de Filippo, C., Grote, S. *et al.* (2017) A high-coverage Neandertal genome from Vindija Cave in Croatia. *Science*, **358**, 655–658. https://doi.org/10.1126/science.aao1887
23. Prufer, K., Racimo, F., Patterson, N. *et al.* (2014) The complete genome sequence of a Neanderthal from the Altai Mountains. *Nature*, **505**, 43–9. https://doi.org/10.1038/nature12886
24. Mafessoni, F., Grote, S., de Filippo, C. *et al.* (2020) A high-coverage Neandertal genome from Chagyrskaya Cave. *Proc Natl Acad Sci U S A*, **117**, 15132–15136. https://doi.org/10.1073/pnas.2004944117

25. Meyer, M., Kircher, M., Gansauge, M.-T. *et al.* (2012) A high-coverage genome sequence from an archaic Denisovan individual. *Science*, **338**, 222–6. https://doi.org/10.1126/science.1224344

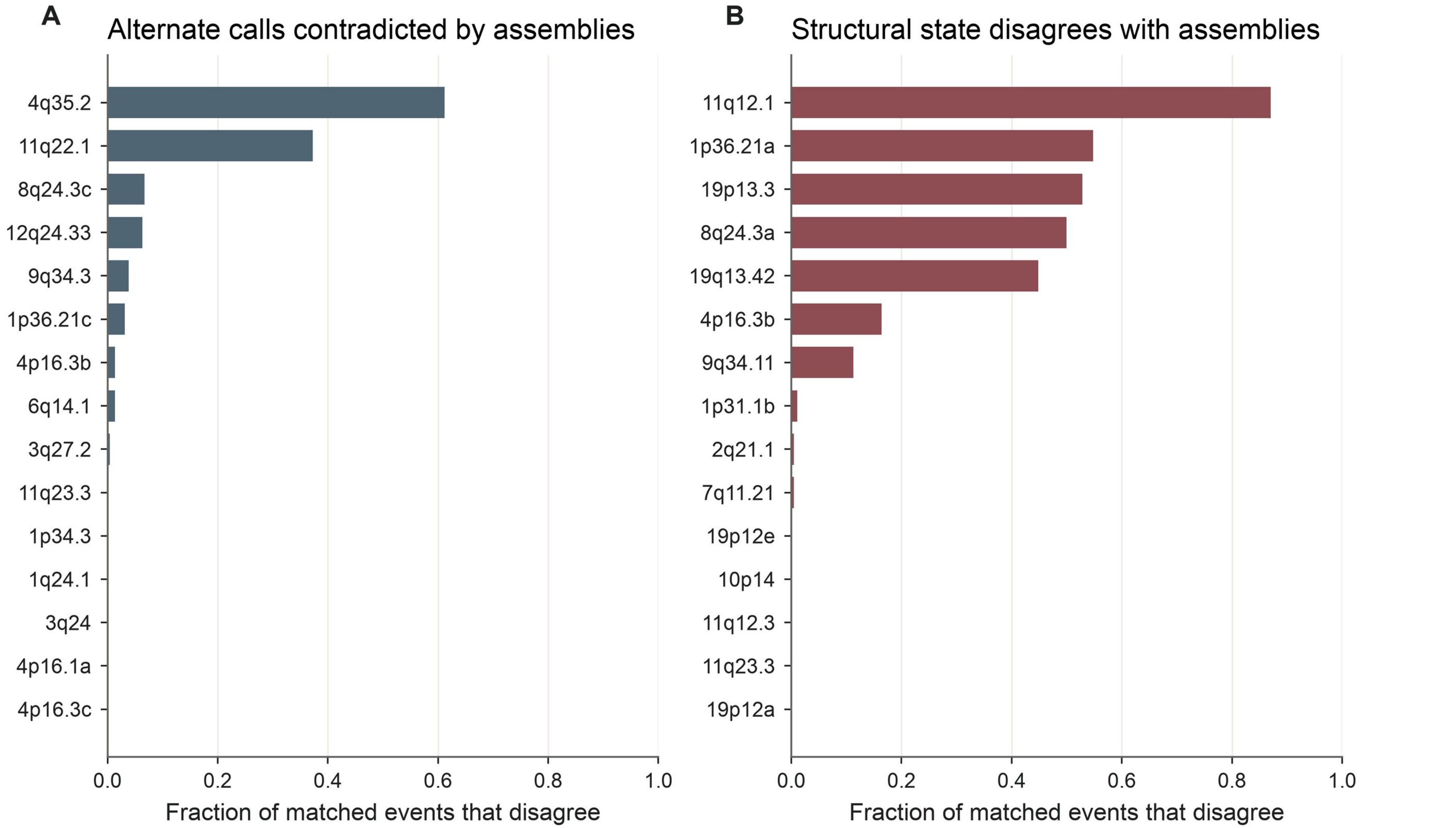


**Figure S1:** Comparison of Illumina ensemble structural-variant calls with matched long-read assemblies. **(A)** Alternate calls contradicted by the assembly. **(B)** Structural states that disagree with the assembly. We averaged records within each donor–locus combination. Each panel shows the 15 loci with the highest disagreement among those with at least ten matched combinations.

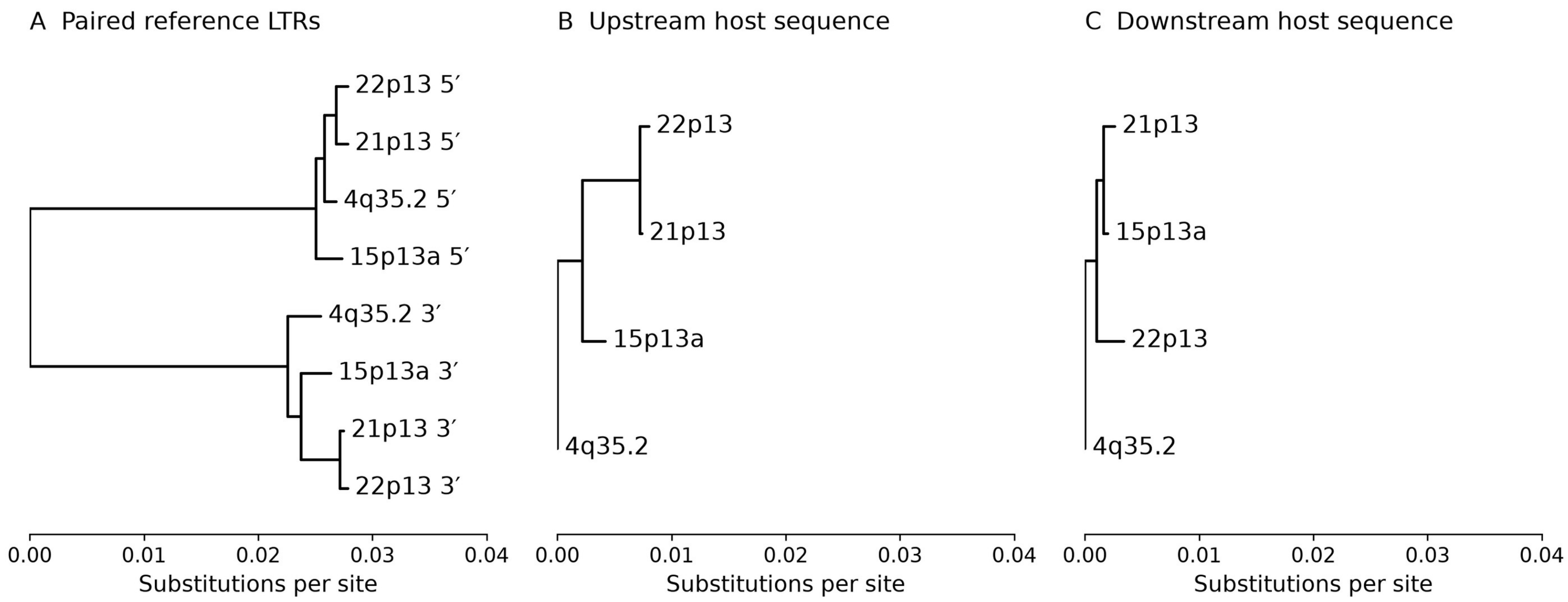


**Figure S2:** LTR and host-flank relationships in the telomeric Type-II family. **(A)** Paired reference LTRs from GRCh38 (4q35.2) and CHM13 (15p13a, 21p13 and 22p13). **(B)** Upstream 5 kb. **(C)** Downstream 5 kb. All panels share the same scale in nucleotide substitutions per site.

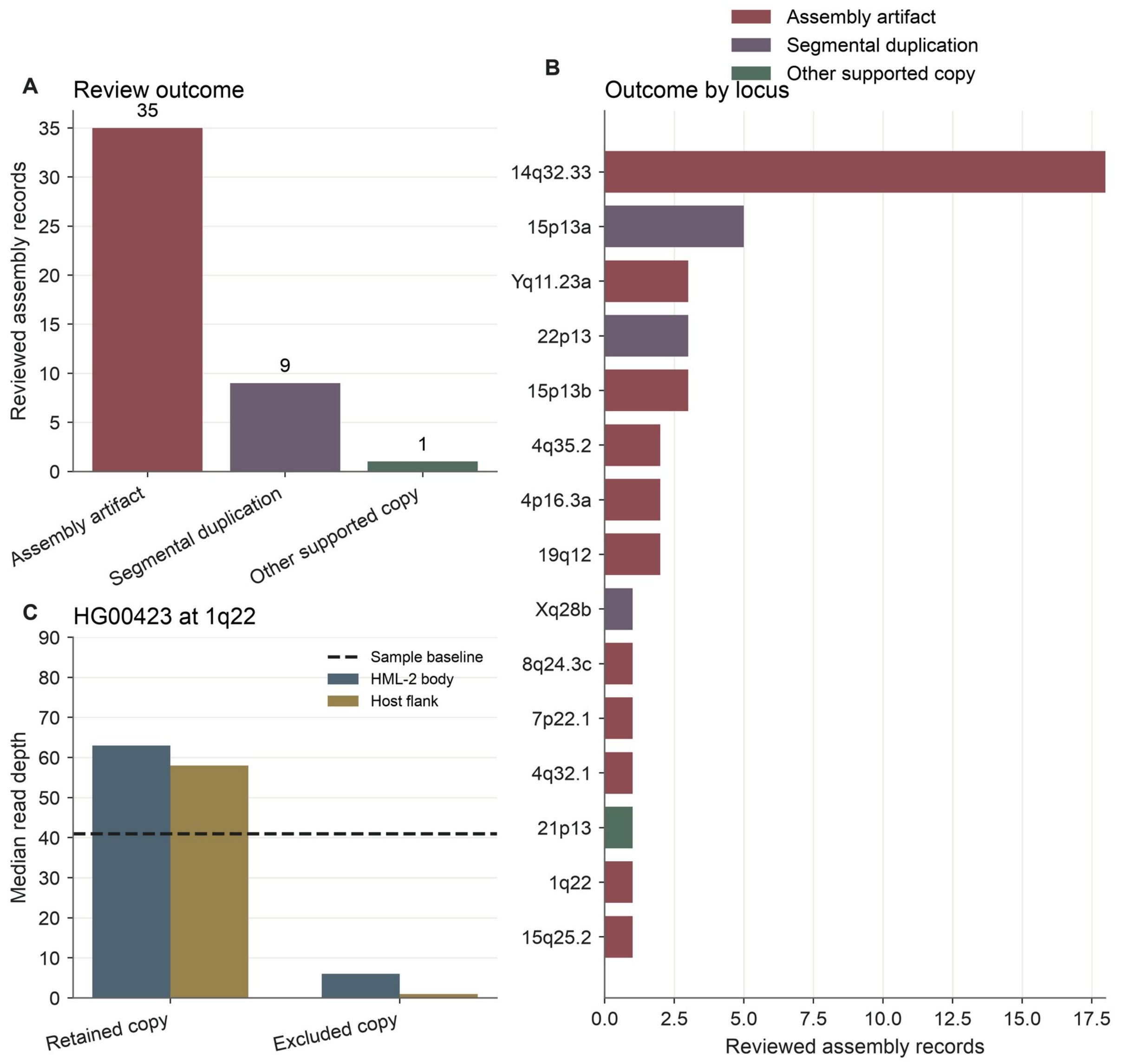


**Figure S3:** Read-depth and assembly review of suspected additional HML-2 copies. **(A)** Classification of 45 candidate records. **(B)** Review outcomes by locus. **(C)** Body and flanking read depth for supported and unsupported paternal 1q22 records from HG00423. The diploid baseline is shown separately.

**Figure S4A Coding status by position within tandem arrays**

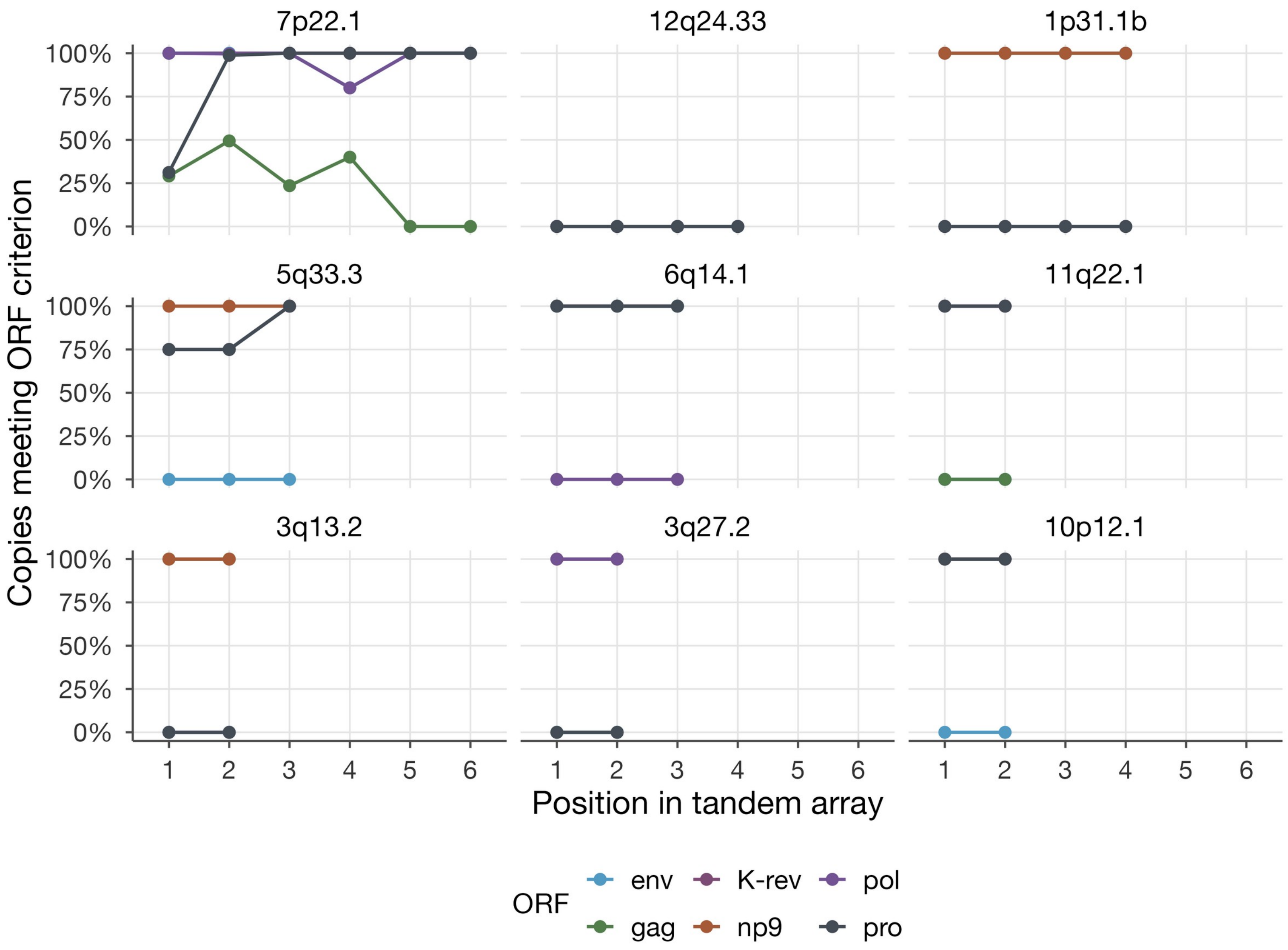

**Figure S4B Tandem-array sizes**

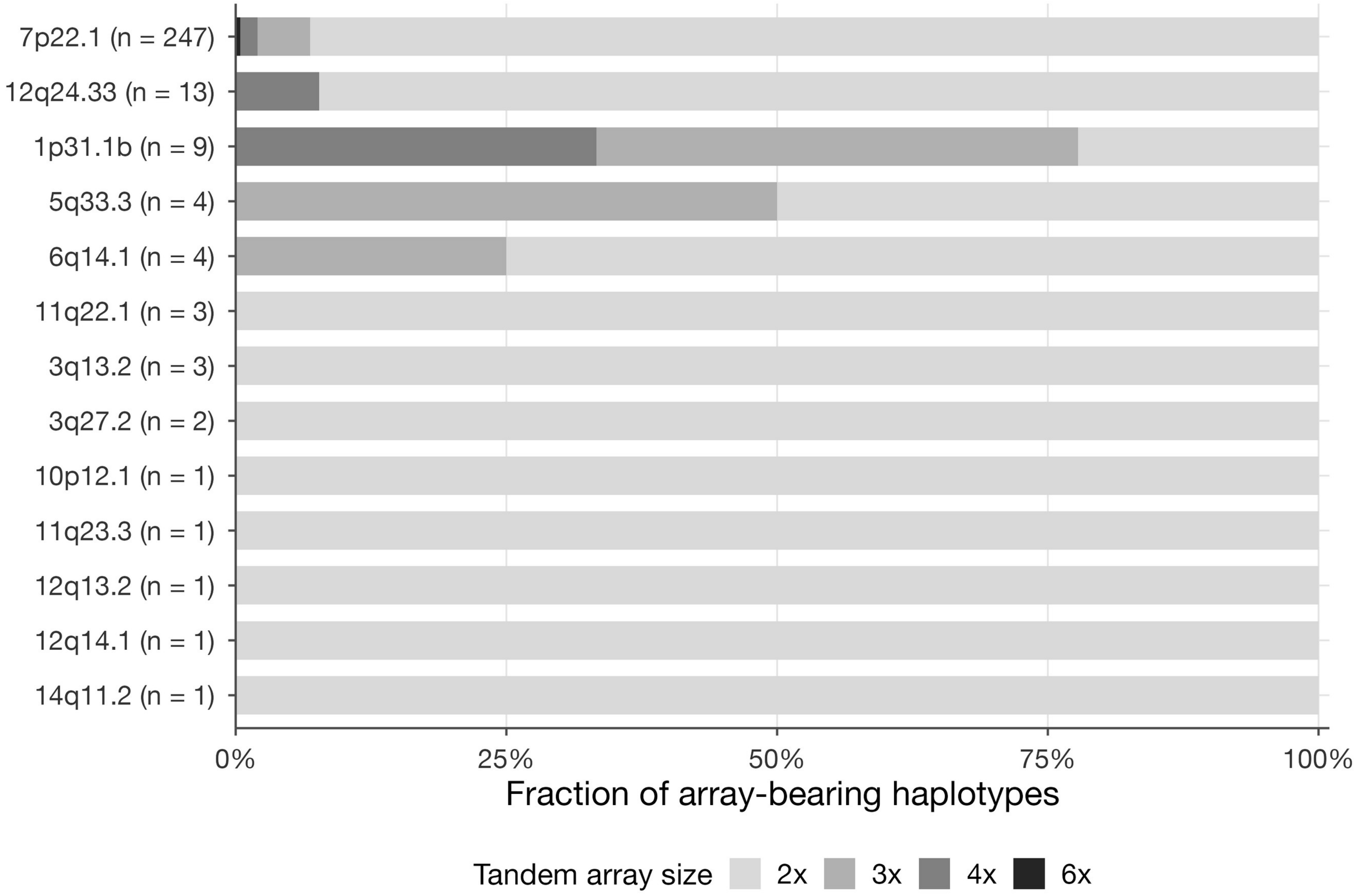


Figure S4: ORF retention and array sizes at tandem HML-2 loci. (A) Fraction of proviral copies passing the combined ORF screen at each array position in the nine most frequent tandem-array loci. (B) Array-size distributions. Labels give array-bearing haplotype counts. K-rev denotes Rec.

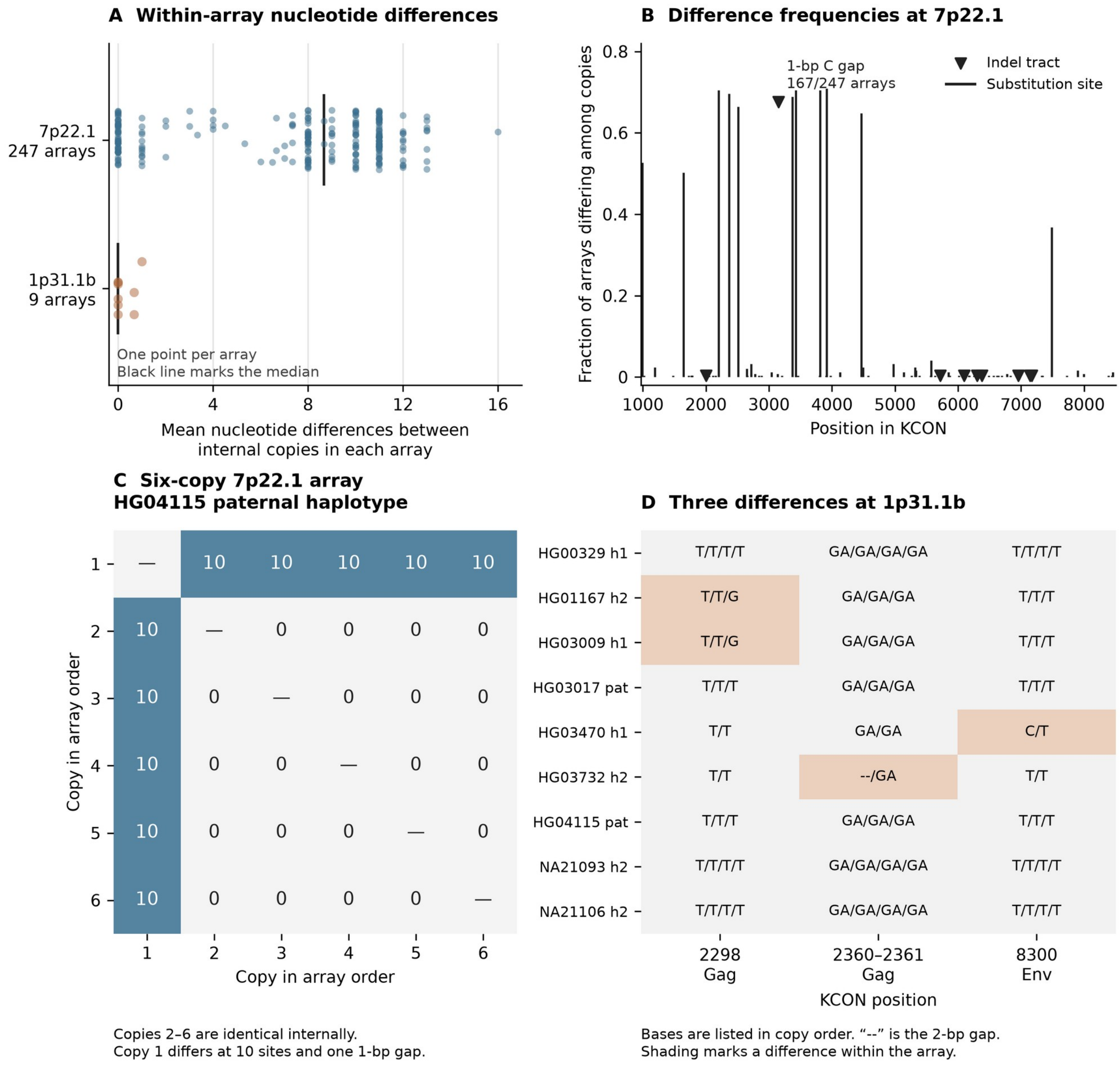


**Figure S5.** Nucleotide differences within tandem arrays. **(A)** Mean pairwise substitution counts within each array. Black lines mark medians. **(B)** Fraction of 7p22.1 arrays containing a difference at each position. Black lines denote substitutions, and black triangles denote gap tracts. **(C)** Pairwise substitution counts in the six-copy HG04115 paternal 7p22.1 array. **(D)** Variable sites in 1p31.1b arrays. Shading marks within-array differences. Dashes denote a two-base gap. Copies follow proviral orientation. Positions use Type-II KCON coordinates.

**S6A Candidate differences between paired target-site duplications**

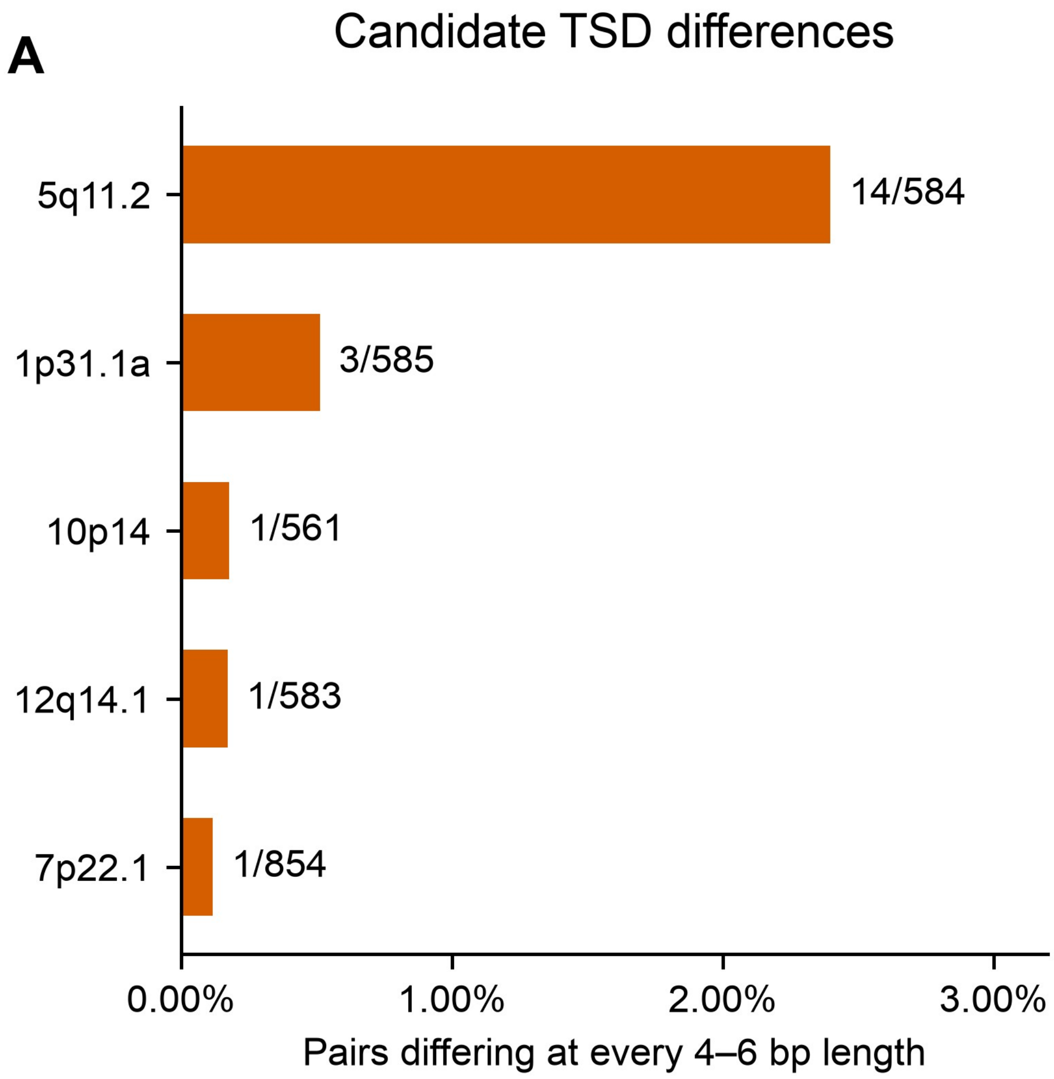

**S6B The candidate TSD length at 8p23.1a**

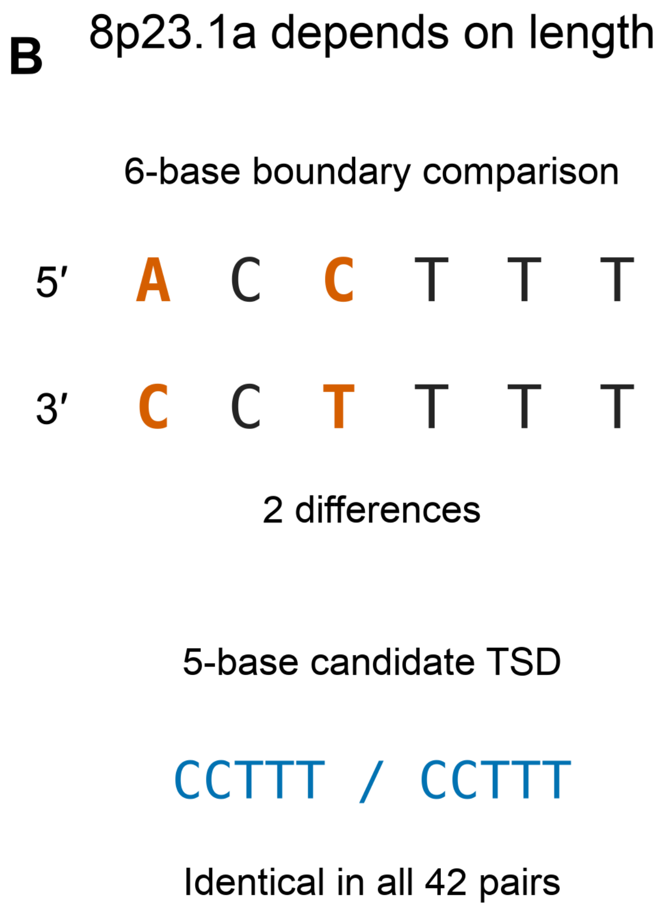


**Figure S6:** Candidate HML-2 target-site duplication differences. **(A)** Fraction of paired boundaries that differ at every candidate TSD length from 4 to 6 bp. Labels indicate differing/paired observations. **(B)** The 8p23.1a boundary comparison. All 42 six-base pairs differ, but the adjacent five-base pairs match.

### S7A Gag phylogeny

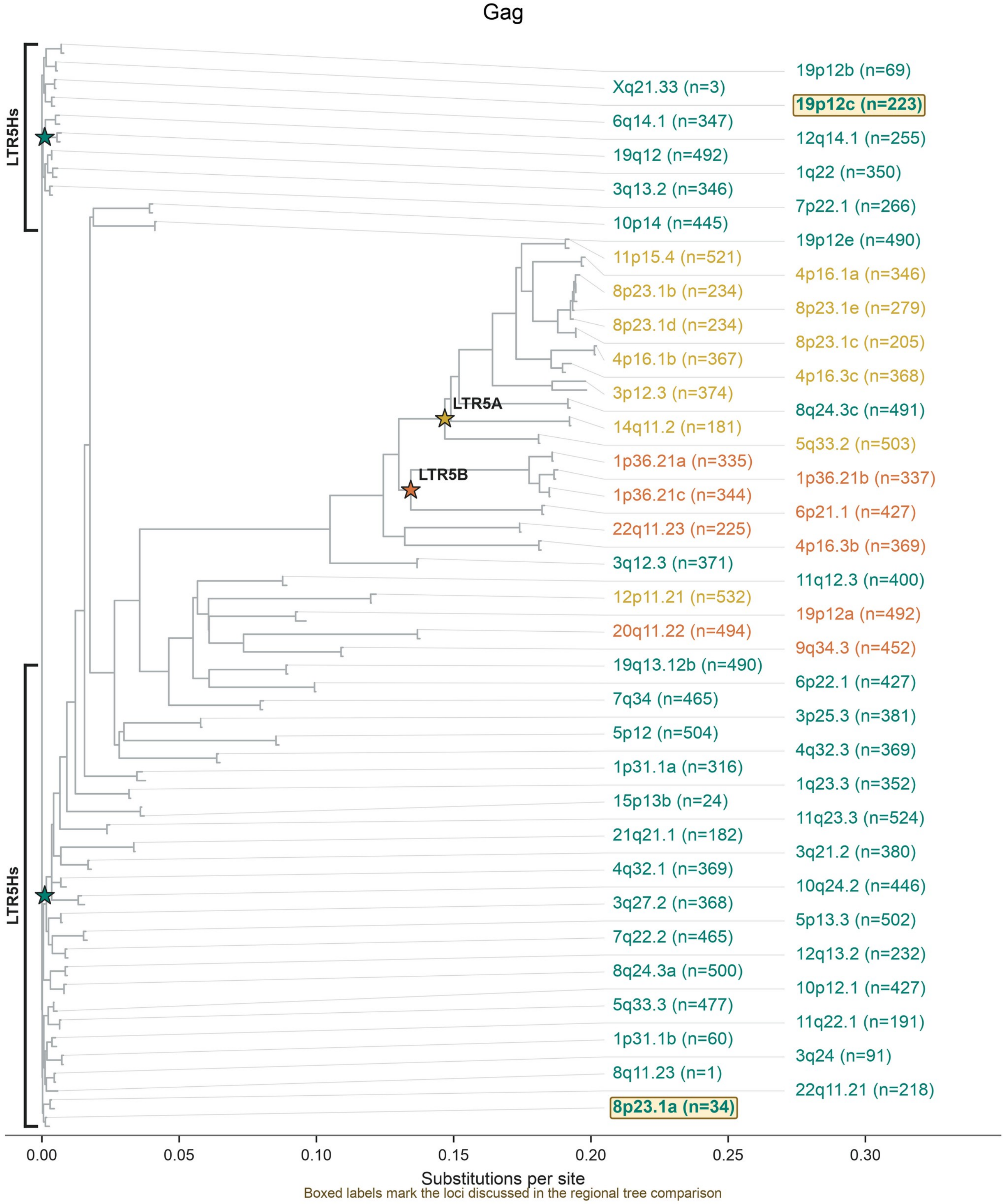


Boxed labels mark the loci discussed in the regional tree comparison

n = retained source-bound sequences; up to two modal clusters per locus

Teal: LTR5Hs Gold: LTR5A Orange: LTR5B; HML-11 comparators excluded

**S7B Pro phylogeny**

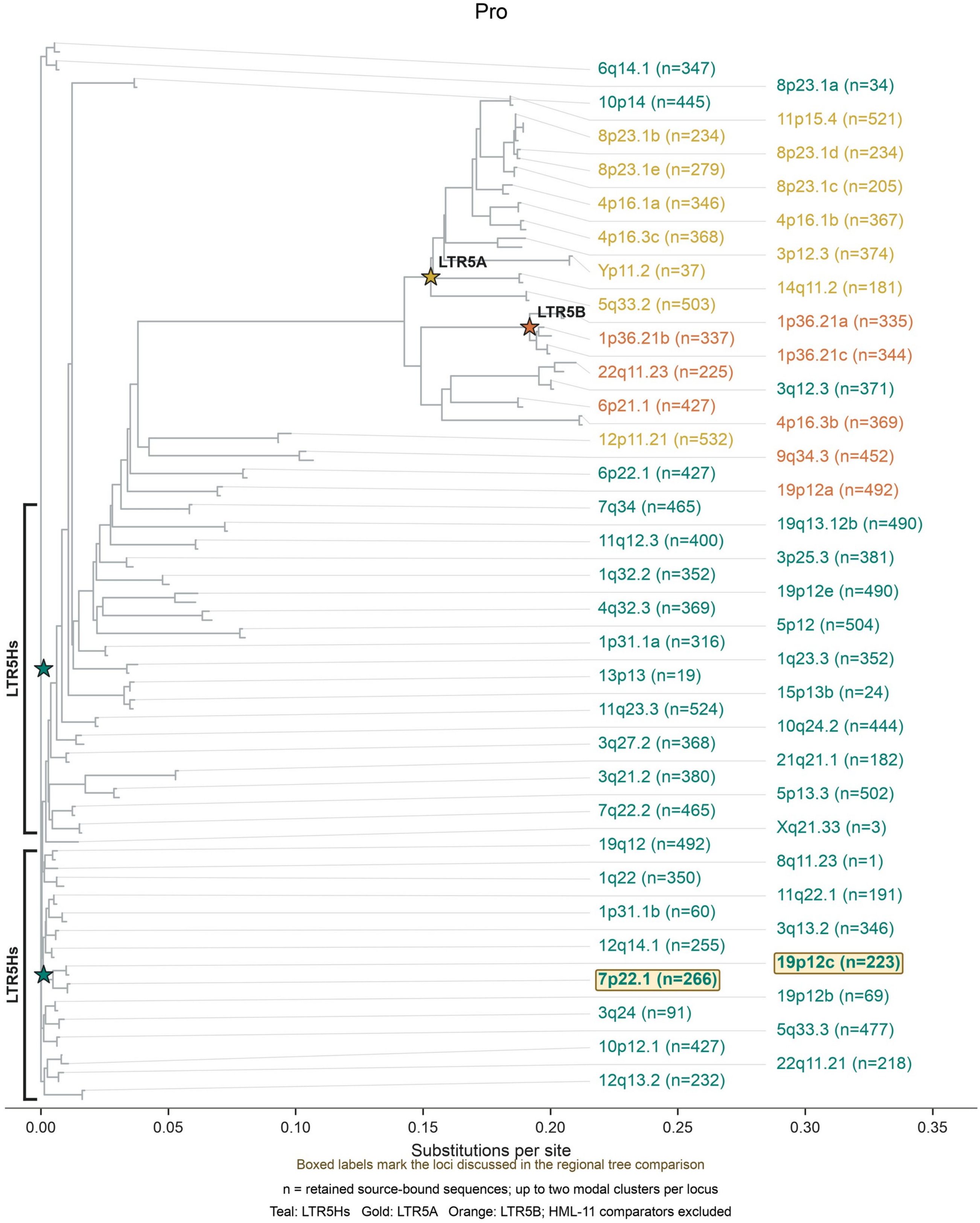


**Figure S7:** Nucleotide neighbor-joining trees of **(A)** gag and **(B)** pro. Up to two sequence clusters are shown per locus. Labels show the locus, LTR subfamily, and alignment-observation count. Boxes mark loci discussed in the text.

### S8A LTR phylogeny

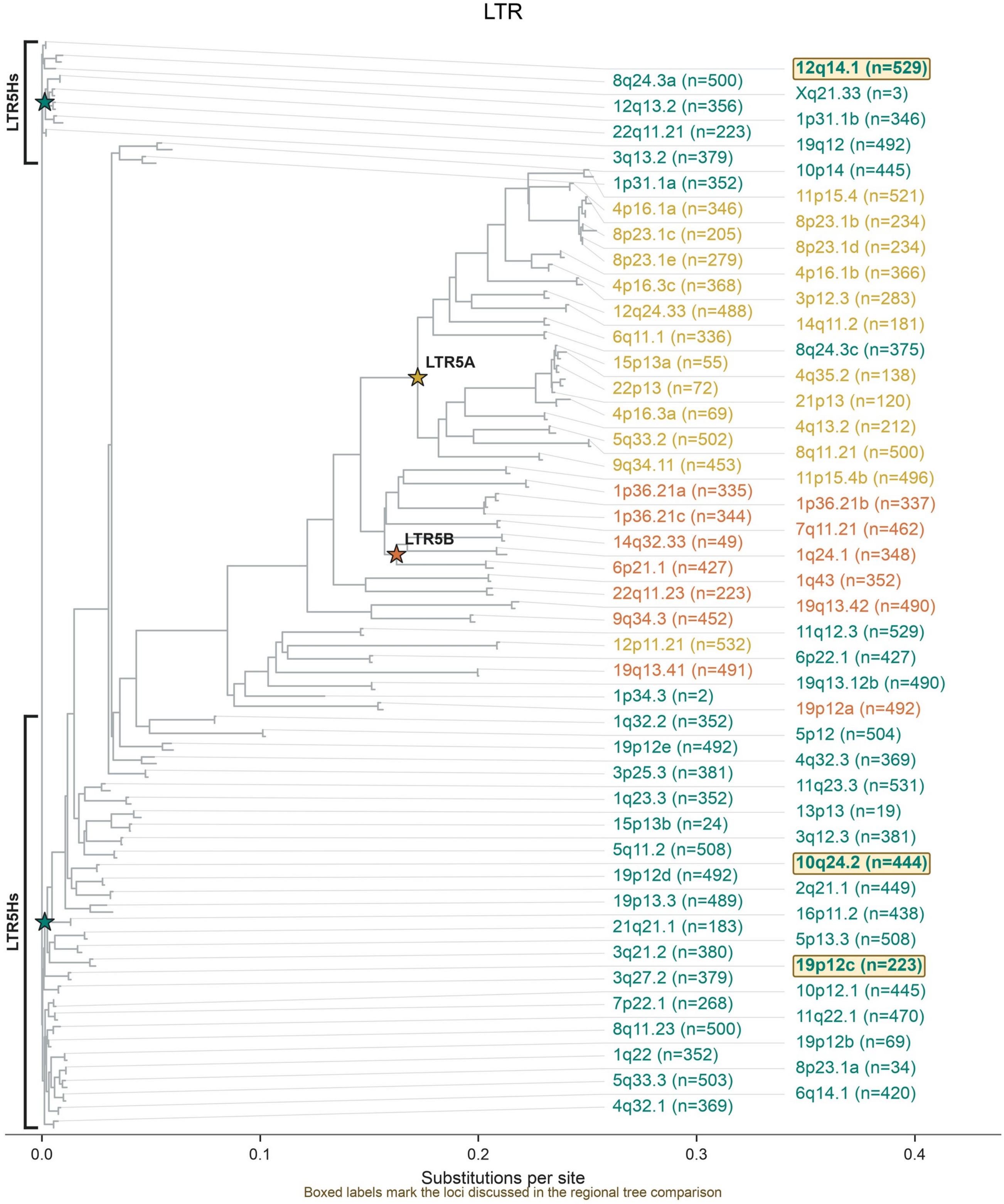


Boxed labels mark the loci discussed in the regional tree comparison

n = retained source-bound sequences; up to two modal clusters per locus

Teal: LTR5Hs Gold: LTR5A Orange: LTR5B; HML-11 comparators excluded

### S8B Env phylogeny

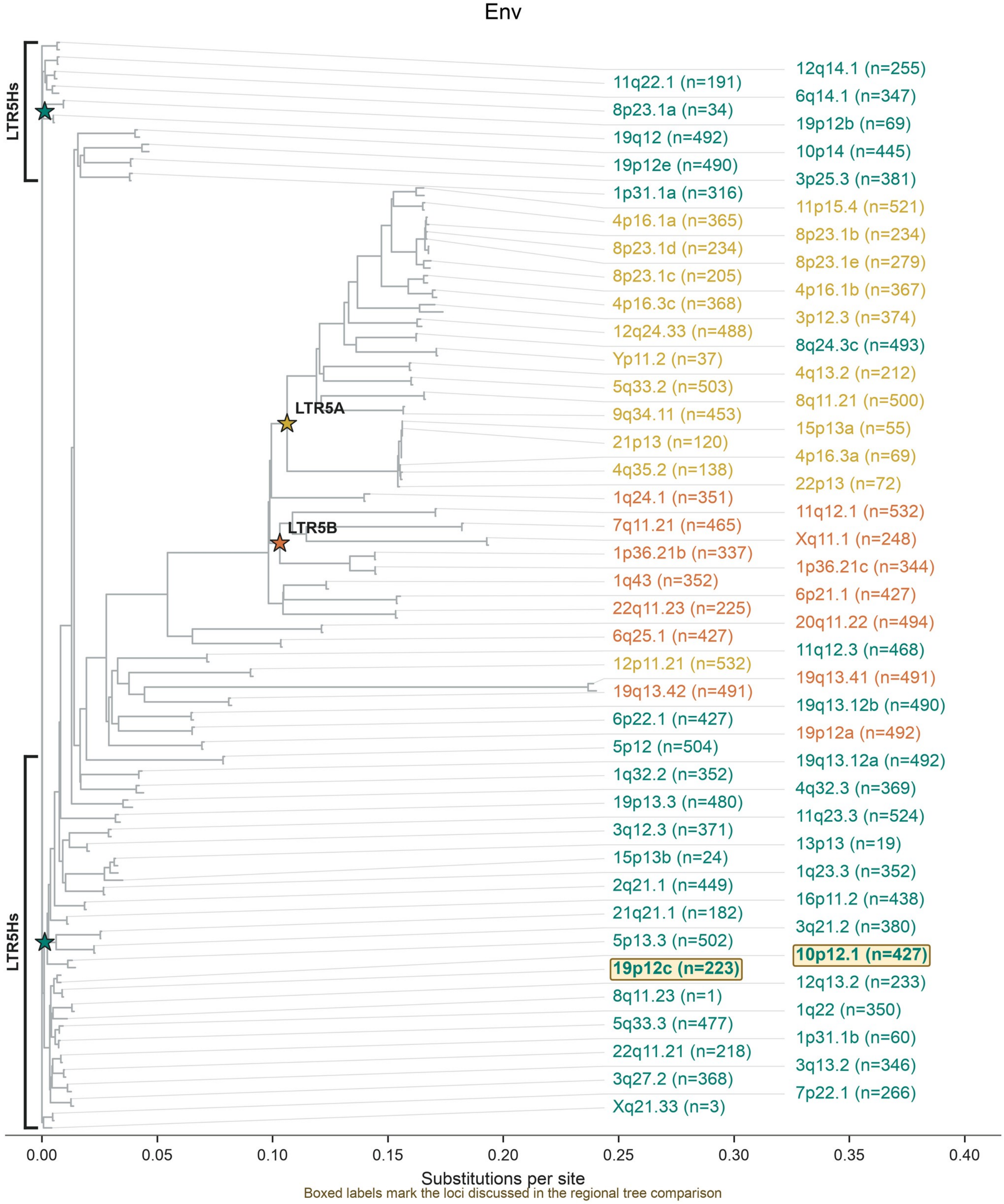


Boxed labels mark the loci discussed in the regional tree comparison

n = retained source-bound sequences; up to two modal clusters per locus

Teal: LTR5Hs Gold: LTR5A Orange: LTR5B; HML-11 comparators excluded

### S8C Pol phylogeny

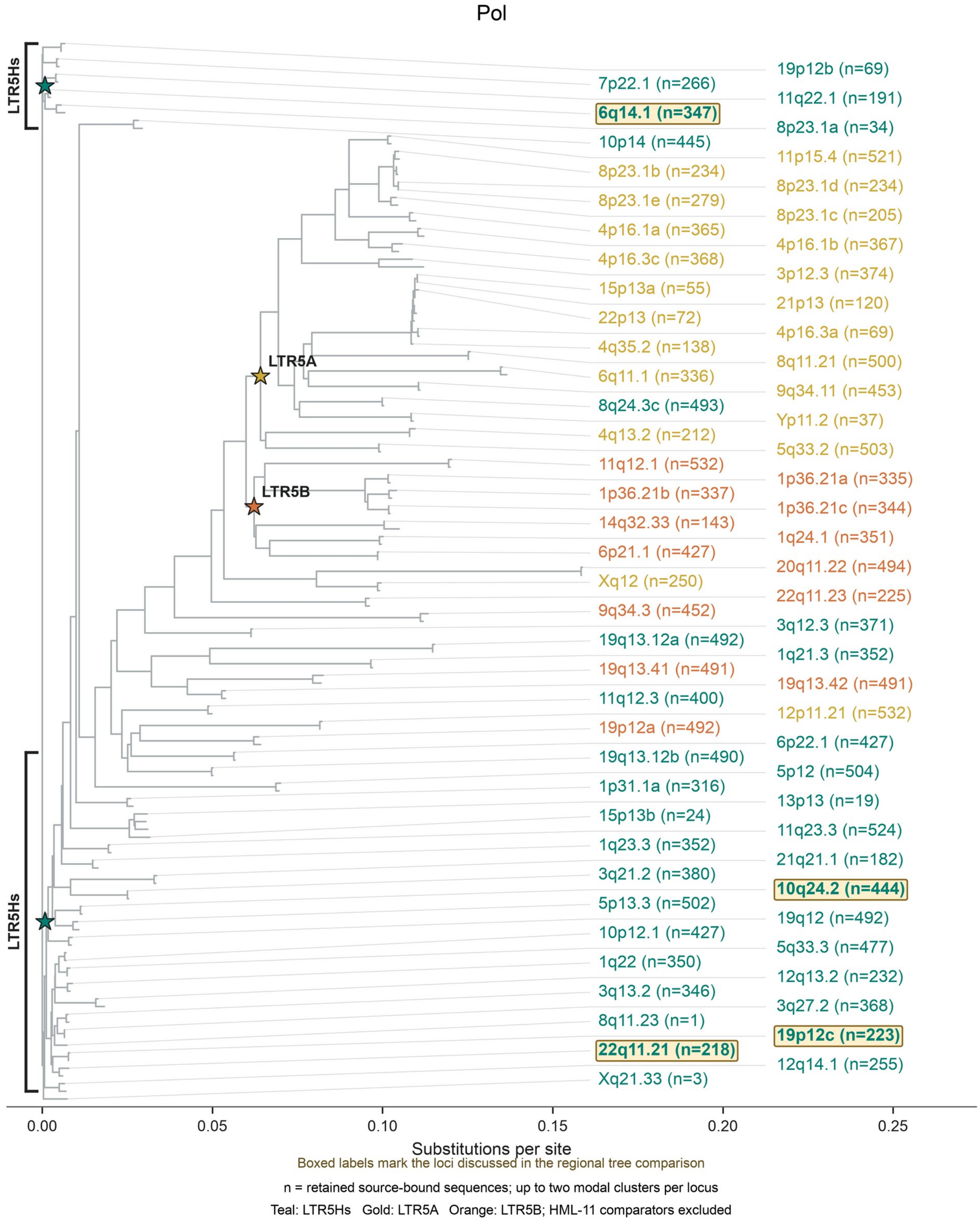


**Figure S8:** Nucleotide neighbor-joining trees of **(A)** LTR, **(B)** env, and **(C)** pol regions from 82, 76, and 70 loci, respectively. Up to two sequence clusters are shown per locus. Labels give LTR subfamily and alignment-observation count. Boxes mark loci discussed in the text.

**S9A Protein-changing variants**

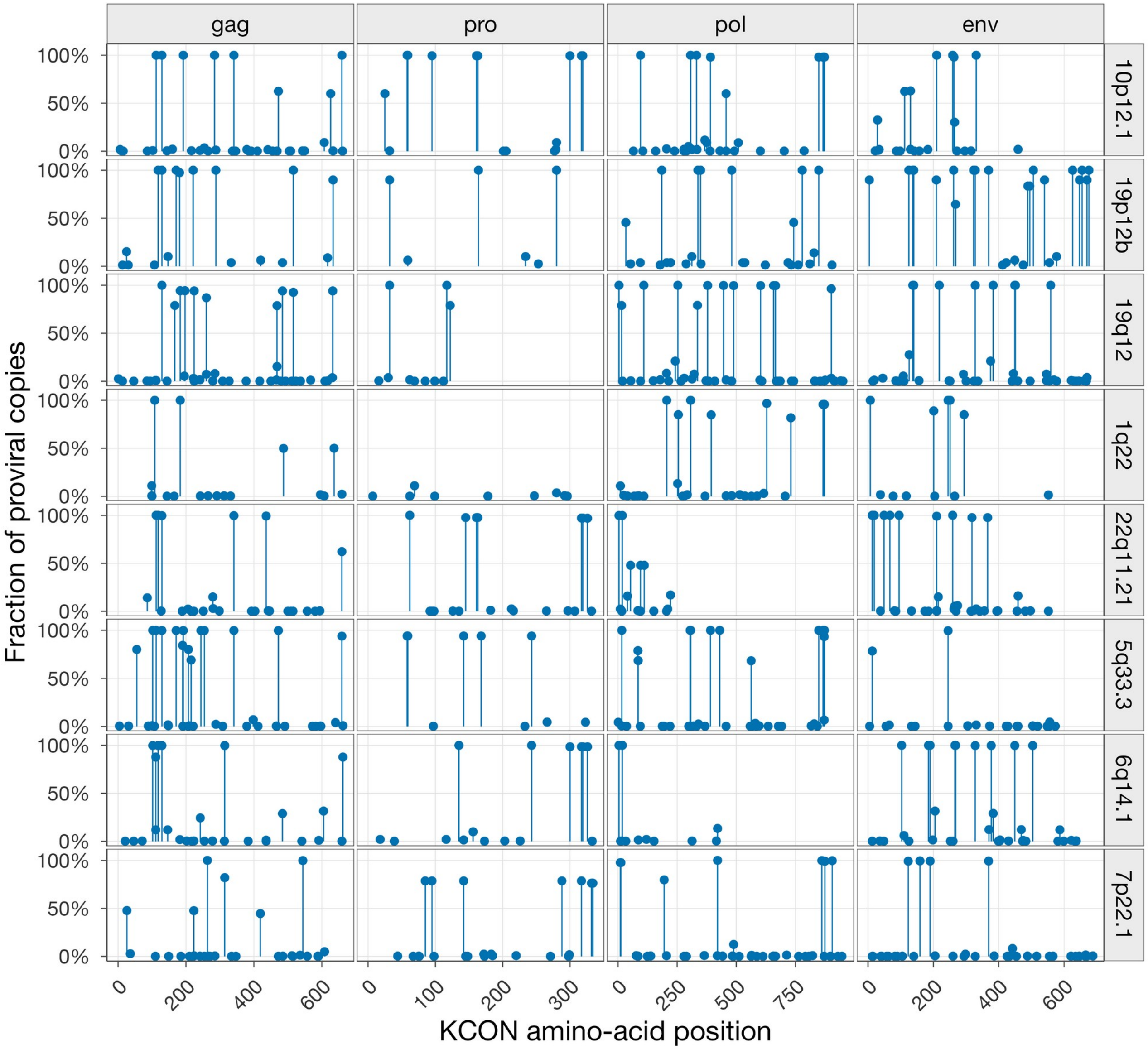

**S9B Frameshifts and Pol Y195C**

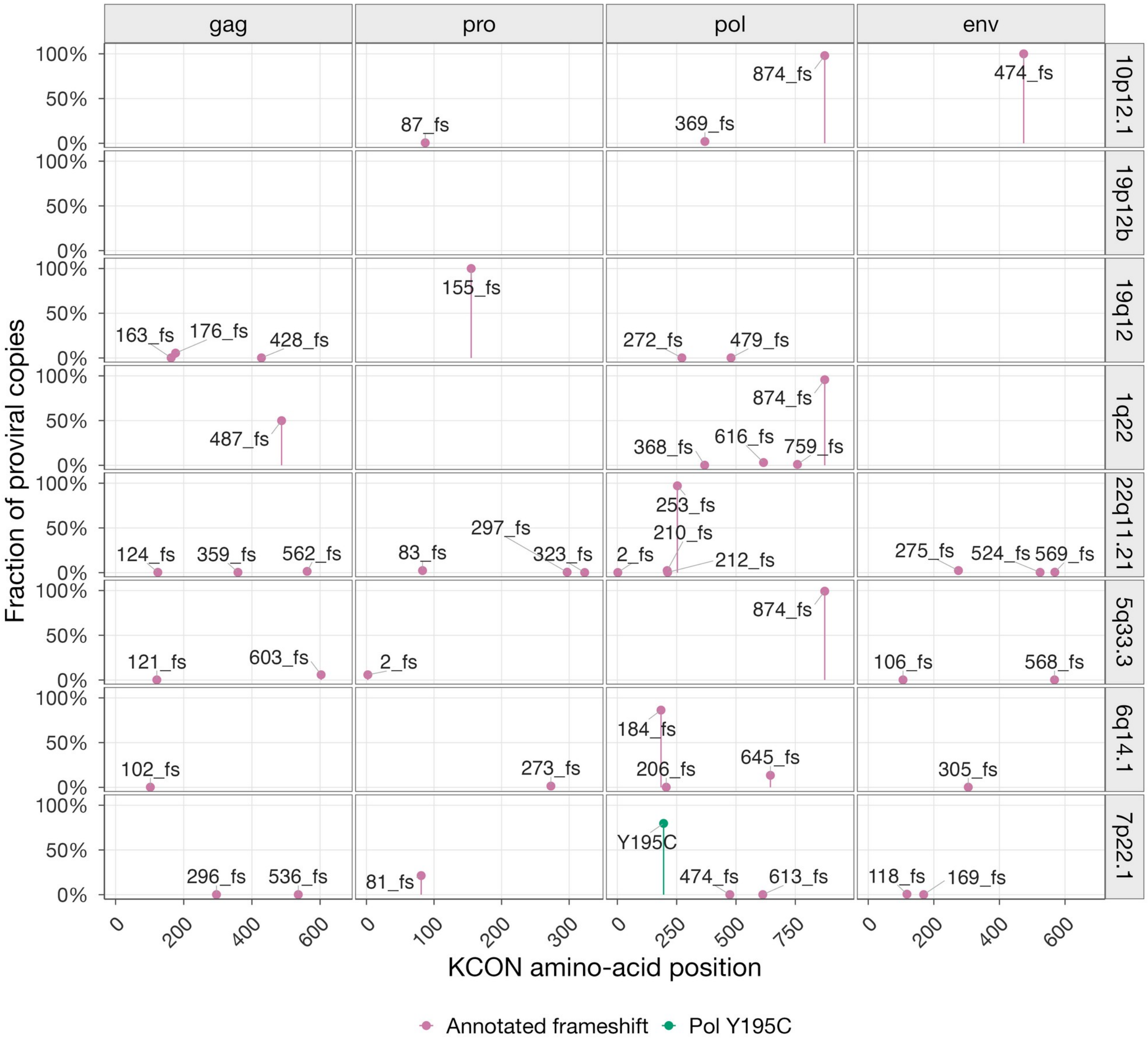


**Figure S9:** Protein-variant positions at eight HML-2 loci. **(A)** Amino-acid substitutions. **(B)** Frameshift positions (pink) and Pol Y195C at 7p22.1 (green). Y195C changes the conserved reverse-transcriptase YIDD motif to CIDD. Coordinates refer to KCON proteins. Frequencies use retained proviral copies at each locus.

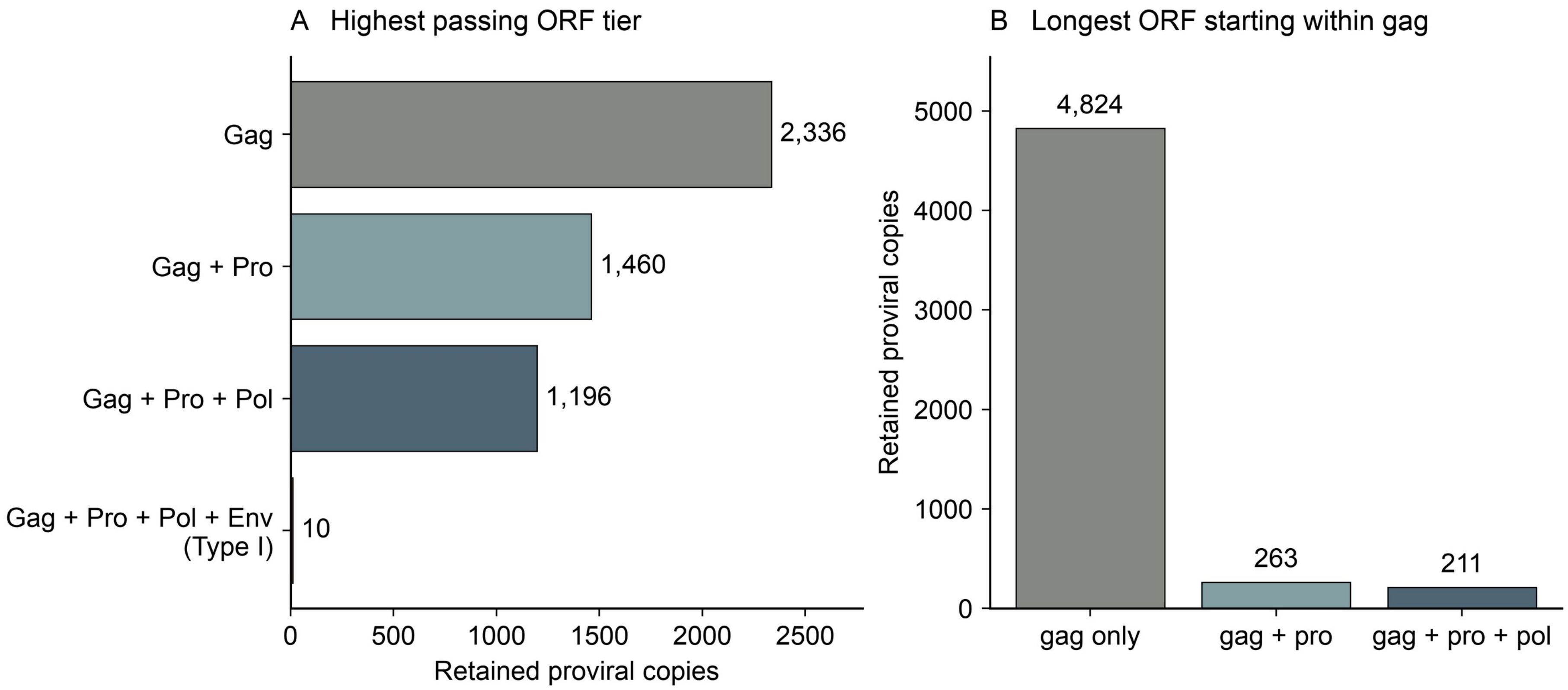


**Figure S10:** ORF combinations and single-frame coding sequences. **(A)** Longest consecutive Gag–Pro–Pol combination passing the combined screen. The fourth group also requires the theoretical Type-I Env annotation. Translation of this N-terminally truncated product has not been demonstrated. **(B)** Gene regions overlapped by the longest ATG-starting ORF of at least 200 amino acids that begins in gag.

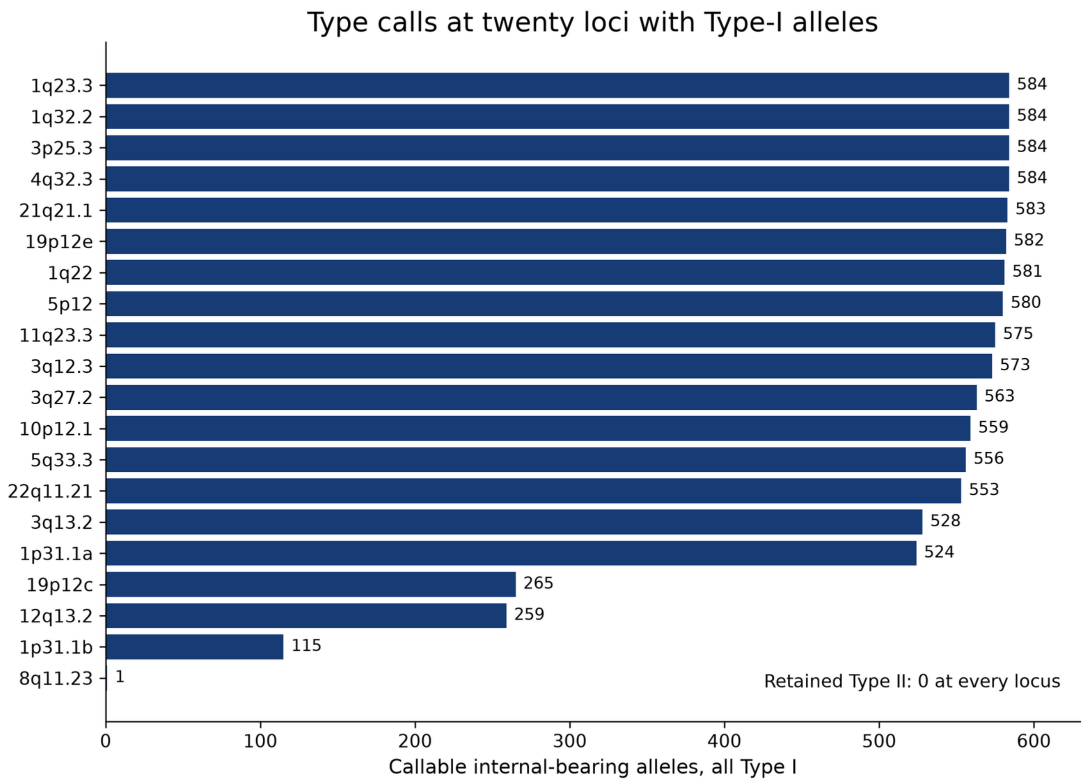


**Figure S11:** Direct Type-I cassette calls. Labels give callable haplotype counts. Each donor-haplotype-locus combination is counted once. Table S12 provides the underlying calls.

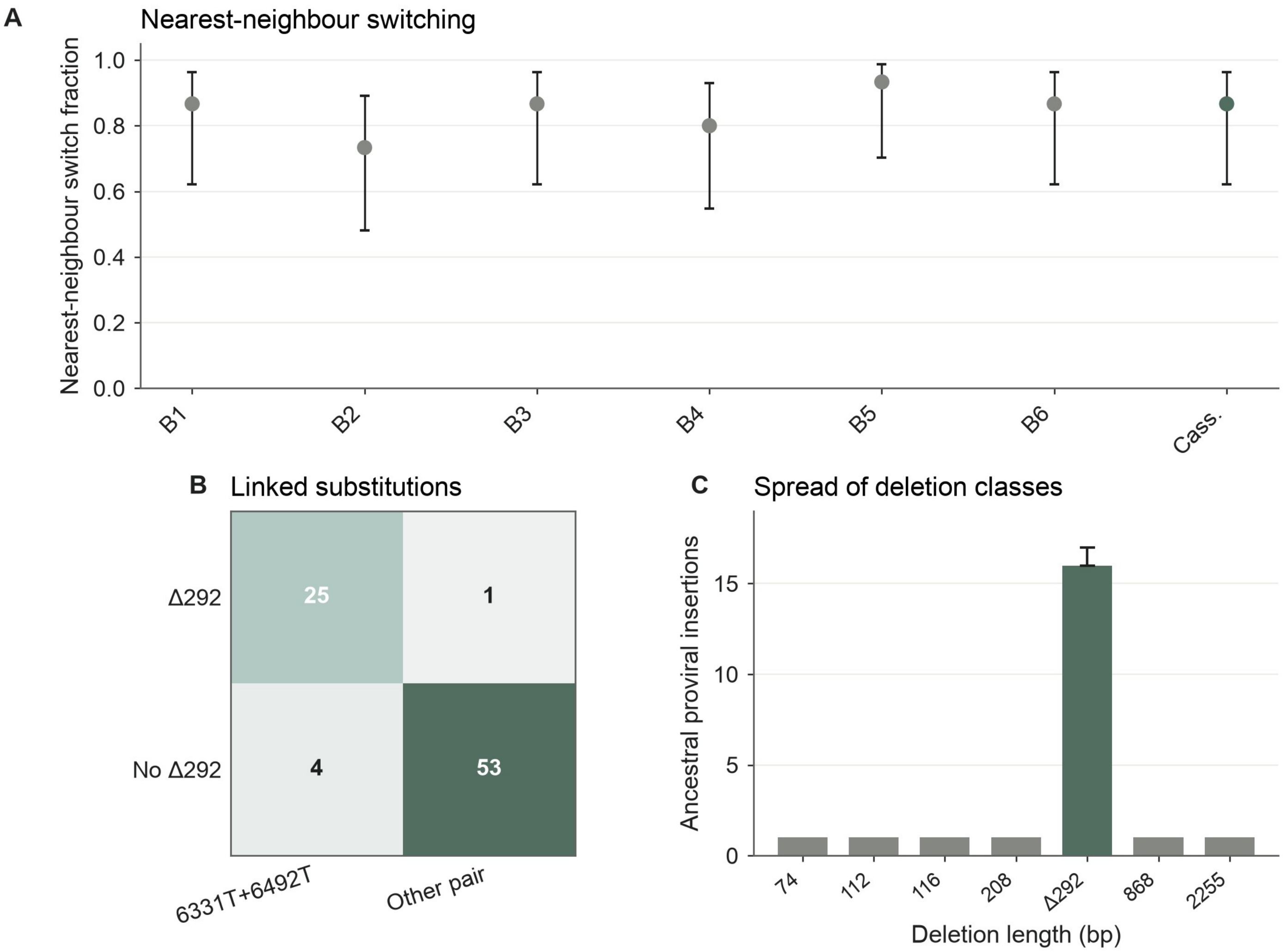


Figure S12: Sequence comparisons across the Type-I cassette. (A) Fraction of 15 Type-I representatives whose nearest Type-II neighbor changes between each window and the remaining sequence. Windows match Figure 5A. Bars show binomial 95% intervals. (B) Δ292 and the linked T/T sites in 83 nonhuman primate sequence clusters. Orthologs from different species are separate observations. (C) Ancestral proviral insertions carrying Δ292 and six other large deletions detected in the primate panel. Host-flank orthology distinguishes 16 Δ292-bearing insertions; the interval extends to one possible additional integration.

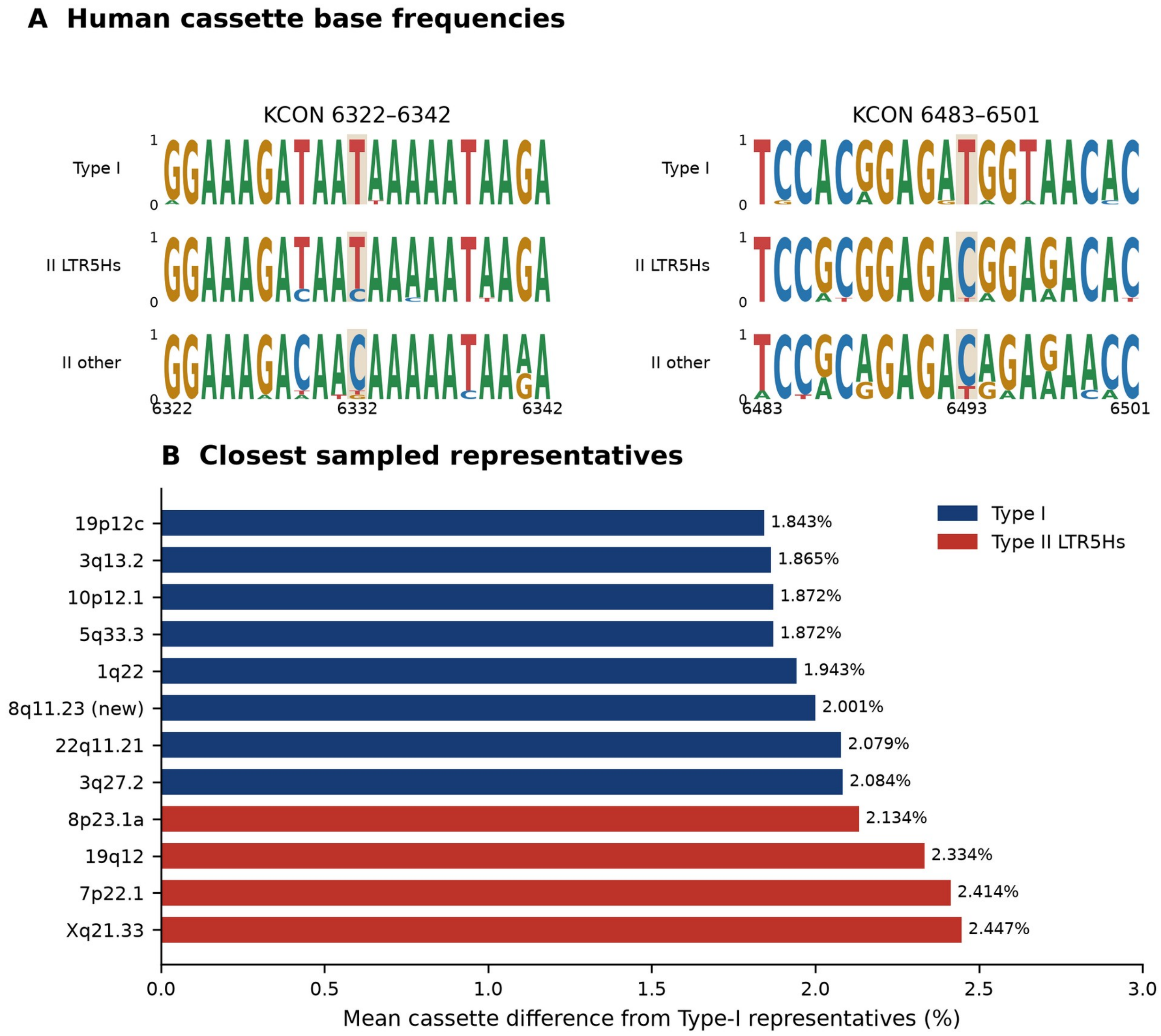


**Figure S13.** Cassette variation among human proviruses. **(A)** Nucleotide-frequency logos around KCON sites 6332 and 6493 (gold). Each group contains 15 representatives. Coordinates are one-based. **(B)** Sampled representatives ranked by mean cassette divergence from the Type-I panel. Complete aligned FASTA files are in the Table S11 supporting data.

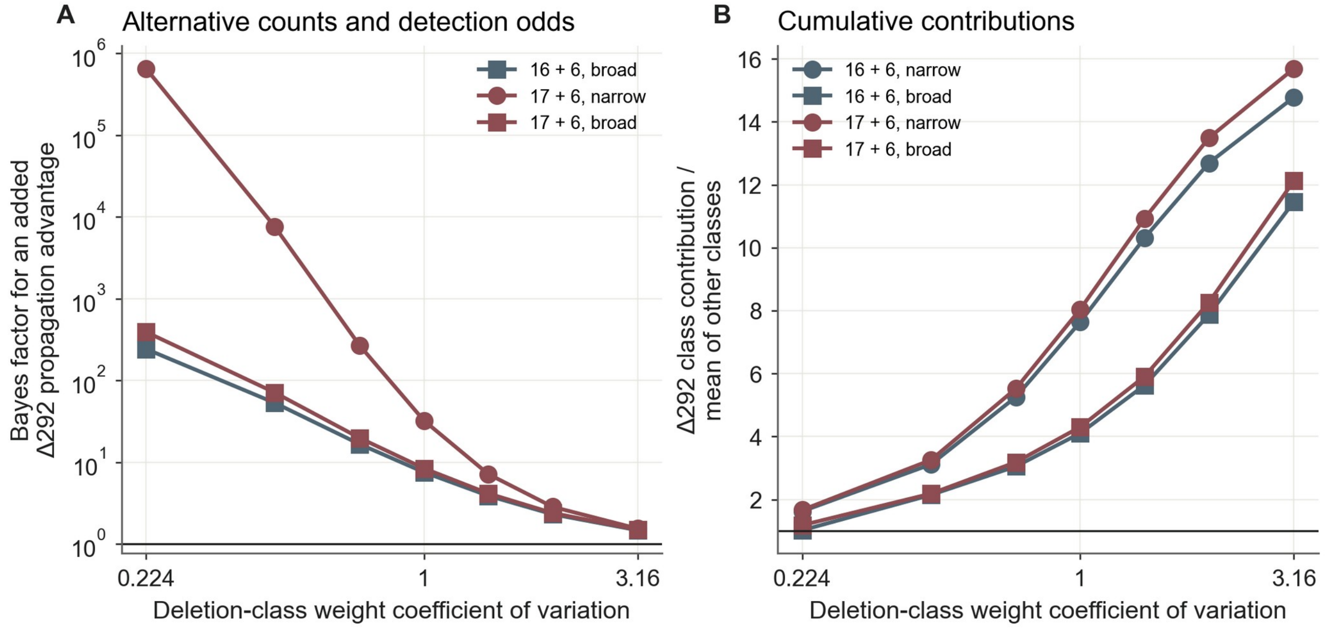


Figure S14: Deletion-class contributions and Δ292 enrichment. (A) Bayes factors for an added Δ292 propagation advantage under the conservative 16-plus-six count and the inclusive 17-plus-six count, and alternative detection odds. The primary conservative-count, narrow-odds curve is shown in Figure 5D. (B) Median relative contribution of the Δ292 deletion class in the model without an added deletion-specific effect. Narrow and broad relative detection odds range from 0.67–1.5 and 0.25–4, respectively.

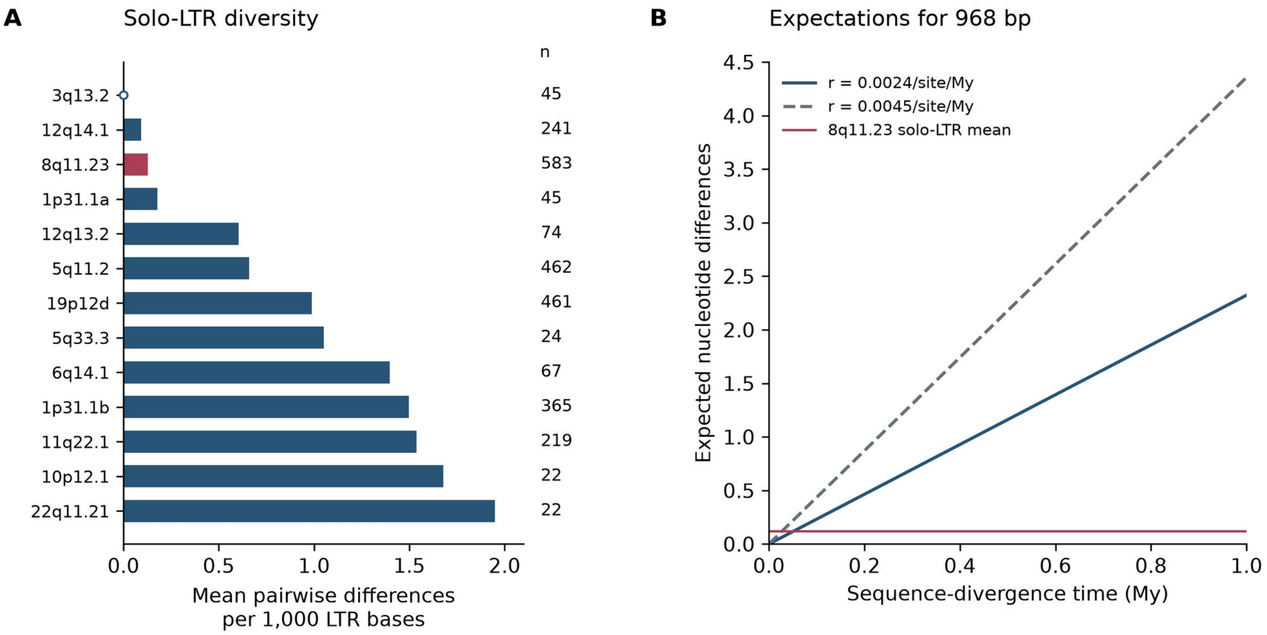


Figure S15. Solo-LTR variation and substitution expectations. (A) Nucleotide diversity at 13 loci. Red marks 8q11.23. Labels give sequence counts. (B) Expected differences across 968 bp under two pairwise substitution rates. The red line marks the observed mean among 8q11.23 solo-LTR sequences. The time axis describes sequence divergence, not an inferred age of the occupied insertion site.

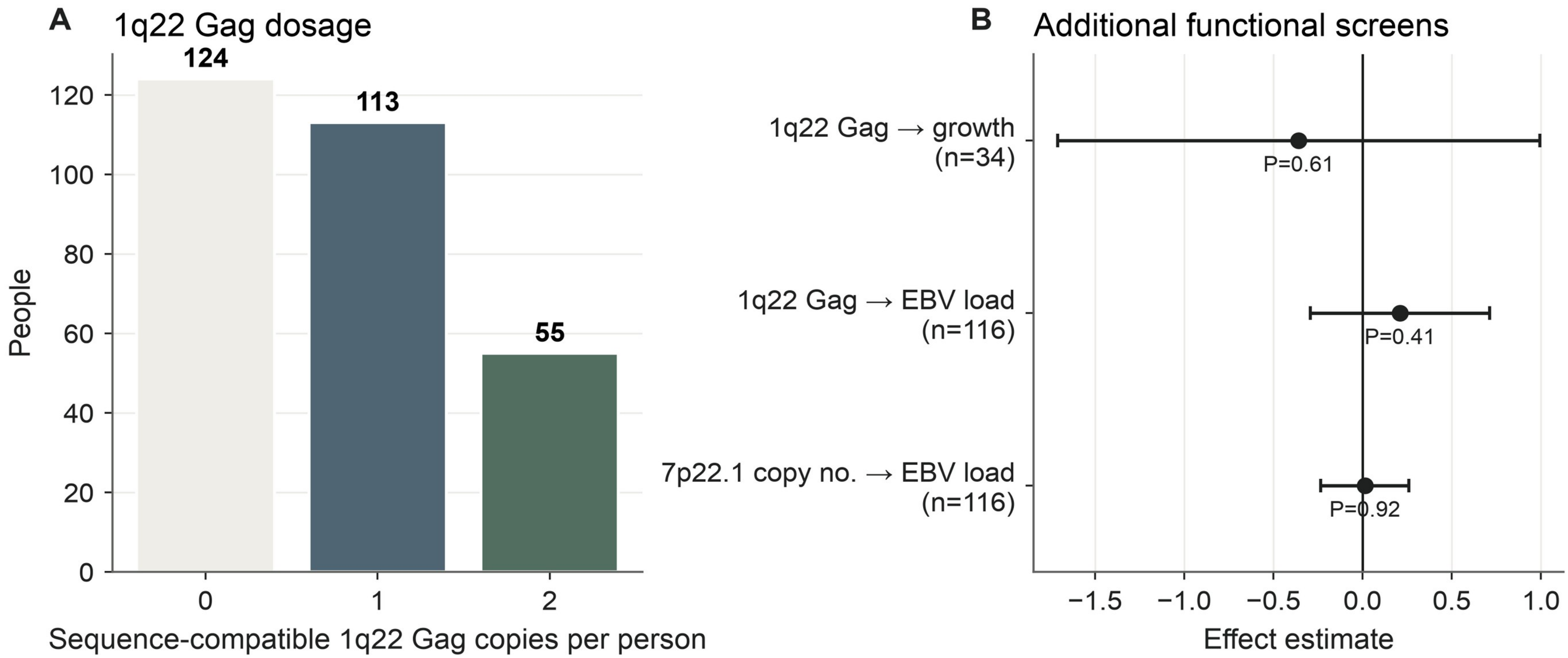


Figure S16: Proviral variation, cell growth and Epstein–Barr virus abundance in donor-derived lymphoblastoid cell lines. (A) 1q22 Gag copy counts under the combined ORF screen. (B) The first two rows compare donors carrying at least one 1q22 Gag copy passing the combined screen with donors carrying none. The third row gives the effect per additional diploid 7p22.1 copy. Effects are expressed as changes in growth divided by 10,000 or in $\log_2$ EBV DNA abundance. Points show coefficients; intervals span ±1.96 HC3 standard errors.

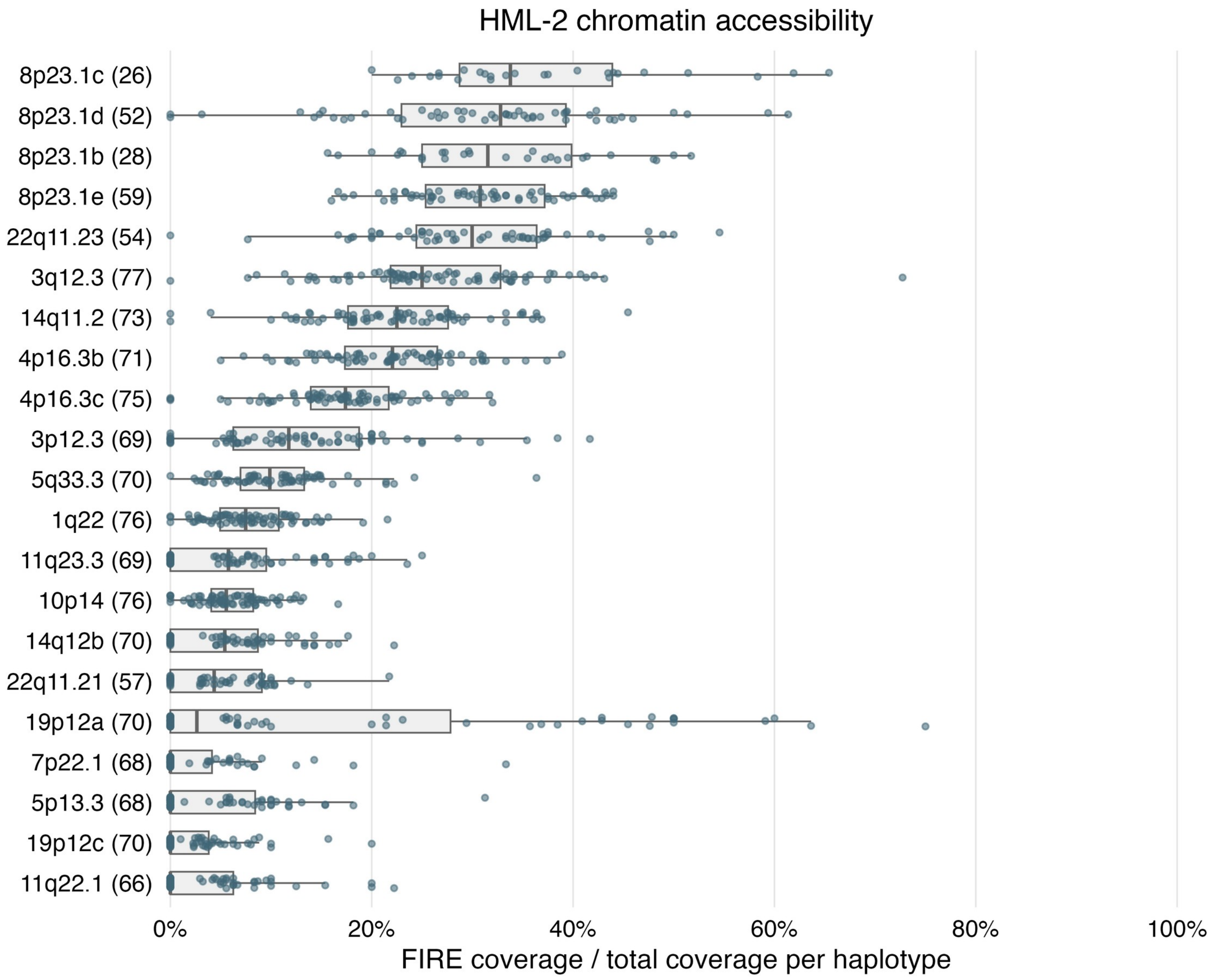


Figure S17: HML-2 chromatin accessibility in lymphoblastoid cell lines from 39 donors. Each point represents a primary haplotype. Parentheses give haplotype counts. Boxes show medians and interquartile ranges. Whiskers extend to the most extreme observations within 1.5 times the interquartile range. The 7p22.1 assay covers one proviral segment.

**Supplementary Tables**

**Table S1. HML-2 structural-analysis catalog**

This table summarizes 103 HML-2 labels: 99 named loci and four copy groups. Named loci give called/total chromosome counts and structural fractions among called haplotypes. Copy groups have no single-locus denominator. Counts exclude alias labels and unsupported copies. HML-11 sequences are retained separately. The last column reports selected combinations that pass the combined ORF screen: Gag, Gag–Pro, Gag–Pro–Pol, Env, Np9, and Rec. It does not enumerate isolated Pro or Pol calls. *Env at Type-I loci denotes the theoretical N-terminally truncated product whose translation has not been demonstrated.

| Catalog label | Former name(s) | Type | T2T-CHM13 coordinates | Maximum retained copies per haplotype | Maximum array size | Structural observations | Selected ORF combinations |
|---|---|---|---|---|---|---|---|
| 10p12.1 | HERV-K103, K(C10) | Type I | chr10:26915413-26924592 | 2 | 2 | Called 584/584. provirus 558/584, multi-copy 1/584, solo-ltr 25/584 | Gag, Gag–Pro, Gag–Pro–Pol, Np9 |
| 10p14 | K(C11a), K33, ERVK-16 | Type II | chr10:6824399-6833862 | 1 | 1 | Called 583/584. provirus 582/583, solo-ltr 1/583 | Rec |
| 10q24.2 | HERV-K128, ERVK-17, c10_B | Type II | chr10:100703746-100711001 | 1 | 1 | Called 584/584. provirus 582/584, fragment 2/584 | Rec |
| 11p15.4 | K7, ERVK3-4 | Type II | chr11:3492878-3502431 | 1 | 1 | Called 582/584. provirus 573/582, noncarrier call 9/582 | Rec |
| 11p15.4b | | Type II | chr11:4524557-4526594 | 1 | 0 | Called 584/584. solo-ltr 548/584, noncarrier call 36/584 | None |
| 11q12.1 | | Type II | chr11:58949479-58955236 | 1 | 0 | Called 584/584. fragment 584/584 | None |
| 11q12.3 | | Type II | chr11:62357829-62360337 | 2 | 1 | Called 584/584. provirus 379/584, multi-copy 1/584, fragment 202/584, noncarrier call 2/584 | None |
| 11q22.1 | HERV-K118, K(C11c), K36, ERVK-25 | Type II | chr11:101694640-101715612 | 2 | 2 | Called 584/584. provirus 213/584, multi-copy 3/584, solo-ltr 304/584, noncarrier call 64/584 | Rec |
| 11q23.3 | K(C11b), K37, ERVK-20 | Type I | chr11:118740087-118749246 | 2 | 2 | Called 584/584. provirus 574/584, multi-copy 1/584, solo-ltr 9/584 | Np9 |

| Catalog label | Former name(s) | Type | T2T-CHM13 coordinates | Maximum retained copies per haplotype | Maximum array size | Structural observations | Selected ORF combinations |
|---|---|---|---|---|---|---|---|
| 12p11.21 | | Type II | chr12:34497120-34506786 | 1 | 1 | Called 584/584. provirus 584/584 | Rec |
| 12q13.2 | HERV-K134 | Type I | chr12:55294991-55305959 | 2 | 2 | Called 584/584. provirus 258/584, multi-copy 1/584, solo-ltr 130/584, noncarrier call 195/584 | Gag, Gag–Pro, Env* |
| 12q14.1 | HERV-K119, K(C12), K41, ERVK-21 | Type II | chr12:58305265-58314721 | 2 | 2 | Called 582/584. provirus 274/582, multi-copy 1/582, solo-ltr 307/582 | Gag, Gag–Pro, Env, Rec |
| 12q24.33 | K42 | Type II | chr12:133147853-133153794 | 4 | 4 | Called 571/584. multi-copy 13/571, solo-ltr 20/571, fragment 537/571, noncarrier call 1/571 | None |
| 13p13 | | Type I | chr13:5381952-5388076 | 3 | | Called 469/584. multi-copy 45/469, solo-ltr 4/469, fragment 173/469, noncarrier call 247/469 | None |
| 14q11.2 | K(OLDAL136419), K71 | Type II | chr14:18172903-18178983 | 3 | 2 | Called 584/584. multi-copy 1/584, fragment 580/584, noncarrier call 3/584 | None |
| 14q32.33 | | Type II | chr14:99945707-99948598 | 2 | 0 | Called 556/584. multi-copy 5/556, fragment 520/556, noncarrier call 31/556 | None |
| 15p13a | | Type II | chr15:12216-19499 | 4 | | Called 438/584. multi-copy 131/438, fragment 193/438, noncarrier call 114/438 | None |
| 15p13b | | Type I | chr15:2092086-2101275 | 4 | | Called 347/584. provirus 186/347, multi-copy 50/347, solo-ltr 23/347, fragment 26/347, noncarrier call 62/347 | Np9 |
| 15q25.2 | | Type II | chr15:82026222-82030353 | 1 | 0 | Called 584/584. solo-ltr 461/584, fragment 122/584, noncarrier call 1/584 | None |

| Catalog label | Former name(s) | Type | T2T-CHM13 coordinates | Maximum retained copies per haplotype | Maximum array size | Structural observations | Selected ORF combinations |
|---|---|---|---|---|---|---|---|
| 16p11.2 | | Type II | chr16:39129141-39131888 | 2 | 0 | Called 580/584. multi-copy 3/580, fragment 571/580, noncarrier call 6/580 | None |
| 19p12a | K52 | Type II | chr19:20414092-20424206 | 1 | 1 | Called 584/584. provirus 584/584 | None |
| 19p12b | HERV-K113, De1, ERVK26 | Type II | chr19:21795000-21809000 | 1 | 1 | Called 584/584. provirus 79/584, noncarrier call 505/584 | Gag, Gag–Pro, Gag–Pro–Pol, Env, Rec |
| 19p12c | | Type I | chr19:22713637-22720374 | 1 | 1 | Called 584/584. provirus 265/584, noncarrier call 319/584 | Env*, Np9 |
| 19p12d | De11 | Type I | chr19:22365220-22383437 | 1 | 1 | Called 584/584. provirus 1/584, solo-ltr 583/584 | Gag, Np9 |
| 19p12e | K51 | Type I | chr19:22416059-22430176 | 1 | 0 | Called 584/584. solo-ltr 2/584, fragment 582/584 | None |
| 19p13.3 | ERVK-22 | Type II | chr19:333888-336426 | 1 | 0 | Called 580/584. solo-ltr 9/580, fragment 569/580, noncarrier call 2/580 | None |
| 19q12 | | Type II | chr19:30156120-30164983 | 2 | 1 | Called 584/584. provirus 582/584, multi-copy 1/584, noncarrier call 1/584 | Gag, Env, Rec |
| 19q13.12a | | Type II | chr19:38117259-38121486 | 1 | 0 | Called 584/584. fragment 584/584 | None |
| 19q13.12b | K(OLDAC012309), KOLD12309, K50F | Type II | chr19:38361524-38365751 | 1 | 1 | Called 582/584. provirus 582/582 | None |
| 19q13.41 | | Type II | chr19:55829484-55834919 | 1 | 0 | Called 583/584. fragment 583/583 | None |
| 19q13.42 | LTR13 | Type II | chr19:56438594-56444294 | 1 | 0 | Called 583/584. fragment 583/583 | None |
| 1p31.1a | | Type I | chr1:72962780-72971964 | 2 | 1 | Called 584/584. provirus 523/584, multi-copy 1/584, solo-ltr 60/584 | None |
| 1p31.1b | HERV-K116, K4, ERVK-1 | Type I | chr1:82060947-82061157 | 4 | 4 | Called 584/584. multi-copy 9/584, solo-ltr 469/584, fragment 106/584 | Env*, Np9 |

| Catalog label | Former name(s) | Type | T2T-CHM13 coordinates | Maximum retained copies per haplotype | Maximum array size | Structural observations | Selected ORF combinations |
|---|---|---|---|---|---|---|---|
| 1p34.3 | | Type II | chr1:36351865-36354008 | 1 | 0 | Called 584/584. fragment 584/584 | None |
| 1p36.21a | | Type II | chr1:12324389-12330209 | 1 | 1 | Called 584/584. provirus 1/584, solo-ltr 4/584, fragment 562/584, noncarrier call 17/584 | None |
| 1p36.21b | | Type II | chr1:12529193-12538734 | 1 | 1 | Called 581/584. provirus 558/581, noncarrier call 23/581 | None |
| 1p36.21c | HERV-K(OLDAL023753), K6, K76 | Type II | chr1:12793963-12803479 | 1 | 1 | Called 584/584. provirus 576/584, noncarrier call 8/584 | Rec |
| 1q21.3 | | Type II | chr1:149756845-149759931 | 1 | 0 | Called 584/584. fragment 584/584 | None |
| 1q22 | HERV-K102, K(C1b), K50a, ERVK-7 | Type I | chr1:154765232-154774409 | 1 | 1 | Called 584/584. provirus 581/584, solo-ltr 3/584 | Gag, Gag–Pro, Gag–Pro–Pol, Env*, Np9 |
| 1q23.3 | HERV-K110, K18, K(C1a), ERVK-18 | Type I | chr1:159827889-159837121 | 1 | 1 | Called 584/584. provirus 584/584 | None |
| 1q24.1 | K12 | Type II | chr1:165951274-165956929 | 1 | 0 | Called 584/584. fragment 583/584, noncarrier call 1/584 | None |
| 1q32.2 | | Type I | chr1:206881854-206886033 | 1 | 0 | Called 584/584. fragment 584/584 | None |
| 1q43 | | Type II | chr1:238174724-238176902 | 1 | 0 | Called 584/584. fragment 584/584 | None |
| 20q11.22 | K(OLDAL136419), K59 | Type II | chr20:35854131-35862963 | 2 | 0 | Called 582/584. multi-copy 1/582, fragment 580/582, noncarrier call 1/582 | None |
| 21p13 | | Type II | chr21:2699539-2710506 | 5 | | Called 443/584. multi-copy 259/443, fragment 144/443, noncarrier call 40/443 | None |
| 21q21.1 | HERV-K133, K60, ERVK-23 | Type I | chr21:16920971-16929274 | 1 | 1 | Called 584/584. provirus 583/584, solo-ltr 1/584 | Np9 |
| 22p13 | | Type II | chr22:10726-18005 | 4 | | Called 451/584. multi-copy 75/451, fragment 178/451, noncarrier call 198/451 | None |

| Catalog label | Former name(s) | Type | T2T-CHM13 coordinates | Maximum retained copies per haplotype | Maximum array size | Structural observations | Selected ORF combinations |
|---|---|---|---|---|---|---|---|
| 22q11.21 | HERV-K101, K(C22), ERVK-24 | Type I | chr22:19314068-19323246 | 1 | 1 | Called 581/584. provirus 553/581, solo-ltr 25/581, noncarrier call 3/581 | Gag, Gag–Pro, Env*, Np9 |
| 22q11.23 | K(OLDAP000345), KOLD345, ERVK-32 | Type II | chr22:23983293-23995659 | 3 | 1 | Called 581/584. provirus 573/581, multi-copy 2/581, fragment 5/581, noncarrier call 1/581 | None |
| 2q21.1 | HERV-K120 | Type I | chr2:130391600-130394679 | 1 | 0 | Called 583/584. fragment 583/583 | None |
| 3p12.3 | | Type II | chr3:75590244-75592148 | 1 | 1 | Called 581/584. provirus 574/581, noncarrier call 7/581 | Rec |
| 3p25.3 | K11, ERVK-2 | Type I | chr3:9839669-9846559 | 1 | 0 | Called 584/584. fragment 584/584 | None |
| 3q12.3 | K(II), ERVK-5 | Type I | chr3:104404564-104413685 | 1 | 1 | Called 584/584. provirus 573/584, solo-ltr 5/584, fragment 6/584 | Gag, Np9 |
| 3q13.2 | HERV-K106, K(C3), K68, ERVK-3 | Type I | chr3:115745233-115754391 | 2 | 2 | Called 584/584. provirus 525/584, multi-copy 3/584, solo-ltr 56/584 | Gag, Np9 |
| 3q21.2 | HERV-K121, K(I), ERVK-4 | Type II | chr3:128531680-128533592 | 1 | 1 | Called 584/584. provirus 583/584, noncarrier call 1/584 | Rec |
| 3q24 | HERV-K122, ERVK-13 | Type II | chr3:151308675-151328606 | 1 | 0 | Called 584/584. fragment 134/584, noncarrier call 450/584 | None |
| 3q27.2 | HERV-K117, K50b, ERVK-11 | Type I | chr3:188378372-188387551 | 2 | 2 | Called 584/584. provirus 561/584, multi-copy 2/584, solo-ltr 21/584 | Np9 |
| 4p16.1a | K17b | Type II | chr4:9001209-9003138 | 2 | 1 | Called 583/584. provirus 532/583, multi-copy 1/583, fragment 48/583, noncarrier call 2/583 | Rec |
| 4p16.1b | K50c | Type II | chr4:9088575-9098157 | 1 | 1 | Called 582/584. provirus 582/582 | Rec |

| Catalog label | Former name(s) | Type | T2T-CHM13 coordinates | Maximum retained copies per haplotype | Maximum array size | Structural observations | Selected ORF combinations |
|---|---|---|---|---|---|---|---|
| 4p16.3a | | Type II | chr4:12164-19439 | 2 | 0 | Called 572/584. multi-copy 4/572, solo-ltr 11/572, fragment 108/572, noncarrier call 449/572 | None |
| 4p16.3b | K77 | Type II | chr4:234544-239013 | 1 | 0 | Called 584/584. fragment 583/584, noncarrier call 1/584 | None |
| 4p16.3c | | Type II | chr4:3976712-3986301 | 1 | 1 | Called 584/584. provirus 583/584, noncarrier call 1/584 | Rec |
| 4q13.2 | | Type II | chr4:72039073-72044540 | 1 | 0 | Called 542/584. fragment 314/542, noncarrier call 228/542 | None |
| 4q32.1 | HERV-K124 | Type II | chr4:164009164-164011582 | 1 | 0 | Called 584/584. fragment 584/584 | None |
| 4q32.3 | K5, ERVK-12 | Type I | chr4:168343347-168350576 | 1 | 0 | Called 584/584. fragment 584/584 | None |
| 4q35.2 | | Type II | chr4:190106259-190113546 | 2 | 0 | Called 479/584. multi-copy 1/479, fragment 206/479, noncarrier call 272/479 | None |
| 5p12 | | Type I | chr5:46253263-46261226 | 2 | 1 | Called 582/584. provirus 579/582, multi-copy 1/582, noncarrier call 2/582 | None |
| 5p13.3 | HERV-K104, K50d | Type II | chr5:30600649-30610094 | 1 | 1 | Called 584/584. provirus 576/584, solo-ltr 8/584 | Env, Rec |
| 5q11.2 | | Type II | chr5:60282082-60283050 | 1 | 0 | Called 584/584. solo-ltr 584/584 | None |
| 5q33.2 | K18b | Type II | chr5:155169928-155178636 | 1 | 1 | Called 583/584. provirus 578/583, noncarrier call 5/583 | Rec |
| 5q33.3 | HERV-K107, K10, K(C5), ERVK-10 | Type I | chr5:157176701-157185880 | 3 | 3 | Called 583/584. provirus 552/583, multi-copy 4/583, solo-ltr 27/583 | Gag, Gag–Pro, Gag–Pro–Pol, Np9 |
| 6p21.1 | K(OLDAL035587), KOLD35587 | Type II | chr6:42722387-42732345 | 1 | 1 | Called 584/584. provirus 584/584 | None |

| Catalog label | Former name(s) | Type | T2T-CHM13 coordinates | Maximum retained copies per haplotype | Maximum array size | Structural observations | Selected ORF combinations |
|---|---|---|---|---|---|---|---|
| 6p22.1 | K(OLDAL121932), K69, K20 | Type II | chr6:28553904-28564272 | 1 | 1 | Called 584/584. provirus 584/584 | None |
| 6q11.1 | | Type II | chr6:61822916-61828905 | 1 | 0 | Called 584/584. fragment 448/584, noncarrier call 136/584 | None |
| 6q14.1 | HERV-K109, K(C6), ERVK-9 | Type II | chr6:78894317-78903741 | 3 | 3 | Called 582/584. provirus 482/582, multi-copy 4/582, solo-ltr 93/582, noncarrier call 3/582 | Gag, Gag–Pro, Env, Rec |
| 6q25.1 | | Type II | chr6:152060170-152062995 | 1 | 0 | Called 584/584. solo-ltr 1/584, fragment 583/584 | None |
| 7p22.1 | HERV-K108, K(HLM-2.HOM), K(C7), ERVK-6 | Type II | chr7:4699540-4709011 | 6 | 6 | Called 583/584. provirus 334/583, multi-copy 247/583, solo-ltr 2/583 | Gag, Gag–Pro, Gag–Pro–Pol, Env, Rec |
| 7q11.21 | | Type II | chr7:67226777-67229896 | 2 | 0 | Called 584/584. multi-copy 1/584, solo-ltr 2/584, fragment 581/584 | None |
| 7q22.2 | HERV-K126, ERVK-14 | Type II | chr7:106062272-106067170 | 1 | 0 | Called 584/584. fragment 584/584 | None |
| 7q34 | K(OLDAC004979), ERVK-15 | Type II | chr7:143066952-143071964 | 1 | 0 | Called 584/584. fragment 584/584 | None |
| 8p23.1 copy group | | Type II | Not uniquely assigned here | 1 | | Unlocalized records in 6 haplotypes, provirus 6. Not a single-locus frequency. | Rec |
| 8p23.1a | HERV-K115, ERVK-8 | Type II | chr8:6881573-6883482 | 1 | 1 | Called 515/584. provirus 42/515, noncarrier call 473/515 | Env, Rec |
| 8p23.1b | K27 | Type II | chr8:6940112-6942025 | 1 | 1 | Called 531/584. provirus 284/531, noncarrier call 247/531 | Rec |
| 8p23.1c | | Type II | chr8:11536938-11546458 | 1 | 1 | Called 504/584. provirus 252/504, noncarrier call 252/504 | Rec |
| 8p23.1d | KOLD130352 | Type II | chr8:7409812-7411728 | 2 | 1 | Called 501/584. provirus 281/501, multi-copy 5/501, noncarrier call 215/501 | Rec |

| Catalog label | Former name(s) | Type | T2T-CHM13 coordinates | Maximum retained copies per haplotype | Maximum array size | Structural observations | Selected ORF combinations |
|---|---|---|---|---|---|---|---|
| 8p23.1e | | Type II | chr8:7508401-7517950 | 2 | 1 | Called 550/584. provirus 328/550, multi-copy 1/550, noncarrier call 221/550 | Rec |
| 8q11.21 | | Type II | chr8:46642148-46650152 | 1 | 0 | Called 584/584. fragment 584/584 | None |
| 8q11.23 | novel (this study) | Type I | chr8:54409084-54410051 | 1 | 1 | Called 584/584. provirus 1/584, solo-ltr 583/584 | Env* |
| 8q24.3a | HERV-K127 | Type II | chr8:140580372-140583458 | 1 | 0 | Called 584/584. fragment 584/584 | None |
| 8q24.3c | | Type II | chr8:146196392-146203288 | 2 | 0 | Called 582/584. multi-copy 1/582, fragment 577/582, noncarrier call 4/582 | None |
| 9q34.11 | K31, DE7, ERVK16 | Type II | chr9:141055003-141062224 | 1 | 0 | Called 582/584. fragment 581/582, noncarrier call 1/582 | None |
| 9q34.3 | K30 | Type II | chr9:149013195-149022657 | 1 | 1 | Called 582/584. provirus 370/582, fragment 210/582, noncarrier call 2/582 | None |
| Xq11.1 | | Type II | chrX:61161158-61163663 | 1 | 0 | Called 430/432. solo-ltr 91/430, fragment 339/430 | None |
| Xq12 | | Type II | chrX:64892059-64894111 | 1 | 0 | Called 431/432. fragment 431/431 | None |
| Xq21.33 | De9 | Type II | chrX:92796779-92806254 | 1 | 1 | Called 431/432. provirus 6/431, noncarrier call 425/431 | Gag, Gag–Pro, Gag–Pro–Pol, Env, Rec |
| Xq28a | K63 | Type II | chrX:152825089-152832431 | 1 | 0 | Called 427/432. fragment 210/427, noncarrier call 217/427 | None |
| Xq28b | K63 | Type II | chrX:152849562-152852185 | 2 | 0 | Called 427/432. multi-copy 1/427, fragment 281/427, noncarrier call 145/427 | None |
| Yp11.2 | | Type II | chrY:6610727-6617670 | 1 | 0 | Called 144/144. fragment 124/144, noncarrier call 20/144 | None |

| Catalog label | Former name(s) | Type | T2T-CHM13 coordinates | Maximum retained copies per haplotype | Maximum array size | Structural observations | Selected ORF combinations |
|---|---|---|---|---|---|---|---|
| Yq11.23 copy group | | Type II | Not uniquely assigned here | 1 | | Unlocalized records in 6 haplotypes, solo-ltr 6. Not a single-locus frequency. | None |
| Yq11.23a | | Type II | chrY:24607688-24610887 | 2 | 0 | Called 114/144. multi-copy 2/114, solo-ltr 32/114, noncarrier call 80/114 | None |
| Yq11.23b | | Type II | chrY:26227634-26230834 | 3 | 0 | Called 114/144. multi-copy 6/114, solo-ltr 88/114, noncarrier call 20/114 | None |
| Telomeric Type I | 13p13, 15p13b | Type I | chr13:5381952-5388076 / chr15:2092086-2101275 | 3 | 1 | Unlocalized records in 180 haplotypes, solo-ltr 30, fragment 105, provirus 12, multiple records 33. Not a single-locus frequency. | Np9 |
| Telomeric Type II | 15p13a, 21p13, 22p13 | Type II | chr15:12216-19499 / chr21:2699539-2710506 / chr22:10726-18005 | 4 | 0 | Unlocalized records in 277 haplotypes, fragment 191, multiple records 86. Not a single-locus frequency. | None |

**Table S2. Review of candidate additional copies**

Per-copy read-depth evidence and classification. Table_S4_NucFreq_region_summary.tsv gives regional mixed-base counts. Table_S4_raw_read_source_index.tsv lists the HPRC raw-read files, available SRA accessions, and BioProjects indexed for the 93 donors represented in the validation tables. Tab-delimited files are in HML2_Supplementary_Data.zip.

**Table S3. ORF-associated variant recovery and structural-variant diagnostics**

Native short-read genotype support for 564 ORF-associated variant alleles identified in matched long-read sequences. The supporting files include the donor–locus–variant observations, complete quality and unresolved-state classifications, per-gene and per-locus summaries, sequence and coordinate provenance, native genotype slices, and reproduction scripts. Separate Illumina ensemble structural-variant diagnostics are provided in Figure S1. These results do not reconstruct whole ORFs from a variant-only VCF.

**Table S4. Solo-LTR diversity and expected nucleotide differences**

Part A gives sequence counts, common callable LTR bases, nucleotide diversity, exact difference and denominator counts, and mean differences per sequence pair at each comparison locus. Part B gives expected differences across 968 bp under the two pairwise clocks as a function of sequence-divergence time. Source sequence paths, primary alignments, and excluded loci accompany the table.

**Table S5. Structural-state source table**

Structural-state counts and calls/total chromosome counts at each locus. Record-level structural and source evidence shown. Copy groups are reported separately. Provided as tab-delimited files in HML2_Supplementary_Data.zip.

**Table S6. Array copy-number distributions**

Resolved 7p22.1 and 1p31.1b copy-number distributions and haplotype calls. At 7p22.1, two solo-LTR haplotypes have zero proviral copies. HG00658 pat remains uncalled and is excluded from the called-haplotype denominator of 583. All 584 sampled haplotypes are retained in the call table in HML2_Supplementary_Data.zip.

**Table S7. Tandem arrays and solo-LTR alleles**

Counts use the retained population array table and haplotypes with a locus-level structural call. Frequencies are shown as numerator/denominator. The solo/tandem ratio is the number of solo-LTR observations divided by the number of tandem-array observations.

| Locus | Type | Maximum array size | Tandem-allele frequency | Solo-LTR frequency | Single-provirus frequency | Solo/tandem ratio |
|---|---|---|---|---|---|---|
| 7p22.1 | Type II | 6 | 247/583 (42.4%) | 2/583 (0.3%) | 334/583 (57.3%) | 0.01 |
| 12q24.33 | Type II | 4 | 13/571 (2.3%) | 20/571 (3.5%) | 0/571 (0.0%) | 1.54 |
| 1p31.1b | Type I | 4 | 9/584 (1.5%) | 469/584 (80.3%) | 0/584 (0.0%) | 52.11 |
| 5q33.3 | Type I | 3 | 4/583 (0.7%) | 27/583 (4.6%) | 552/583 (94.7%) | 6.75 |
| 6q14.1 | Type II | 3 | 4/582 (0.7%) | 93/582 (16.0%) | 482/582 (82.8%) | 23.25 |
| 11q22.1 | Type II | 2 | 3/584 (0.5%) | 304/584 (52.1%) | 213/584 (36.5%) | 101.33 |
| 3q13.2 | Type I | 2 | 3/584 (0.5%) | 56/584 (9.6%) | 525/584 (89.9%) | 18.67 |
| 3q27.2 | Type I | 2 | 2/584 (0.3%) | 21/584 (3.6%) | 561/584 (96.1%) | 10.50 |
| 10p12.1 | Type I | 2 | 1/584 (0.2%) | 25/584 (4.3%) | 558/584 (95.5%) | 25.00 |
| 11q23.3 | Type I | 2 | 1/584 (0.2%) | 9/584 (1.5%) | 574/584 (98.3%) | 9.00 |
| 12q13.2 | Type I | 2 | 1/584 (0.2%) | 130/584 (22.3%) | 258/584 (44.2%) | 130.00 |
| 12q14.1 | Type II | 2 | 1/582 (0.2%) | 307/582 (52.7%) | 274/582 (47.1%) | 307.00 |
| 14q11.2 | Type II | 2 | 1/584 (0.2%) | 0/584 (0.0%) | 0/584 (0.0%) | 0.00 |

**Table S8. Nucleotide variation within tandem arrays**

Copy provenance and extracted sequences are linked by the retained copy identifier. Pairwise tables report jointly called length, substitution count, nucleotide divergence, transitions, internal gap tracts, and terminal differences in extent. Array and locus summaries retain their respective denominators. Variable-site and indel tables show observed patterns in copy order. The duplication-order table lists all closest-pair ties for arrays containing at least three copies. The site-frequency table uses the number of arrays callable at each site as its denominator. The conditional-clock table applies the stated neutral rate sensitivity separately to each pair. It includes exact count uncertainty and one-sided upper limits for identical sequences.

**Table S9. Phylogenetic and host-flank relationships among HML-2 loci**
Regional nucleotide comparisons and retained reference-alignment records supporting Figure 3C are supplied in Supplementary_Data/Table_S14/4q_acrocentric_relationship/. Paired-LTR measurements, population sequence comparisons, reference-flank alignments, tree files, and bootstrap results for Figure S2 are supplied in Supplementary_Data/Table_S14/telomeric_phylogeny/. Figure 3 alignments, bootstrap support, nearest-neighbor frequencies, average pairwise nucleotide differences, and sequence denominators are supplied in Supplementary_Data/Table_S14/phylogeny_review/.

**Table S10. Combined ORF-screen annotations**
Per-locus evaluable and passing ORF-annotation counts. Provided as tab-delimited files in HML2_Supplementary_Data.zip.

**Table S11. Type-I cassette comparisons and complete alignment data**

The source files contain the 45 human representative identifiers and subfamily assignments, seven-window callability, pairwise nucleotide-difference numerators and denominators, group summaries, similarity rankings, nearest Type-II representatives, human per-site base frequencies and consensus sequences, and the 83-cluster nonhuman primate manifest with species and type calls. The supporting data include the full aligned FASTA files. The input-provenance table records the exact retained source files and their SHA-256 digests. The numerical windows and FASTA coordinates are recorded as zero-based half-open intervals in machine-readable tables. Figure S13 coordinates are one-based and inclusive. The TypeI_deletion_models directory supplies the deletion-class counts, ancestral-insertion assignments, corrected source-model outputs, and reproduction scripts.

**Table S12. Direct type-state audit**
Direct cassette calls and source evidence at 20 Type-I loci, counted once per donor, haplotype, and locus. Solo-LTRs, noncarrier records, and fragments without the cassette do not enter the cassette-callable denominator. Provided in HML2_Supplementary_Data.zip.

**Table S13. Recurrent conversion and within-locus Type-I/II polymorphism**
Probabilities of observing no mixed Type-I/II samples across 54 conversion-rate, population-size, and age scenarios. Files include locus identities, sample sizes, age bounds, numerical checks, and analysis code. Files are in HML2_Supplementary_Data.zip.

**Table S14. Archaic reads spanning the 8q11.23 host–LTR junctions**
Junction-read counts from four archaic genomes using 10-, 20-, and 30-bp flanking anchors. Reference alignments and chimpanzee/gorilla empty-site evidence are included. Files are in HML2_Supplementary_Data.zip.

**Table S15. Functional association results**
Tables S15a–d contain the 237-model screen, 51-model burden analysis, nonoverlapping-cohort SLC44A5 estimates, and 40 anti-CD20 models. Tables S15e–h provide the complete MAGE discovery tests, candidate eligibility and aliases, GEUVADIS SLC44A5 follow-up tests, and MAGE HC3 sensitivity models. Table S15a identifies the 174 nonmissing P values used in its correction. Table S15g identifies the 30 tests used in the GEUVADIS follow-up correction. Files are in HML2_Supplementary_Data.zip.

**Table S16. 1q22 Gag copy number**
Artifact-filtered 1q22 Gag observations. Provided as tab-delimited files in HML2_Supplementary_Data.zip.

**Table S17. ORF annotations disrupted in GRCh38 but passing the combined screen in other alleles**

Counts are donors with at least one copy meeting the combined Intact and Intact_FS_End screen among 292 donors. Gag–Pro requires both annotations on the same provirus. At Type-I loci, env refers to the remaining frame for a theoretical N-terminally truncated product whose translation has not been demonstrated.

| **Locus** | **Former name** | ORF annotation | GRCh38 ORF-disrupting change | Donors with a passing ORF annotation |
|---|---|---|---|---|
| 7p22.1 | HERV-K108 | Gag | nonsense | 269/292 (92.1%) |
| 7p22.1 | HERV-K108 | Gag-Pro (Pro) | frameshift | 269/292 (92.1%) |
| 1q22 | HERV-K102 | Gag | nonsense | 168/292 (57.5%) |
| 1q22 | HERV-K102 | Gag-Pro (Pro) | nonsense | 168/292 (57.5%) |
| 5p13.3 | HERV-K104 | Env | nonsense | 34/292 (11.6%) |
