## Supplementary material for "Structural polymorphism and population-variable coding capacity of HERV-K(HML-2) in human pangenomes": coriell_GM24631.html

Coriell Institute for Medical Research

- Request a Quote for Custom Services
- Donate
- Login
- View Cart

Samples
OR

Website

Sample Catalog |
Custom Services |
Core Facilities |
Genomic Data Search

Navigation Header

- Biobank
  - NRGR
  - NIGMS
  - NINDS
  - NIA
  - NHGRI
  - NEI
  - Allen Cell Collection
  - Rett Syndrome iPSC Collection
  - Autism Research Resource
  - HD Community Biorepository
  - CDC Cell and DNA
  - J. Craig Venter Institute
  - Orphan Disease Center Collection
  - All Biobanks
- Research
  - Overview
  - Meet Our Scientists
    - Our Faculty
    - Our Scientific Staff
  - Camden Cancer Research Center
  - Epigenetic Therapies SPORE
  - Core Facilities
  - Epigenomics
  - Camden Opioid Research Initiative (CORI)
  - The Issa Lab
  - The Jian Huang Lab
  - The Luke Chen Lab
    - The Lab
    - The Team
    - Publications
  - The Scheinfeldt Lab
  - The Shumei Song Lab
  - The Nora Engel Lab
    - The Lab
    - The Team
    - Publications
  - Publications
- Services
  - Overview
  - Biobanking Services
    - Core Services
    - Project Management
    - Research Support Services
    - Sample Cataloging
    - Sample Collection Kits
    - Sample Data Management
    - Sample Distribution
    - Sample Management
    - Sample Procurement
    - Sample Storage
  - Bioinformatics and Biostatistics Services
  - Cellular and Molecular Services
    - Biomarker Research Solutions
    - Cell Culture
    - Nucleic Acid Isolation and Quality Control
  - Clinical Trial Support
    - Overview
    - Sample Collection
    - Data Management
    - Sample Processing and QC
    - Storage and Distribution
    - Biomarker Services
    - Data Analaysis
  - Core Facilties
    - Overview
    - Animal and Xenograft
    - Bioinformatics and Biostatistics
    - Cell Imaging
    - CRISPR Gene Engineering
    - Flow Cytometry and Cell Sorting
    - Genomics and Epigenomics
    - iPSC - Induced Pluripotent Stem Cells
    - Organoids
  - Coriell Marketplace
  - Genomic, Epigenomic and Multiomics Services
  - Stem Cells and iPSC Services
    - Core Services
    - Reprogramming
    - Characterization and Quality Control
    - Differentiated Cell Lines
    - iPSC-Derived Organoids
    - iPSC Expansion
    - iPSC Gene Editing
- Ordering
  - Stem Cells
  - Cell Lines
  - DNA and RNA
  - Featured Products
    - FFPE
    - HMW DNA
  - Genomic Data Search
  - Search by Catalog ID
  - Help
    - Create Account
    - Order Online
    - Ordering FAQ
    - FAQs/Culture Instructions
    - Reference Materials
      - Biobanks
      - NIGMS Repository
      - NHGRI Repository
      - NINDS Repository
      - NIA Repository
      - NIST
      - GeT-RM
    - Secondary Distribution Policies
    - MTA Assurance Form
    - Shipment Policy
    - Contact Customer Service
- About Us
  - About Coriell
  - Meet Our Team
  - Meet Our Board
  - Education
    - Science Fair
    - Outreach
    - College Internships
  - Press Room
    - Press Releases
    - Coriell Blog
    - Annual Report
  - Careers
    - Working at Coriell
    - Verifications of Employment
  - Giving
    - Donate
    - Giving FAQ
  - Contact Us
  - Notices
    - Legal Notice
    - IBC Minutes

- Login

  View Cart

search submit

GM24631

**LCL** from **B-Lymphocyte**

##### Description:

##### PERSONAL GENOME PROJECT

##### Affected:

##### Unknown

##### Sex:

##### Male

##### Age:

##### 33 YR (At Sampling)

Sample Description

- **Overview**
- **Characterizations**
- **Phenotypic Data**
- **Publications**

- **Culture Protocols**

### Overview

|  |  |
| --- | --- |
| **Repository** | NIGMS Human Genetic Cell Repository |
| **Subcollection** | Apparently Healthy Collection PIGI Consented Sample |
| **Biopsy Source** | Peripheral vein |
| **Cell Type** | B-Lymphocyte |
| **Tissue Type** | Blood |
| **Transformant** | Epstein-Barr Virus |
| **Sample Source** | LCL from B-Lymphocyte |
| **Race** | Asian |
| **Ethnicity** | Chinese |
| **Country of Origin** | USA |
| **Family Member** | 1 |
| **Family History** | N |
| **Relation to Proband** | proband |
| **Species** | Homo sapiens |
| **Common Name** | Human |
| **Remarks** | Participant (hu91BD69) in the Personal Genome Project: http://www.personalgenomes.org history of: eczema; lactose intolerance; nearsightedness; same subject as GM26107 (stem cell); father is GM24694 (Lymph); mother is GM24695 (Lymph). |

### Characterizations

|  |  |
| --- | --- |
| **IDENTIFICATION OF SPECIES OF ORIGIN** | Species of Origin Confirmed by LINE assay |

### Phenotypic Data

|  |  |
| --- | --- |
| **Remarks** | Participant (hu91BD69) in the Personal Genome Project: http://www.personalgenomes.org history of: eczema; lactose intolerance; nearsightedness; same subject as GM26107 (stem cell); father is GM24694 (Lymph); mother is GM24695 (Lymph). |

### Publications

**Cai K, Li S, Pan M, Lu H, Wang L, Fang S, Gou L, Tang J, Kong Y, Zhao L, Ren Y**, Comparative assessment of the Sikun 2000 sequencing platform for whole genome sequencing Scientific reports15:19070 2025

**PubMed ID: 40447879**

**Emiliani FE, Ismail AAO, Hughes EG, Tsongalis GJ, Zanazzi GJ, Lin CC**, Nanopore-based random genomic sampling for intraoperative molecular diagnosis Genome medicine17:6 2025

**PubMed ID: 39833913**

**Longo GMC, Sayols S, Kotini AG, Heinen S, Möckel MM, Beli P, Roukos V**, Linking CRISPR-Cas9 double-strand break profiles to gene editing precision with BreakTag Nature biotechnology17:6 2025

**PubMed ID: 38740992**

**Mitchell R, Peck M, Gorden E, Just R**, MixDeR: A SNP mixture deconvolution workflow for forensic genetic genealogy Forensic science international Genetics76:103224 2025

**PubMed ID: 39862579**

**Verner EL, Jackson JB, Maddox C, Valkenburg KC, White JR, Occean J, Morris L, Karandikar A, Gerding KMR, Sausen M, Koohestani F, Severson EA, Jensen TJ, Caveney BJ, Eisenberg M, Ramkissoon SH, Greer AE**, Analytical Validation of the Labcorp Plasma Complete Test, a Cell-Free DNA Comprehensive Genomic Profiling Tool for Precision Oncology The Journal of molecular diagnostics : JMD27:216-231 2025

**PubMed ID: 39818317**

**Pedroza Matute S, Turvey K, Iyavoo S**, Advancing human genotyping: The Infinium HTS iSelect Custom microarray panel (Rita) development study Forensic science international Genetics71:103049 2024

**PubMed ID: 38653142**

**Smullen M, Olson MN, Reichert JM, Dawes P, Murray LF, Baer CE, Wang Q, Readhead B, Church GM, Lim ET, Chan Y**, Reliable multiplex generation of pooled induced pluripotent stem cells Cell reports methods3:100570 2023

**PubMed ID: 37751688**

**Verner EL, Jackson JB, Severson E, Valkenburg KC, Greer AE, Riley DR, Sausen M, Maddox C, McGregor PM, Karandikar A, Hastings SB, Previs RA, Reddy VP, Jensen TJ, Ramkissoon SH**, Validation of the Labcorp Plasma Focus Test to Facilitate Precision Oncology Through Cell-Free DNA Genomic Profiling of Solid Tumors The Journal of molecular diagnostics : JMD3:100570 2023

**PubMed ID: 37068734**

**Steiert TA, Fuß J, Juzenas S, Wittig M, Hoeppner MP, Vollstedt M, Varkalaite G, ElAbd H, Brockmann C, Görg S, Gassner C, Forster M, Franke A**, High-throughput method for the hybridisation-based targeted enrichment of long genomic fragments for PacBio third-generation sequencing NAR genomics and bioinformatics4:lqac051 2022

**PubMed ID: 35855323**

**Foox J, Nordlund J, Lalancette C, Gong T, Lacey M, Lent S, Langhorst BW, Ponnaluri VKC, Williams L, Padmanabhan KR, Cavalcante R, Lundmark A, Butler D, Mozsary C, Gurvitch J, Greally JM, Suzuki M, Menor M, Nasu M, Alonso A, Sheridan C, Scherer A, Bruinsma S, Golda G, Muszynska A, Labaj PP, Campbell MA, Wos F, Raine A, Liljedahl U, Axelsson T, Wang C, Chen Z, Yang Z, Li J, Yang X, Wang H, Melnick A, Guo S, Blume A, Franke V, Ibanez de Caceres I, Rodriguez-Antolin C, Rosas R, Davis JW, Ishii J, Megherbi DB, Xiao W, Liao W, Xu J, Hong H, Ning B, Tong W, Akalin A, Wang Y, Deng Y, Mason CE**, The SEQC2 epigenomics quality control (EpiQC) study Genome biology22:332 2021

**PubMed ID: 34872606**

**Ralf A, Zandstra D, Weiler N, van Ijcken WFJ, Sijen T, Kayser M**, RMplex: An efficient method for analyzing 30 Y-STRs with high mutation rates Forensic science international Genetics55:102595 2021

**PubMed ID: 34543845**

**Rehder C, Bean LJH, Bick D, Chao E, Chung W, Das S, O'Daniel J, Rehm H, Shashi V, Vincent LM, ACMG Laboratory Quality Assurance Committee LM**, Next-generation sequencing for constitutional variants in the clinical laboratory, 2021 revision: a technical standard of the American College of Medical Genetics and Genomics (ACMG) Genetics in medicine : official journal of the American College of Medical Genetics23:1399-1415 2021

**PubMed ID: 33927380**

**Hynst J, Navrkalova V, Pal K, Pospisilova S**, Bioinformatic strategies for the analysis of genomic aberrations detected by targeted NGS panels with clinical application PeerJ9:e10897 2020

**PubMed ID: 33850640**

**Min YK, Lee YK, Nam SH, Kim JK, Park KS, Kim JW**, Quantitative and Qualitative QC of Next-Generation Sequencing for Detecting Somatic Variants: An Example of Detecting Clonal Hematopoiesis of Indeterminate Potential Clinical chemistry9:e10897 2020

**PubMed ID: 32395759**

**Atkins A, Gupta P, Zhang BM, Tsai WS, Lucas J, Javey M, Vora A, Mei R**, Detection of Circulating Tumor DNA with a Single-Molecule Sequencing Analysis Validated for Targeted and Immunotherapy Selection Molecular diagnosis & therapy9:e10897 2019

**PubMed ID: 31209714**

**Fujiki R, Ikeda M, Ohara O**, Short DNA Probes Developed for Sample Tracking and Quality Assurance in Gene Panel Testing The Journal of molecular diagnostics : JMD9:e10897 2019

**PubMed ID: 31445212**

**Lazzarotto CR, Malinin NL, Li Y, Zhang R, Yang Y, Lee G, Cowley E, He Y, Lan X, Jividen K, Katta V, Kolmakova NG, Petersen CT, Qi Q, Strelcov E, Maragh S, Krenciute G, Ma J, Cheng Y, Tsai SQ**, CHANGE-seq reveals genetic and epigenetic effects on CRISPR-Cas9 genome-wide activity Nature biotechnology9:e10897 2019

**PubMed ID: 32541958**

**Vegesna R, Tomaszkiewicz M, Medvedev P, Makova KD**, Dosage regulation, and variation in gene expression and copy number of human Y chromosome ampliconic genes PLoS genetics15:e1008369 2019

**PubMed ID: 31525193**

**Cleveland MH1, Zook JM2, Salit M3, Vallone PM2.**, Determining Performance Metrics for Targeted Next-Generation Sequencing Panels Using Reference Materials Journal of Molecular Diagnostics18:1525-1578 2018

**PubMed ID: 29959024**

**Lincoln SE, Truty R, Lin CF, Zook JM, Paul J, Ramey VH, Salit M, Rehm HL, Nussbaum RL, Lebo MS**, A Rigorous Interlaboratory Examination of the Need to Confirm Next-Generation Sequencing-Detected Variants with an Orthogonal Method in Clinical Genetic Testing The Journal of molecular diagnostics : JMD18:1525-1578 2018

**PubMed ID: 30610921**

**Soukupova J, Zemankova P, Lhotova K, Janatova M, Borecka M, Stolarova L, Lhota F, Foretova L, Machackova E, Stranecky V, Tavandzis S, Kleiblova P, Vocka M, Hartmannova H, Hodanova K, Kmoch S, Kleibl Z**, Validation of CZECANCA (CZEch CAncer paNel for Clinical Application) for targeted NGS-based analysis of hereditary cancer syndromes PloS one13:e0195761 2017

**PubMed ID: 29649263**

**Mao Q, Ciotlos S, Zhang RY, Ball MP, Chin R, Carnevali P, Barua N, Nguyen S, Agarwal MR, Clegg T, Connelly A, Vandewege W, Zaranek AW, Estep PW, Church GM, Drmanac R, Peters BA.**, The whole genome sequences and experimentally phased haplotypes of over 100 personal genomes. Gigascience.5(1):42 2016

**PubMed ID: 27724973**

**Church GM**, The personal genome project. Mol Syst Biol.1, 2005.0030:42 2005

**PubMed ID: 16729065**

### Culture Protocols

|  |  |
| --- | --- |
| **Split Ratio** | 1:3 |
| **Temperature** | 37 C |
| **Percent CO2** | 5% |
| **Percent O2** | AMBIENT |
| **Medium** | Roswell Park Memorial Institute Medium 1640 with 2mM L-glutamine or equivalent |
| **Serum** | 15% fetal bovine serum Not Inactivated |
| **Substrate** | None specified |
| **Subcultivation Method** | dilution - add fresh medium |
| **Supplement** | - |

Pricing

International/Commercial/For-profit:

$448.00USD

U.S. Academic/Non-profit/Government:

$262.00USD

Add to Cart

How to Order

- Ordering Instructions
- MTA / Assurance Form
- Statement of Research Intent Form

Related Products

Same Subject

- NA24631 - DNA
- HM24631 - High Molecular Weight DNA
- GM26107 - Stem cell

Same Family

- 3150

Miscellaneous

- DNA on Demand
- Custom Services

Our mission is to prevent and cure disease through biomedical research.

CONTACT US

Catalog Inquiries

Mailing Address

Coriell Institute for Medical Research  
403 Haddon Avenue Camden, NJ 08103, USA  

Subscribe to our newsletter here

Ⓒ 2026 Coriell Institute. All rights reserved.

- Facebook
- Linkedin

- Youtube
