## Supplementary material for "Structural polymorphism and population-variable coding capacity of HERV-K(HML-2) in human pangenomes": coriell_HG02109.html

- Login


  View Cart


search submit

HG02109

**LCL** from **B-Lymphocyte**

##### Description:

##### AFRICAN CARIBBEAN IN BARBADOS

##### Affected:

##### No Data

##### Sex:

##### Female

##### Age:

##### No Data

Sample Description

- **Overview**
- **Phenotypic Data**


- **Culture Protocols**


### Overview

|  |  |
| --- | --- |
| **Repository** | NHGRI Sample Repository for Human Genetic Research |
| **Subcollection** | NHGRI Sample Repository for Human Genetic Research |
| **Biopsy Source** | Peripheral vein |
| **Cell Type** | B-Lymphocyte |
| **Tissue Type** | Blood |
| **Transformant** | Epstein-Barr Virus |
| **Sample Source** | LCL from B-Lymphocyte |
| **Country of Origin** | BARBADOS |
| **Family Member** | 3 |
| **Family History** | N |
| **Relation to Proband** | child |
| **Species** | Homo sapiens |
| **Common Name** | Human |
| **Remarks** | All four grandparents consider themselves African-Caribbean and were born in Barbados |

### Phenotypic Data

|  |  |
| --- | --- |
| **Remarks** | All four grandparents consider themselves African-Caribbean and were born in Barbados |

### Culture Protocols

|  |  |
| --- | --- |
| **Split Ratio** | 1:2 |
| **Temperature** | 37 C |
| **Percent CO2** | 5% |
| **Percent O2** | AMBIENT |
| **Medium** | Roswell Park Memorial Institute Medium 1640 with 2mM L-glutamine or equivalent |
| **Serum** | 15% fetal bovine serum Not Inactivated |
| **Substrate** | None specified |
| **Subcultivation Method** | dilution - add fresh medium |
| **Supplement** | - |

Pricing

International/Commercial/For-profit:

$448.00USD

U.S. Academic/Non-profit/Government:

$225.00USD

Add to Cart

How to Order

- Ordering Instructions
- MTA / Assurance Form
- Statement of Research Intent Form

Related Products

Same Subject

- HG02109 - DNA

Same Family

- BB18

Miscellaneous

- Custom Services

Our mission is to prevent and cure disease through biomedical research.

CONTACT US

Catalog Inquiries

Mailing Address

Coriell Institute for Medical Research  
403 Haddon Avenue Camden, NJ 08103, USA  

Subscribe to our newsletter here

Ⓒ 2026 Coriell Institute. All rights reserved.

- Facebook
- Linkedin

- Youtube
