## Supplementary material for "Structural polymorphism and population-variable coding capacity of HERV-K(HML-2) in human pangenomes": coriell_HG06807.html

- Login


  View Cart


search submit

HG06807

**LCL** from **B-Lymphocyte**

##### Description:

##### AFRICAN AMERICANS LIVING IN ST. LOUIS, MISSOURI

##### Affected:

##### No Data

##### Sex:

##### Female

##### Age:

##### No Data

Sample Description

- **Overview**
- **Phenotypic Data**


- **Culture Protocols**


### Overview

|  |  |
| --- | --- |
| **Repository** | NHGRI Sample Repository for Human Genetic Research |
| **Subcollection** | NHGRI Sample Repository for Human Genetic Research |
| **Cell Type** | B-Lymphocyte |
| **Transformant** | Epstein-Barr Virus |
| **Sample Source** | LCL from B-Lymphocyte |
| **Country of Origin** | USA |
| **Family Member** | 2 |
| **Family History** | N |
| **Relation to Proband** | mother |
| **ISCN** | 46,XX[20] |
| **Species** | Homo sapiens |
| **Common Name** | Human |
| **Remarks** | These cell lines and DNA samples were prepared from peripheral blood samples collected from people living in the St. Louis, MO metropolitan area who self-identified as African American. |

### Phenotypic Data

|  |  |
| --- | --- |
| **Remarks** | These cell lines and DNA samples were prepared from peripheral blood samples collected from people living in the St. Louis, MO metropolitan area who self-identified as African American. |

### Culture Protocols

|  |  |
| --- | --- |
| **Split Ratio** | 1:4 |
| **Temperature** | 37 C |
| **Percent CO2** | 5% |
| **Percent O2** | AMBIENT |
| **Medium** | Roswell Park Memorial Institute Medium 1640 with 2mM L-glutamine or equivalent |
| **Serum** | 15% fetal bovine serum Not Inactivated |
| **Substrate** | None specified |
| **Subcultivation Method** | dilution - add fresh medium |
| **Supplement** | - |

Pricing

International/Commercial/For-profit:

$448.00USD

U.S. Academic/Non-profit/Government:

$225.00USD

Add to Cart

How to Order

- Ordering Instructions
- MTA / Assurance Form
- Statement of Research Intent Form

Related Products

Same Subject

- HG06807 - DNA
- HG06807 - Stem cell

Same Family

- 3559

Miscellaneous

- Custom Services

Our mission is to prevent and cure disease through biomedical research.

CONTACT US

Catalog Inquiries

Mailing Address

Coriell Institute for Medical Research  
403 Haddon Avenue Camden, NJ 08103, USA  

Subscribe to our newsletter here

Ⓒ 2026 Coriell Institute. All rights reserved.

- Facebook
- Linkedin

- Youtube
