## Supplementary material for "Structural polymorphism and population-variable coding capacity of HERV-K(HML-2) in human pangenomes": hprc_release2_sample_README.md

# Samples Included In Release 2

## Overview
Release 2 includes 234 samples (466 haplotypes). The samples can be broken down into three categories:
* **216 HPRC samples**
  * Trio phasing was used when available, otherwise phasing was performed with Hi-C:
    * 126 samples phased with trio information (parental Illumina data)
    * 90 samples were phased with Hi-C
* **14 HPP samples**: sequencing and assembly were led in other projects but which are in collaboration with the HPRC
  * All HPP samples were phased with Hi-C
* **4 Extramural samples**: sequenced and assembled by other projects and which are used by the HPRC as reference-level assemblies. This includes:
  * HG06807
  * HG002
  * GRCh38
  * CHM13

The majority of samples in Release 2 are from the 1000G project and are EBV transformed cell lines sourced from Coriell. Being part of the 1000G project means that the samples are openly consented and include no clinical information. A few samples were sequenced or are from other collections/consortiums. These samples are as openly consented as the 1000G samples, but their sequencing data may come from other cell types.

# Important Nomenclature
## HPRC and HPP
**HPRC samples** are samples that were sequenced and assembled by the HPRC in it's production efforts. In the context of Release 2, samples designated HPRC may have been designated "HPRC PLUS" (see below).

**HPP samples** are samples where sequencing and assembly were led in other projects but which are in collaboration with the HPRC. These samples were not selected by the Population and Sampling Working group, but rather by the projects themselves.

## HPRC and HPRC PLUS
**HPRC samples** were initially selected by the HPRC's Population and Sampling Working Group as the set of samples from the 1000G cohort which provides an efficient sampling of global diversity. The 1000G samples were subset to samples which:
* Have low passage cells available
* Were child of a trio in order to allow for assembly phasing. 
    * Later sampling, in anticipation of Hi-C phasing, broadened to samples that are not children in trios.

Samples selected were karyotyped and screened then sequenced by the HPRC's Production Working Group with PacBio HiFi, PacBio Kinnex, ONT, and Illumina Hi-C technologies. The HPRC uses Illumina sequencing data from the NYGC's High Coverage (30X) sequencing of the 1000G samples.

**HPRC PLUS Samples** were samples who were not selected by the HPRC's Population and Sampling Working Group, but which had good quality data available. These samples were assembled by the HPRC in Release 1 and later the HPRC's Production Working Group added sequencing data for these samples in an attempt to achieve uniform coverage and quality with the HRPC samples. The samples that were designated HPRC PLUS may not have low passage cell lines available.

##  Population Descriptions

When describing a sample's population in this repository we try to follow [the NHGRI's guidelines for referring to populations](https://catalog.coriell.org/1/NHGRI/About/Guidelines-for-Referring-to-Populations). The guidelines do not include the use of superpopulation so those are not included in the sample metadata index. A few samples have "N/A" for population description columns when no population descriptor was available or appropriate.

**Important note:** Release 2 contains samples from the 1000G "Mexican Ancestry in Los Angeles, California, USA" sampling (MXL). These samples were not included by the HPRC as proxies for indigenous ancestry and we advise against efforts to infer indigenous ancestry from them.

# Noteworthy And Missing Samples

For information about:
* Why certain samples with sufficient sequencing data are missing from Release 2
* Samples which are included in the release, but had either unexpected genomic features (which were either left alone or manually fixed) that may affect downstream processing

Read the `noteworthy_samples.md` document in this folder.
