## Supplementary material for "Structural polymorphism and population-variable coding capacity of HERV-K(HML-2) in human pangenomes": README.md

# Figure 7 population metadata

The retained IGSR table did not contain HG005, HG02109, HG06807 or NA21309. These four donors accounted for eight observations previously labelled Unknown in the 8q11.23 solo-LTR network. The companion `../Figure_8_population_metadata.tsv` supplements the exact-ID lookup using recorded donor metadata.

| Donor | Recorded population | Display group | Primary source |
|---|---|---|---|
| HG005 | Chinese | EAS | Coriell GM24631 ethnicity field |
| HG02109 | African Caribbean in Barbados (ACB) | AFR | HPRC sample sheet, line 224 |
| HG06807 | African Americans living in St. Louis, Missouri (ASL) | AFR | HPRC sample sheet, line 2 |
| NA21309 | Maasai in Kinyawa, Kenya (MKK) | AFR | HPRC sample sheet, line 228 |

The HPRC sheet is retained at commit `5a939042026331a823a6307fe36a3d7e0188a6e0`. Source URLs, file hashes and retrieval records accompany the downloaded metadata. Line numbers are one-based physical file lines, including the header. BioSample SAMN03283350 directly binds NIST_ID HG005 to isolate NA24631. Coriell's GM24631 record links the NA24631 DNA product and records ethnicity as Chinese. The relevant saved HTML field is `span#lblEthnicity`. HG06807's BioSample population field is empty of information, but the HPRC description is corroborated by its Coriell record.

AFR and EAS are broad display categories harmonized from the recorded population descriptions. The HPRC sample sheet does not itself provide superpopulation codes. HG005 was not assigned a specific 1000 Genomes population, and the recorded ASL label was not replaced by ASW. No population was inferred from genetic similarity or a sample-name prefix.

`eight_corrected_observations.tsv` identifies the exact donor-haplotype records, sequence ranks and FASTA hashes. Seven observations belong to the dominant sequence. The HG02109 paternal observation belongs to singleton rank 10. All 461 observations, 13 sequence haplotypes, 432 observations of the dominant sequence and sequence differences remain unchanged. AFR changes from 133 to 139 and EAS from 100 to 102. No observations remain Unknown.

The retained count table keeps its historical filename `Figure_7_solo_LTR_haplotype_counts.tsv`. It is the input for current Figure 7.
