## Supplementary material for "Structural polymorphism and population-variable coding capacity of HERV-K(HML-2) in human pangenomes": PORTABLE_REPRODUCTION.md

# Portable reproduction

The current runnable analysis and complete retained input closure are in `analysis/september2026_revision/` at the repository root. Use its `reproduce.py` command and README. The scripts in this supplementary folder preserve the original analysis history and may refer to historical workspace paths.

Final figure numbering is Figure 3 for `Table_S14/phylogeny_review`, Figure S5 for `Table_S15`, Figure S18 for `Table_S16`, Figures 5 and S13 for `Table_S17`, and Figure S6 for `Figure_S7_boundary_evidence`. Complete cassette aligned FASTA files and sequence/frequency tables in `Table_S17` are retained as final Table S11 supporting data. The separate 24-page alignment PDF is omitted from the submission. Its historical rendering remains reproducible. Earlier figure and table names in archived scripts do not identify the current manuscript labels; see `FIGURE_TABLE_MAP.md` and `SUPPLEMENTARY_TABLE_NUMBERING.tsv` at the repository root.
