## Supplementary material for "Structural polymorphism and population-variable coding capacity of HERV-K(HML-2) in human pangenomes": README.md

# Figure S7 boundary measurements and cell-type clarification

Figure S7 uses the current retained terminal-boundary evidence joined to the manuscript catalog. The historical `5′_TSD` and `3′_TSD` columns in the structural/ORF catalog are retained annotations. They are not the measurements used for this figure and are not overwritten by this package.

The source caller retained the longest adjacent 6-, 5- or 4-base comparison with at most two differences. This length rule does not establish the original TSD length. `different_pairs` and `fraction_different` in the per-locus source table describe literal retained-window comparisons. Use `length_robust_candidate_differences` for Figure S7A. The 42 length-ambiguous comparisons at 8p23.1a are not counted as TSD substitutions because all have an exact adjacent five-base pair.

The public-donor analysis includes 59,656 catalog records from 292 donors and 584 haplotypes. There are 12,095 retained paired calls, including 20 candidate differences that remain unequal across all 4–6 bp comparisons. There are 8,582 one-sided calls, 38,114 unavailable calls, and 865 records without an exact retained boundary match. Counts are physical sequence records or boundary pairs, not independent people. An array can contribute multiple copies.

`manuscript_terminal_boundary_measurements.tsv` retains every manuscript catalog record and exact source join, including the 1,453 unmatched records across the entire 61,936-record catalog. Unmatched records have blank boundary sequences and an explicit no-match state. No fuzzy or donor-only matches were used. Exactly repeated source rows were collapsed only when every field agreed. `summary.json` identifies and hashes the biological input files. These tables do not replace structural/ORF annotations.

The old all-different claims at 3q21.2, 4q32.3 and 5p12 do not survive terminal-junction validation. The old 3q12.3 population comparison is also withdrawn. The retained correction audit identifies its historical second LTR as a separate downstream element. Current source measurements resolve only one outward terminal junction at these four loci. The updated S7B therefore explains the length ambiguity at 8p23.1a instead of displaying unsupported 3q12.3 population calls. `Figure_S7_TSD_population_denominators.tsv` gives donor, haplotype and callable-record denominators for the supported current data without implying a population association.

Fiber-seq cell type is lymphoblastoid cell lines. The HPRC2 source paper describes Fiber-seq for 38 paired HPRC LCLs plus HG002 LCLs for benchmarking, producing its 39-sample consensus-peak panel. The retained local HML-2 summary contains HG002 and 38 HPRC donors, listed in `Fiberseq_cell_type_roster.tsv`. Source: https://www.biorxiv.org/content/10.64898/2026.07.21.739710v1.full, sections “A paired long-read chromatin epigenome map of the pangenome” and “Cell Line Production.”

Historical package name `Figure_S10_TSD_per_locus.tsv` refers to the previous numbering of Figure S7. Replace its contents with `Figure_S7_TSD_per_locus.tsv` and retain this mapping in the package index.
