## Supplementary material for "Structural polymorphism and population-variable coding capacity of HERV-K(HML-2) in human pangenomes": type1_drift_state_test_20260914.py

```
"""Outcome-blind Type-I/II state-spectrum sensitivities with neutral drift.

No manuscript is edited. Outputs are conditional model predictions, not p-values.
"""
import os
for key in ('OPENBLAS_NUM_THREADS','OMP_NUM_THREADS','MKL_NUM_THREADS','VECLIB_MAXIMUM_THREADS'):
    os.environ[key] = '1'
from pathlib import Path
import csv
import json
import math
import hashlib
import time
import numpy as np
from scipy.linalg import eigh_tridiagonal, expm
from scipy.optimize import minimize_scalar, brentq
from scipy.stats import binom

ROOT = Path(__file__).resolve().parent.parent
OUT = ROOT / 'recomputed_results'
SOURCE = ROOT / 'frozen_input.tsv'
OUT.mkdir(parents=True,exist_ok=True)
raw = SOURCE.read_bytes()
rows = list(csv.DictReader(raw.decode().splitlines(), delimiter='\t'))
primary = [r for r in rows if r['primary_precision_n_ge_59']=='true']
complete = [r for r in primary if r['complete_age_primary_included']=='true']
assert len(primary)==58 and len(complete)==34
assert sum(int(r['n_type1']) for r in primary)==6543
assert sum(int(r['n_type2']) for r in primary)==14505
assert all(int(r['n_type1'])*int(r['n_type2'])==0 for r in primary)
(OUT/'frozen_input.tsv').write_bytes(raw)

def grid(ne, flank, middle):
    lo=np.geomspace(1/(2*ne),.01,flank)
    x=np.unique(np.r_[0,lo,np.linspace(.01,.99,middle),1-lo[::-1],1])
    return x

def rates(x,ne,lam,recurrent):
    """Neutral diffusion plus one-way conversion, row-generator orientation.

    At zero, introductions occur at total locus rate lambda and add one copy.
    When recurrent=True, remaining Type-II copies can also convert while
    polymorphic. Interior conversion drift is lambda*(1-x)/(2Ne).
    """
    up=np.zeros(len(x));down=np.zeros(len(x))
    up[0]=lam
    hleft=x[1:-1]-x[:-2]; hright=x[2:]-x[1:-1]
    variance=x[1:-1]*(1-x[1:-1])/(2*ne)
    down[1:-1]=variance/(hleft*(hleft+hright))
    up[1:-1]=variance/(hright*(hleft+hright))
    if recurrent:
        up[1:-1]+=lam*(1-x[1:-1])/(2*ne*hright)
    return up,down

def factor(x,ne,lam,recurrent):
    up,down=rates(x,ne,lam,recurrent)
    diagonal=-(up+down)[:-1]
    off=np.sqrt(up[:-2]*down[1:-1])
    logd=np.r_[0,np.cumsum(.5*np.log(up[:-2]/down[1:-1]))]
    assert np.max(abs(logd))<300
    eig,vec=eigh_tridiagonal(diagonal,off,lapack_driver='stev')
    return eig,vec,logd,ne,up,down

def evolve(fac,t):
    eig,vec,logd,ne,up,down=fac
    # Avoid subtracting near-unit transient mass when recent-locus fixation is tiny.
    if t < ne:
        q=np.diag(-(up+down))+np.diag(up[:-1],1)+np.diag(down[1:],-1)
        p=expm(q*t)[0]
        assert p.min()>-1e-12 and abs(p.sum()-1)<2e-7
        return np.maximum(p,0)/p.sum()
    tr=((vec[0]*np.exp(eig*t))@vec.T)*np.exp(logd)
    err=max(0.,float(-tr.min()),float(tr.sum()-1))
    if err>2e-7:
        raise ArithmeticError(f'probability error {err}')
    tr=np.maximum(tr,0)
    p=np.r_[tr,max(0,1-tr.sum())]
    return p/p.sum()

def advance_without_conversion(x,ne,p,t):
    up,down=rates(x,ne,0,False)
    q=np.diag(-(up+down))+np.diag(up[:-1],1)+np.diag(down[1:],-1)
    result=p@expm(q*t)
    assert result.min()>-1e-9 and abs(result.sum()-1)<2e-7
    return np.maximum(result,0)/result.sum()

def metrics(selected,x,probabilities):
    n=np.array([int(r['n_typed']) for r in selected])
    y=np.array([int(r['n_type1'])>0 for r in selected])
    a=np.array([np.dot(p,x**ni) for p,ni in zip(probabilities,n)])
    b=np.array([np.dot(p,(1-x)**ni) for p,ni in zip(probabilities,n)])
    mixed=np.maximum(0,1-a-b)
    ll=float(np.log(a[y]).sum()+np.log(b[~y]).sum())
    joint=np.ones(1);count_i=np.ones(1);positive_count=np.ones(1)
    for ai,bi in zip(a,b):
        joint=np.convolve(joint,[bi,ai])
        count_i=np.convolve(count_i,[1-ai,ai])
        positive_count=np.convolve(positive_count,[bi,1-bi])
    k=int(y.sum())
    jt=float(joint[k:].sum()); it=float(count_i[k:].sum())
    return {'log_likelihood':ll,'expected_all_I':float(a.sum()),
            'expected_mixed':float(mixed.sum()),'expected_all_II':float(b.sum()),
            'expected_TypeI_positive':float((1-b).sum()),
            'p_zero_mixed_given_exact_observed_TypeI_positive':float(joint[k]/positive_count[k]) if positive_count[k] else None,
            'p_zero_sampled_mixed':float(np.prod(a+b)),
            'p_at_least_observed_I_and_zero_mixed':jt,
            'p_zero_mixed_given_at_least_observed_I':jt/it if it else None,
            'p_exact_observed_I_and_zero_mixed':float(joint[k]),
            'observed_all_I':k,'observed_all_II':len(selected)-k,'loci':len(selected)}

def evaluate(selected,ne,times,loglam,recurrent,flank,middle):
    x=grid(ne,flank,middle)
    fac=factor(x,ne,10**loglam,recurrent)
    cache={t:evolve(fac,t) for t in set(times)}
    return metrics(selected,x,[cache[t] for t in times])

def fit(selected,ne,times,recurrent,flank,middle,center=None):
    bounds=(-7,0) if center is None else (max(-7,center-.3),min(0,center+.3))
    result=minimize_scalar(lambda l:-evaluate(selected,ne,times,l,recurrent,flank,middle)['log_likelihood'],bounds=bounds,method='bounded',options={'xatol':.001,'maxiter':24})
    l=float(result.x)
    m=evaluate(selected,ne,times,l,recurrent,flank,middle)
    return {'lambda_per_locus_generation':10**l,'log10_lambda':l,
            'lambda_at_upper_boundary':l>-.002,'grid':f'{flank}/{middle}',**m}

def write_tsv(name,records):
    keys=list(dict.fromkeys(k for r in records for k in r))
    with (OUT/name).open('w',newline='') as f:
        w=csv.DictWriter(f,fieldnames=keys,delimiter='\t');w.writeheader();w.writerows(records)

def validation():
    result=[]
    # Independent dense matrix exponential, same finite-volume process.
    for ne,lam,recurrent in [(1280,.001,False),(10000,.03,False),(10000,.03,True)]:
        x=grid(ne,25,51);up,down=rates(x,ne,lam,recurrent)
        q=np.diag(-(up+down))+np.diag(up[:-1],1)+np.diag(down[1:],-1)
        spec=evolve(factor(x,ne,lam,recurrent),100000)
        dense=expm(q*100000)[0]
        err=float(np.max(abs(spec-dense)))
        assert err<2e-7
        result.append({'test':'spectral_vs_dense','Ne':ne,'lambda':lam,'recurrent':recurrent,'max_error':err})
    # Independent exact discrete-generation WF chain at small Ne.
    ne=40;N=2*ne;lam=.01;g=300;x=np.arange(N+1)/N
    for recurrent in [False,True]:
        transition=np.stack([binom.pmf(np.arange(N+1),N,
                min(1,xi+lam/N*(1-xi)) if recurrent or j==0 else xi)
                for j,xi in enumerate(x)])
        p=np.zeros(N+1);p[0]=1
        for _ in range(g):p=p@transition
        # Uniform-grid CTMC, no spectral grid assumptions used here.
        up,down=rates(x,ne,lam,recurrent)
        q=np.diag(-(up+down))+np.diag(up[:-1],1)+np.diag(down[1:],-1)
        ct=expm(q*g)[0]
        e1=np.array([p@(x**20),p@((1-x)**20)])
        e2=np.array([ct@(x**20),ct@((1-x)**20)])
        err=float(max(abs(e1-e2)))
        assert err<.01
        result.append({'test':'discrete_WF_vs_diffusion_small_Ne','Ne':ne,'lambda':lam,'recurrent':recurrent,'max_error':err})
    # Drift has exact neutral fixation probability equal to starting frequency.
    x=grid(10000,50,101);up,down=rates(x,10000,0,False)
    q=np.diag(-(up+down))+np.diag(up[:-1],1)+np.diag(down[1:],-1)
    h=np.linalg.solve(-q[1:-1,1:-1],q[1:-1,-1])
    err=float(max(abs(h-x[1:-1])));assert err<1e-10
    result.append({'test':'neutral_fixation_equals_frequency','max_error':err})
    write_tsv('validation.tsv',result)
    print('Validation passed',flush=True)

def main():
    start=time.time();validation();fits=[];refinements=[]
    # Full 58-locus age-free sensitivity, shared elapsed time is an assumption.
    for recurrent in [False,True]:
      for ne in [1280,10000,50000]:
        for t in [40000,80000,200000,400000,1000000]:
          base={'dataset':'all_58_shared_duration_sensitivity','Ne':ne,'recurrent_during_polymorphism':recurrent,'elapsed_generations':t,'age_scenario':'hypothetical_shared_duration_not_assigned_locus_ages'}
          m=fit(primary,ne,[t]*58,recurrent,25,51)
          fine=fit(primary,ne,[t]*58,recurrent,50,101,m['log10_lambda'])
          fits.append({**base,**fine})
          refinements.append({**base,'logL_change':abs(fine['log_likelihood']-m['log_likelihood']),'expected_mixed_change':abs(fine['expected_mixed']-m['expected_mixed']),'log_p0_change':abs(math.log(fine['p_zero_sampled_mixed'])-math.log(m['p_zero_sampled_mixed']))})
          print('full',ne,t,recurrent,'expected mixed',round(fine['expected_mixed'],3),'P0',fine['p_zero_sampled_mixed'],flush=True)
          write_tsv('fits.tsv',fits);write_tsv('refinement.tsv',refinements)
    # Actual published bounds only, never an imputed age or retired dating output.
    for ne in [1280,10000,50000]:
      for sc in ['oldest_allowed_numeric_mya','closed_lower_else_strict_upper_mya']:
       for recurrent in [False,True]:
        times=[float(r[sc])*1e6/25 for r in complete]
        base={'dataset':'34_published_age_complete_cases','Ne':ne,'recurrent_during_polymorphism':recurrent,'age_scenario':sc}
        m=fit(complete,ne,times,recurrent,25,51)
        fine=fit(complete,ne,times,recurrent,50,101,m['log10_lambda'])
        fits.append({**base,**fine})
        refinements.append({**base,'logL_change':abs(fine['log_likelihood']-m['log_likelihood']),'expected_mixed_change':abs(fine['expected_mixed']-m['expected_mixed']),'log_p0_change':abs(math.log(fine['p_zero_sampled_mixed'])-math.log(m['p_zero_sampled_mixed']))})
        print('ages',ne,sc,recurrent,'expected mixed',round(fine['expected_mixed'],3),'P0',fine['p_zero_sampled_mixed'],flush=True)
        write_tsv('fits.tsv',fits);write_tsv('refinement.tsv',refinements)
    # Exact old-pulse absorption limit. Frequency is fitted, timing not identified.
    pulses=[]
    for selected,label in [(primary,'all_58'),(complete,'34_age_complete')]:
        k=sum(int(r['n_type1'])>0 for r in selected);L=len(selected);theta=k/L
        pulses.append({'dataset':label,'theta':theta,'loci':L,'observed_all_I':k,
            'log_likelihood':k*math.log(theta)+(L-k)*math.log1p(-theta),
            'p_zero_sampled_mixed':1,'p_at_least_observed_I_and_zero_mixed':float(binom.sf(k-1,L,theta)),
            'p_exact_observed_I_and_zero_mixed':float(binom.pmf(k,L,theta)),
            'status':'analytic_limit_after_all_lineages_absorb_not_a_dated_or_rate_calibrated_history'})
    write_tsv('absorbed_pulse_limit.tsv',pulses)
    report={'input_sha256':hashlib.sha256(raw).hexdigest(),'primary_loci':58,'all_I':19,'all_II':39,'mixed':0,
            'age_complete_loci':34,'age_missing_loci':24,'selection':0,
            'scope':'Neutral post-integration conversion with drift, loss, retry, fixation and actual per-locus binomial emissions. Both rare retry-after-loss and recurrent-during-polymorphism processes tested.',
            'not_claimed':'No empirically fitted historical conversion rate, shared duration or Ne. No unconditional rejection probability. No new dating analysis.',
            'cross_locus_assumption':'Independent trajectories conditional on shared parameters. Does not model linkage, shared bottlenecks or changing donor availability.',
            'elapsed_seconds':time.time()-start,'status':'computed_pending_grid_refinement_review'}
    (OUT/'summary.json').write_text(json.dumps(report,indent=2)+'\n')
    print(json.dumps(report),flush=True)

if __name__=='__main__':main()
```
