## Supplementary material for "Structural polymorphism and population-variable coding capacity of HERV-K(HML-2) in human pangenomes": type1_literature_rate_test_20260914.py

```
"""Fixed literature-rate sensitivities, with exact neutral-diffusion moments.

No locus ages are inferred and no rate is fitted. No manuscript is edited.
The empirical per-site rate is borrowed as a successful cassette-transfer rate,
not multiplied by 292. This transport assumption is explicit in the report.
"""
import os
for key in ('OPENBLAS_NUM_THREADS', 'OMP_NUM_THREADS', 'MKL_NUM_THREADS', 'VECLIB_MAXIMUM_THREADS'):
    os.environ[key] = '1'
import csv
import json
import math
import time
import numpy as np
from scipy.linalg import expm
import type1_drift_state_test_20260914 as m

RATES = [
    ('Dumont_Eichler_2013', 'low_across_families', 6.1e-7),
    ('Dumont_Eichler_2013', 'mean_across_families', 3.2e-6),
    ('Dumont_Eichler_2013', 'high_across_families', 1.05e-5),
    ('Harpak_2017', 'lower_95CI', 0.8e-7),
    ('Harpak_2017', 'estimate', 2.5e-7),
    ('Harpak_2017', 'upper_95CI', 5e-7),
]
N = np.array([int(r['n_typed']) for r in m.primary])
K = sum(int(r['n_type1']) > 0 for r in m.primary)

def exact_moments(ne, mu, generations, sample_sizes=None):
    """L f = mu(1-x)f' + x(1-x)/(4Ne)f'', x(0)=0.

    a_n=E[x^n]: a'_n=(n*mu+n*(n-1)/(4Ne))*(a_(n-1)-a_n).
    b_n=E[(1-x)^n]: b'_n=n*(n-1)/(4Ne)*(b_(n-1)-b_n)-n*mu*b_n.
    Both are triangular positive linear systems. No tiny tail is obtained by
    subtracting near-unit transient probability, and there is no grid error.
    """
    sample_sizes = N if sample_sizes is None else np.asarray(sample_sizes, dtype=int)
    n = np.arange(sample_sizes.max()+1, dtype=float)
    d = n*(n-1)/(4*ne)
    c = d+n*mu
    qa = np.diag(-c)+np.diag(c[1:], -1)
    qb = np.diag(-c)+np.diag(d[1:], -1)
    a = expm(qa*generations)[:, 0]
    b = expm(qb*generations) @ np.ones(len(n))
    assert abs(a[1]-(-math.expm1(-mu*generations))) < 1e-10
    assert abs(b[1]-math.exp(-mu*generations)) < 1e-10
    assert a.min() >= 0 and b.min() >= 0
    assert np.max(a[1:]+b[1:]) <= 1+1e-10
    return a[sample_sizes], b[sample_sizes]

def summarize(a, b):
    mixed = 1-a-b
    assert mixed.min() > -1e-9
    joint = np.ones(1)
    positive_count = np.ones(1)
    for ai, bi in zip(a, b):
        joint = np.convolve(joint, [bi, ai])
        positive_count = np.convolve(positive_count, [bi, 1-bi])
    return {
        'expected_all_I': float(a.sum()),
        'expected_mixed': float(mixed.sum()),
        'expected_all_II': float(b.sum()),
        'expected_TypeI_positive': float((1-b).sum()),
        'p_zero_mixed_given_19_TypeI_positive': float(joint[K]/positive_count[K]),
        'p_exact_19_TypeI_positive': float(positive_count[K]),
        'p_exact_19_allI_39_allII_zero_mixed': float(joint[K]),
        'p_zero_mixed': float(np.prod(a+b)),
    }

def main():
    start = time.time()
    results = []
    checks = []
    # The original table's full range is primary. The newer paper is a
    # separately reported estimate, not a reason to discard the earlier range.
    for ne in [10000, 5000, 20000]:
      for source, label, mu in RATES:
        for years in [2000000, 5000000, 8000000]:
          t = years/25
          a, b = exact_moments(ne, mu, t)
          result = {'source': source, 'rate_label': label,
                    'borrowed_transfer_rate_per_chromosome_generation': mu,
                    'Ne': ne, 'elapsed_years_assumed': years,
                    'generation_years_assumed': 25, **summarize(a, b)}
          results.append(result)
          print(json.dumps(result), flush=True)
          if ne == 10000 and years == 2000000:
            x = m.grid(ne, 200, 401)
            p = m.evolve(m.factor(x, ne, 2*ne*mu, True), t)
            fa = np.array([p@(x**ni) for ni in N])
            fb = np.array([p@((1-x)**ni) for ni in N])
            approx = summarize(fa, fb)
            checks.append({'source': source, 'rate_label': label,
                'max_allI_absolute_error_finite_volume': float(max(abs(fa-a))),
                'max_allII_absolute_error_finite_volume': float(max(abs(fb-b))),
                'log10_conditional_probability_difference': abs(math.log10(approx['p_zero_mixed_given_19_TypeI_positive'])-math.log10(result['p_zero_mixed_given_19_TypeI_positive']))})
    m.write_tsv('literature_rate_ongoing_exact.tsv', results)
    m.write_tsv('literature_rate_independent_numerical_check.tsv', checks)
    print('COMPLETE', len(results), 'scenarios', time.time()-start, 'seconds', flush=True)

if __name__ == '__main__':
    main()
```
