## Supplementary material for "Structural polymorphism and population-variable coding capacity of HERV-K(HML-2) in human pangenomes": type1_observed_loci_uniformity_20260914.py

```
"""Test zero mixed samples at the actual Type-I-positive locus identities.

Condition on Type I being detected, NOT on fixation. Use individual older age
bounds and 2 Ma for recent entries. Unaged loci are granted probability one
of uniformity, yielding an upper bound within the stated neutral model.
"""
import type1_literature_rate_test_20260914 as lit
import type1_drift_state_test_20260914 as m
import numpy as np
import math
import json
import time

POSITIVE = [r for r in m.primary if int(r['n_type1']) > 0]
DATED = [r for r in POSITIVE if r['complete_age_primary_included']=='true']
RECENT = [r for r in DATED if r['age_relation']=='strict_upper_resolution_limit']
assert len(POSITIVE)==19 and len(DATED)==16 and len(RECENT)==6

def main():
    start=time.time(); summary=[]; per_locus=[]; checks=[]
    for ne in [10000,5000,20000]:
      for source,label,mu in lit.RATES:
        # Exact moments are shared across loci with the same time and n.
        required={}
        for r in DATED:
          age=float(r['oldest_allowed_numeric_mya'])
          for age_case in [age, age if r in RECENT else 2*age]:
            required.setdefault(age_case,set()).add(int(r['n_typed']))
        probabilities={}
        for age, ns in sorted(required.items()):
          ns=sorted(ns)
          a,b=lit.exact_moments(ne,mu,age*1e6/25,ns)
          for ni,ai,bi in zip(ns,a,b): probabilities[(age,ni)]=(float(ai),float(bi))
        for scenario in ['individual_older_upper_bounds','double_older_upper_bounds','older_loci_unlimited_time']:
          q=[]; which=[]
          for r in DATED:
            age=float(r['oldest_allowed_numeric_mya'])
            if r not in RECENT:
              if scenario=='older_loci_unlimited_time': continue
              if scenario=='double_older_upper_bounds': age*=2
            n=int(r['n_typed']); a,b=probabilities[(age,n)]
            p=a/(1-b)
            assert 0<=p<=1
            q.append(p);which.append(r['locus_id'])
            per_locus.append({'source':source,'rate_label':label,'rate_per_generation':mu,
              'Ne':ne,'scenario':scenario,'locus':r['locus_id'],'n_typed':n,
              'age_bound_Ma':age,'p_all_I':a,'p_all_II':b,
              'p_sample_mixed':1-a-b,'p_all_I_given_TypeI_detected':p})
          result={'source':source,'rate_label':label,'rate_per_generation':mu,'Ne':ne,
            'scenario':scenario,'observed_TypeI_positive_loci':19,
            'modeled_loci':len(q),'loci_granted_certain_uniformity':19-len(q),
            'upper_bound_p_zero_mixed_given_observed_positive_identities':math.prod(q),
            'expected_mixed_among_modeled_positive_loci':sum(1-v for v in q),
            'included_loci':','.join(which)}
          summary.append(result)
          print(json.dumps({k:v for k,v in result.items() if k!='included_loci'}),flush=True)
        if ne==10000:
          x=m.grid(ne,200,401);fac=m.factor(x,ne,2*ne*mu,True)
          exact_values=[];approx_values=[]
          for r in DATED:
            age=float(r['oldest_allowed_numeric_mya']); n=int(r['n_typed'])
            p=m.evolve(fac,age*1e6/25)
            a=p@(x**n); b=p@((1-x)**n)
            aa,bb=probabilities[(age,n)]
            exact_values.append(aa/(1-bb));approx_values.append(a/(1-b))
          err=abs(math.log10(math.prod(exact_values))-math.log10(math.prod(approx_values)))
          assert err<0.03
          checks.append({'source':source,'rate_label':label,
            'exact_vs_finite_volume_log10_probability_difference':err})
        m.write_tsv('observed_locus_uniformity_summary.tsv',summary)
        m.write_tsv('observed_locus_uniformity_per_locus.tsv',per_locus)
        m.write_tsv('observed_locus_uniformity_numerical_checks.tsv',checks)
    status={'complete':True,'elapsed_seconds':time.time()-start,
      'condition':'The exact observed set of Type-I-positive loci, not just its size',
      'recent_loci':[r['locus_id'] for r in RECENT],
      'unaged_loci_granted_probability_one':[r['locus_id'] for r in POSITIVE if r not in DATED],
      'sample_dependence':'Shared within-locus drift trajectory, binomial sampling conditional on its frequency',
      'cross_locus_dependence':'Independent conditional on Ne, rates and ages',
      'scope':'Neutral recurrent one-way conversion in a fixed occupied population. Not a reconstructed insertion-frequency history or a complete RT-versus-conversion likelihood comparison.'}
    (m.OUT/'observed_locus_uniformity_status.json').write_text(json.dumps(status,indent=2)+'\n')
    print(json.dumps(status),flush=True)

if __name__=='__main__':main()
```
