## Supplementary material for "Structural polymorphism and population-variable coding capacity of HERV-K(HML-2) in human pangenomes": README.txt

Recurrent conversion and within-locus Type-I/II polymorphism

The summary contains all 54 scenarios. Joint probabilities condition on the exact observed Type-I-positive locus identities. Unaged loci are assigned certain uniformity. See Supplemental Methods for assumptions.

To reproduce, use Python with NumPy and SciPy and run analysis_code/type1_observed_loci_uniformity_20260914.py. Results are written to recomputed_results. The archived input is frozen_input.tsv. Only filesystem paths were adapted from the original calculation.
