## Supplementary material for "Structural polymorphism and population-variable coding capacity of HERV-K(HML-2) in human pangenomes": README.txt

Figure 3C restores the relationship between 4q35.2 and the acrocentric Type-II group. The regional comparison uses the current selected KCON-aligned sequence clusters. It counts mismatches only at jointly called A/C/G/T positions. Shared host-flank alignments are retained from the original reference-boundary analysis. This is distinct from complete exact gene-sequence identity in Figure 3B. No duplication date is inferred.
