## Supplementary material for "Structural polymorphism and population-variable coding capacity of HERV-K(HML-2) in human pangenomes": analyze.py

```
import os
os.environ['OPENBLAS_NUM_THREADS']='1'
os.environ['OMP_NUM_THREADS']='1'
import csv,json,time,sys,copy
from pathlib import Path
from collections import Counter,defaultdict
from hashlib import sha256
import numpy as np
from Bio import Phylo,SeqIO
from Bio.Phylo.BaseTree import Tree,Clade
from Bio.Phylo.TreeConstruction import DistanceTreeConstructor,DistanceMatrix
OUT=Path(__file__).resolve().parent
ROOT=OUT.parents[2]
PHY=ROOT/'project/results/acroc_resolved_20260914/phylogeny'
NBOOT=1000
SEED=20260921

def read(p): return list(csv.DictReader(p.open(),delimiter='\t'))
def write(name,rows):
 with (OUT/name).open('w') as f:
  w=csv.DictWriter(f,fieldnames=list(rows[0]),delimiter='\t',lineterminator='\n');w.writeheader();w.writerows(rows)
def nj(d,names):
 # Same NJ Q criterion and lower-triangle pair ordering as Biopython.
 d=d.copy();nodes=[Clade(name=n) for n in names]
 while len(nodes)>3:
  n=len(nodes);s=d.sum(axis=1)/(n-2);q=d-s[:,None]-s[None,:]
  ii,jj=np.tril_indices(n,-1);a=np.argmin(q[ii,jj]);i,j=int(ii[a]),int(jj[a])
  nodes[i].branch_length=max(0.,float((d[i,j]+s[i]-s[j])/2))
  nodes[j].branch_length=max(0.,float(d[i,j]-(d[i,j]+s[i]-s[j])/2))
  c=Clade(clades=[nodes[i],nodes[j]])
  nd=(d[i]+d[j]-d[i,j])/2
  d[j,:]=nd;d[:,j]=nd;d[j,j]=0
  d=np.delete(np.delete(d,i,0),i,1);nodes[j]=c;del nodes[i]
 a,b,c=nodes;ab,ac,bc=d[0,1],d[0,2],d[1,2]
 a.branch_length=max(0.,float((ab+ac-bc)/2));b.branch_length=max(0.,float((ab+bc-ac)/2));c.branch_length=max(0.,float((ac+bc-ab)/2))
 return Tree(root=Clade(branch_length=0,clades=nodes),rooted=False)
def split_key(s,universe):
 t=universe-s
 return tuple(sorted(s if (len(s),tuple(sorted(s)))<(len(t),tuple(sorted(t))) else t))
def splits(t):
 u={x.name for x in t.get_terminals()};out=set()
 for c in t.get_nonterminals():
  s={x.name for x in c.get_terminals()}
  if 1<len(s)<len(u)-1:out.add(split_key(s,u))
 return out

def boot():
 rows=read(PHY/'selected_sequence_clusters.tsv')
 regions={r:[x for x in rows if x['gene']==r] for r in ['LTR','pol']}
 trees={r:Phylo.read(PHY/('hml2_pan_ltr_expanded_tree.nwk' if r=='LTR' else 'hml2_pan_orf_pol_tree.nwk'),'newick') for r in regions}
 common=set.intersection(*[{t.name.split('__')[0] for t in v.get_terminals()} for v in trees.values()])
 supportrows=[];nearestrows=[];metadata={};replicate_rows=[]
 for region,rr in regions.items():
  names=[r['tip'] for r in rr];a=np.array([list(r['sequence']) for r in rr]);v=np.isin(a,list('ACGT'));n,L=a.shape
  ii,jj=np.tril_indices(n,-1)
  valid=(v[ii]&v[jj]).astype(np.float32)
  mism=((a[ii]!=a[jj]) & (valid>0)).astype(np.float32)
  assert np.all(valid.sum(1)>0)
  base=np.zeros((n,n));base[ii,jj]=base[jj,ii]=mism.sum(1)/valid.sum(1)
  t0=time.time();ref=nj(base,names)
  # Comparison with Bio NJ using exactly the same numerical distance matrix.
  bio=DistanceTreeConstructor().nj(DistanceMatrix(names,[list(base[i,:i+1]) for i in range(n)]))
  for c in bio.find_clades():
   if c.branch_length is not None:c.branch_length=max(0,c.branch_length)
  sp=splits(ref);bs=splits(bio)
  # Q ties can alter resolutions with zero-length edges; quantify, never conceal.
  diff=max(abs(ref.distance(names[i],names[j])-bio.distance(names[i],names[j])) for i,j in zip(ii,jj))
  original=trees[region]
  original_diff=max(abs(ref.distance(names[i],names[j])-original.distance(names[i],names[j])) for i,j in zip(ii,jj))
  print(region,'tips',n,'fast NJ',time.time()-t0,'Bio maxdiff',diff,'retained maxdiff',original_diff,flush=True)
  assert diff<1e-6 and original_diff<1e-6
  counts=Counter();neigh=defaultdict(Counter);rng=np.random.default_rng(SEED+(region=='pol'))
  focal=[next(n for n in names if n.startswith(loc+'__hap1__')) for loc in ['19p12c','10q24.2','4q35.2_hg38']]
  for rep in range(NBOOT):
   w=rng.multinomial(L,np.repeat(1/L,L)).astype(np.float32)
   den=valid@w;assert np.all(den>0)
   d=np.zeros((n,n));d[ii,jj]=d[jj,ii]=(mism@w)/den
   t=nj(d,names);counts.update(splits(t))
   for f in focal:
    ds=[(t.distance(f,x),x) for x in names if x.split('__')[0] in common and x.split('__')[0]!=f.split('__')[0]]
    best=min(x[0] for x in ds);nn=sorted({x.split('__')[0] for z,x in ds if abs(z-best)<1e-10})
    for x in nn:neigh[f][x]+=1/len(nn)
    replicate_rows.append(dict(region=region,replicate=rep+1,focal_tip=f,nearest_loci=';'.join(nn),nearest_distance=best))
   if (rep+1)%100==0: print(region,rep+1,round(time.time()-t0,1),'sec',flush=True)
  universe=set(names)
  for c in original.get_nonterminals():
   s={x.name for x in c.get_terminals()}
   if 1<len(s)<len(names)-1:
    key=split_key(s,universe);c.confidence=100*counts[key]/NBOOT
    supportrows.append(dict(region=region,split=';'.join(key),tip_count=len(key),bootstrap_replicates=NBOOT,bootstrap_count=counts[key],bootstrap_percent=c.confidence))
  for f in focal:
   ds=[(original.distance(f,x),x) for x in names if x.split('__')[0] in common and x.split('__')[0]!=f.split('__')[0]]
   best=min(z for z,x in ds);basenn={x.split('__')[0] for z,x in ds if abs(z-best)<1e-10}
   for loc,count in neigh[f].most_common():
    nearestrows.append(dict(region=region,focal_tip=f,nearest_locus=loc,retained_tree_neighbor=int(loc in basenn),bootstrap_weighted_count=count,bootstrap_percent=100*count/NBOOT,comparison_loci=len(common)))
  Phylo.write(original,OUT/f'{region}_bootstrap.nwk','newick',format_branch_length='%1.15g')
  metadata[region]=dict(tips=n,loci=len({n.split('__')[0] for n in names}),aligned_positions=L,bootstrap=NBOOT,seed=SEED+(region=='pol'),NJ_implementation_max_pair_distance_difference=diff,retained_tree_max_pair_distance_difference=original_diff,all_baseline_splits_equal=splits(ref)==splits(original),elapsed_seconds=time.time()-t0)
 write('bootstrap_splits.tsv',supportrows);write('bootstrap_nearest_loci.tsv',nearestrows);write('bootstrap_nearest_loci_replicates.tsv',replicate_rows)
 (OUT/'bootstrap_metadata.json').write_text(json.dumps(metadata,indent=2)+'\n')

def apd():
 edges=read(ROOT/'outputs/hml2-pangenome-2026-09-13/Supplementary_Data/Figure_3B_exact_nucleotide_edges.tsv')
 loci={r[k] for r in edges for k in ['locus_1','locus_2']}
 observations=[r for r in read(PHY/'region_observations.tsv') if r['locus'] in loci and r['region']!='LTR']
 admissions={r['ID_Full']:r for r in read(PHY/'alignment_admission.tsv') if r['locus'] in loci}
 inputrows=read(PHY/'alignment_inputs.tsv');paths={Path(r['path']).name:Path(r['path']) for r in inputrows}
 cache={};inputs=[]
 for path in {paths[r['alignment']] for r in admissions.values() if r['alignment'] in paths}:
  digest=sha256(path.read_bytes()).hexdigest();assert digest==next(r['sha256'] for r in inputrows if r['path']==str(path))
  cache.update({(path.name,r.id):str(r.seq).upper() for r in SeqIO.parse(path,'fasta')});inputs.append(dict(path=str(path),sha256=digest))
 projected_path=Path(inputrows[-1]['path'])
 if any((r['alignment'],r['record']) not in cache for r in admissions.values()):
  assert sha256(projected_path.read_bytes()).hexdigest()==inputrows[-1]['sha256']
  for r in read(projected_path):cache[(Path(r['alignment_path']).name,r['alignment_record_id'])]=r['typeII_KCON_sequence']
  inputs.append(inputrows[-1])
 spans={'gag':(1111,3112),'pro':(2913,3918),'pol':(3878,6749),'env':(6450,8550)}
 per=defaultdict(Counter);source_rows=[]
 for r in observations:
  adm=admissions[r['ID_Full']];seq=cache[(adm['alignment'],adm['record'])]
  assert sha256(seq.encode()).hexdigest()==adm['aligned_sequence_sha256']
  seq=seq[slice(*spans[r['region']])]
  assert sha256(seq.encode()).hexdigest()==r['sequence_sha256']
  per[r['locus'],r['region']][seq]+=1;source_rows.append(r)
 # Xq28 has source-bound Env sequences but no retained KCON panel.
 xqaln={r.id:str(r.seq).upper() for r in SeqIO.parse(OUT/'Xq28_env_aligned.fasta','fasta')}
 for r in read(OUT/'Xq28_env_source_observations.tsv'):
  seq=xqaln[r['sha256']]
  assert sha256(seq.replace('-', '').encode()).hexdigest()==r['sha256']
  per[r['locus'],'env'][seq]+=1
 results=[];summary=[]
 for edge in edges:
  l1,l2=edge['locus_1'],edge['locus_2'];sum_dist=0;npairs=0;totalmis=0;totalvalid=0
  for gene in spans:
   x,y=per[l1,gene],per[l2,gene]
   if not x or not y:continue
   weighted=0;count=0;mm=0;vv=0;minvalid=10000;minp=1;maxp=0
   for s,c1 in x.items():
    a=np.array(list(s));v=np.isin(a,list('ACGT'))
    b=np.array([list(z) for z in y]);w=np.array(list(y.values()))*c1
    valid=np.isin(b,list('ACGT')) & v
    den=valid.sum(1);num=((a!=b)&valid).sum(1)
    assert np.all(den>0)
    p=num/den;weighted+=float(p@w);count+=int(w.sum());mm+=int(num@w);vv+=int(den@w)
    minvalid=min(minvalid,int(den.min()));minp=min(minp,float(p.min()));maxp=max(maxp,float(p.max()))
   sum_dist+=weighted;npairs+=count;totalmis+=mm;totalvalid+=vv
   results.append(dict(locus_1=l1,locus_2=l2,region=gene,left_source_copies=sum(x.values()),right_source_copies=sum(y.values()),left_distinct_sequences=len(x),right_distinct_sequences=len(y),source_copy_pairs=count,mean_pairwise_difference=weighted/count,percent_difference=100*weighted/count,mismatch_pairs=mm,callable_base_pairs=vv,minimum_callable_positions=minvalid,minimum_pair_difference=minp,maximum_pair_difference=maxp))
  summary.append(dict(locus_1=l1,locus_2=l2,mean_pairwise_difference=sum_dist/npairs,percent_difference=100*sum_dist/npairs,source_copy_gene_pairs=npairs,mismatch_pairs=totalmis,callable_base_pairs=totalvalid,distinct_identical_sequences=int(edge['distinct_gene_sequences'])))
 write('pairwise_differences_by_region.tsv',results);write('pairwise_differences_network.tsv',summary);write('pairwise_source_observations.tsv',source_rows);write('pairwise_input_files.tsv',inputs)
 print('APD',json.dumps(summary,indent=2),flush=True)
if __name__=='__main__':
 if '--apd' in sys.argv:apd()
 else:boot()
```
