## Supplementary material for "Structural polymorphism and population-variable coding capacity of HERV-K(HML-2) in human pangenomes": METHODS.md

# Figure 3 bootstrap and pairwise nucleotide differences

This package addresses John Coffin's comments 102, 103 and 105. It adds no ORF-break dating, insertion ages or duplication-order claims.

## Tree input and inference

The retained source-bound KCON projections and their region-level sequence clusters are from `project/results/acroc_resolved_20260914/phylogeny`. The LTR input contains 162 exact sequence clusters from 82 loci, aligned over 968 KCON positions. The Pol input contains 138 clusters from 70 loci, aligned over 2,871 positions. Each locus contributes up to its two most frequent exact aligned clusters. Cluster counts record source-copy observations and do not weight the distance matrix or bootstrap sampling. c1/c2 are ranked separately within each region and are not phased across regions.

The retained LTR extraction uses KCON positions [8504,9472), falling back to [0,968) if needed. It requires at least 600 A/C/G/T bases. Pol is [3878,6749) and requires at least 50% canonical bases. Pairwise distance is the number of A/C/G/T mismatches divided by positions canonical in both sequences. Gaps and ambiguous bases are excluded separately for each sequence pair. No multiple-substitution correction is applied. The supplied pairwise-distance table contains the actual mismatch counts and denominators.

The bootstrap uses 1,000 replicates per region, resampling the original number of alignment columns with replacement. It keeps the complete retained cluster set fixed, recalculates the same pairwise distance and rebuilds the neighbor-joining tree. Negative inferred branch lengths are set to zero, matching the existing producer. A vectorized implementation uses the same Q criterion and lower-triangle pair ordering as Biopython. The baseline implementation yields the same unrooted splits as the retained trees. The maximum difference in pairwise tree distance from the retained Newicks is below 4e-9, attributable to numeric precision. Against Biopython using the same distance matrix, the maximum difference is below 3e-17. Random seeds are 20260921 for LTR and 20260922 for Pol.

Support is the frequency of each unrooted bipartition. The figure uses the retained tree topology and branch lengths, then positions the display at its midpoint. Support is reassigned by the unrooted split after repositioning. The display is pruned to the modal cluster at every locus and the extra nearest-neighbor clusters used in the focal comparisons. Values of at least 70% are labeled. Their bipartitions refer to the full input trees. Display positioning does not infer a biological root or evolutionary time. The complete trees, alignments and support table accompany the figure.

## Focal nearest neighbors

For each full retained tree and each bootstrap replicate, the modal cluster of 19p12c, 10q24.2 or 4q35.2 is compared against every other-locus cluster in the same fixed pool of 64 loci represented in both LTR and Pol. Distances are sums of nonnegative tree branch lengths. Distances within 1e-10 of the minimum count as tied. If several tied clusters belong to the same locus, that locus is counted once. A replicate containing k tied loci contributes 1/k to each locus. Every replicate's tied-locus set and shortest distance are retained.

The nearest-neighbor recovery percentages are not internal-branch bootstrap support. They summarize stability of the specified nearest-locus estimator. They are not a formal test of recombination. The observed 19p12c and 10q24.2 switches are weakly recovered and cannot support a strong exchange claim on their own.

The unrestricted LTR nearest neighbor of 10q24.2 is 19p12d c1 at 0.02683040380548737. Because 19p12d has no Pol input, the paired comparison excludes it. The shared-pool LTR neighbor is 12q14.1 c2 at 0.027817259455631602. The next candidates are 12q14.1 c1 at 0.027836991912202354 and 3q13.2 c1 at 0.027952583084347848. These are not ties and are not changed by floating-point differences. The supplied focal-distance table retains both unrestricted and shared-pool results.

## Panel B mean differences

The existing 13 exact-sharing edges are preserved. Their counts continue to appear in `pairwise_differences_network.tsv`. New labels are average pairwise nucleotide differences, expressed as percentages. They include nonidentical sequences at those connected loci.

For 12 edges, every source-copy observation already admitted to the retained KCON region panel was recovered through its recorded alignment and sequence digest. The gene spans are gag [1111,3112), pro [2913,3918), pol [3878,6749) and env [6450,8550). A region must already meet the producer's 50% canonical-coverage threshold. Each left-locus copy is compared with each right-locus copy for each region available in both loci. The sequence-pair difference is mismatches divided by comparable A/C/G/T sites. The network label is the arithmetic mean across all available source-copy/gene-region comparisons. Identical aligned sequence clusters are collapsed for calculation and weighted by their actual copy counts, producing the same mean as enumerating all observed copy pairs.

The four genes are not treated as equally weighted replicates, and this is not a concatenated whole-provirus divergence. Gene regions with more admitted source pairs contribute more comparisons. Overlapping KCON gene spans remain part of their respective region measurements. Region-specific means, sample counts, comparable-base counts and minimum callable coverage are supplied so comparisons can be made on matching regions. Pair counts are descriptive combinatorial denominators, not independent population sample sizes.

Xq28 has no source-bound input in the retained KCON panel. Its existing exact-sharing dataset contains Env observations from 214 Xq28a copies and 157 Xq28b copies. Their 371 raw source FASTA byte digests were reverified, and every retained native sequence exactly matches its current source FASTA. Each Env sequence was recovered from its native sequence by its retained length and SHA-256 digest. The 17 distinct Env sequences were aligned with MAFFT --auto. The result contains 2,110 columns, with no gaps in any sequence. This comparison includes 33,598 copy pairs and 70,891,780 comparable base-pairs. There are 9,499 mismatched base-pairs, giving a mean difference of 0.0001339929678730031 or 0.013399296787300312%. All copies have public HG/NA identifiers and each sample/haplotype/source identity occurs once. Native sequence, alignment and source tables are supplied.

## Scope of verification

No new source admission, physical-locus reassignment or raw-sequence repair was performed. The KCON analyses retain the source-panel coverage limits and fixed reference projection, which omits insertions outside KCON positions. The reported values measure substitutions at comparable aligned bases and do not count indel length as nucleotide mismatches. These results do not imply that a pair sharing one exact gene sequence has uniformly identical copies.

The figure is 6.5 by 7.25 inches. Full-resolution PNG, PDF and editable SVG are provided. Highlighted 19p12c, 10q24.2 and 4q35.2 labels and the existing chromosomal connections are preserved. The old support-free inference and distinct-sequence edge labels are replaced. All execution and outputs for this extension are confined to this directory.
