## Supplementary material for "Structural polymorphism and population-variable coding capacity of HERV-K(HML-2) in human pangenomes": package.py

```
from pathlib import Path
import csv,json,shutil
from hashlib import sha256
import numpy as np
from Bio import Phylo
O=Path(__file__).resolve().parent;ROOT=O.parents[2];P=ROOT/'project/results/acroc_resolved_20260914/phylogeny'
def read(p):return list(csv.DictReader(p.open(),delimiter='\t'))
def write(name,rows):
 with (O/name).open('w') as f:
  w=csv.DictWriter(f,fieldnames=list(rows[0]),delimiter='\t',lineterminator='\n');w.writeheader();w.writerows(rows)
selected=[r for r in read(P/'selected_sequence_clusters.tsv') if r['gene'] in ['LTR','pol']]
write('tree_selected_sequence_clusters.tsv',selected)
inputs=[P/'selected_sequence_clusters.tsv',P/'alignment_admission.tsv',P/'region_observations.tsv',P/'alignment_inputs.tsv',P/'hml2_pan_ltr_expanded_tree.nwk',P/'hml2_pan_orf_pol_tree.nwk']
write('retained_input_files.tsv',[dict(path=str(p),sha256=sha256(p.read_bytes()).hexdigest()) for p in inputs])
point=[];all_d=[];common=set.intersection(*[{r['locus'] for r in selected if r['gene']==g} for g in ['LTR','pol']])
for gene in ['LTR','pol']:
 rr=[r for r in selected if r['gene']==gene];tree=Phylo.read(O/f'{gene}_bootstrap.nwk','newick')
 with (O/f'{gene}_aligned_sequences.fasta').open('w') as f:
  for r in rr:f.write('>'+r['tip']+'\n'+r['sequence']+'\n')
 for i,r in enumerate(rr):
  for s in rr[:i]:
   pairs=[(a,b) for a,b in zip(r['sequence'],s['sequence']) if a in 'ACGT' and b in 'ACGT'];m=sum(a!=b for a,b in pairs)
   all_d.append(dict(region=gene,tip_1=r['tip'],tip_2=s['tip'],mismatches=m,comparable_positions=len(pairs),nucleotide_difference=m/len(pairs),tree_distance=tree.distance(r['tip'],s['tip'])))
 for loc in ['19p12c','10q24.2','4q35.2_hg38']:
  f=next(r['tip'] for r in rr if r['locus']==loc and '__hap1__' in r['tip'])
  for pool in ['all_retained_loci','shared_64_loci']:
   vals=sorted((tree.distance(f,r['tip']),r['tip']) for r in rr if r['locus']!=loc and (pool=='all_retained_loci' or r['locus'] in common));best=vals[0][0]
   for d,t in vals:point.append(dict(region=gene,focal_tip=f,candidate_pool=pool,other_tip=t,tree_distance=d,nearest=int(abs(d-best)<1e-10)))
write('tree_pairwise_distances.tsv',all_d);write('focal_tree_distance_comparisons.tsv',point)
# Explicit conditional denominators for each network edge.
regions=read(O/'pairwise_differences_by_region.tsv')
write('network_sequence_denominators.tsv',[dict(locus_1=r['locus_1'],locus_2=r['locus_2'],region=r['region'],left_source_copies=r['left_source_copies'],right_source_copies=r['right_source_copies'],source_copy_pairs=r['source_copy_pairs'],minimum_callable_positions=r['minimum_callable_positions'],alignment='native Env MAFFT' if r['locus_1']=='Xq28a' else 'retained KCON projection') for r in regions])
files=['Figure_3.svg','Figure_3.pdf','Figure_3.png','LTR_bootstrap.nwk','pol_bootstrap.nwk','LTR_aligned_sequences.fasta','pol_aligned_sequences.fasta','tree_selected_sequence_clusters.tsv','tree_pairwise_distances.tsv','focal_tree_distance_comparisons.tsv','bootstrap_splits.tsv','bootstrap_nearest_loci.tsv','bootstrap_nearest_loci_replicates.tsv','bootstrap_metadata.json','pairwise_differences_by_region.tsv','pairwise_differences_network.tsv','network_sequence_denominators.tsv','pairwise_source_observations.tsv','pairwise_input_files.tsv','retained_input_files.tsv','Xq28_env_exact_sequences.fasta','Xq28_env_aligned.fasta','Xq28_env_source_observations.tsv','Xq28_native_source_files.tsv','analyze.py','xq28.py','render_figure3.py','prepare_render.py','package.py','manuscript_insertions.md','METHODS.md']
(O/'package_files.json').write_text(json.dumps(files,indent=2)+'\n')
print('Packaged',len(files),'files;',len(all_d),'tree distances;',len(point),'focal comparisons')
```
