## Supplementary material for "Structural polymorphism and population-variable coding capacity of HERV-K(HML-2) in human pangenomes": prepare_render.py

```
from pathlib import Path
OUT=Path(__file__).resolve().parent
ROOT=OUT.parents[2]
s=(ROOT/'project/manuscript/restore_figure3_20260914.py').read_text()
s=s.replace("import build_narrative_main_figures as owner", "sys.path.insert(0,"+repr(str(ROOT/'project/manuscript'))+")\nimport build_narrative_main_figures as owner")
s=s.replace("PROJECT = Path(__file__).resolve().parents[1]",'PROJECT = Path('+repr(str(ROOT/'project'))+')')
s=s.replace("OUT = PROJECT/'manuscript/figures/comment_corrections_20260914'",'OUT = Path(__file__).resolve().parent')
s=s.replace("edge_map={tuple(sorted((r['locus_1'],r['locus_2']))):int(r['distinct_gene_sequences']) for r in edges}","edge_map={tuple(sorted((r['locus_1'],r['locus_2']))):float(r['percent_difference']) for r in csv.DictReader((OUT/'pairwise_differences_network.tsv').open(),delimiter='\\t')}")
s=s.replace("full=Phylo.read(PHY/filename,'newick')",'''full=Phylo.read(OUT/('LTR_bootstrap.nwk' if region=='LTR' else 'pol_bootstrap.nwk'),'newick')
    from analyze import split_key
    support_rows=list(csv.DictReader((OUT/'bootstrap_splits.tsv').open(),delimiter='\\t'))
    support={tuple(r['split'].split(';')):float(r['bootstrap_percent']) for r in support_rows if r['region']==('LTR' if region=='LTR' else 'pol')}
    universe={x.name for x in full.get_terminals()}
    full.root_at_midpoint()
    full.rooted=False
    for c in full.get_nonterminals():
        c.confidence=support.get(split_key({x.name for x in c.get_terminals()},universe))''')
s=s.replace("tips=tree.get_terminals()", "tree.ladderize()\n    tips=tree.get_terminals()",1)
s=s.replace("    # Alternate label columns",'''    for c in tree.get_nonterminals():
        if c is not tree.root and c.confidence is not None and c.confidence >=70:
            ax.text(depths[c],ypos[c]+.52,str(round(c.confidence)),fontsize=4.4,ha='right',va='bottom',color='#333333',bbox={'fc':'white','ec':'none','pad':.03},zorder=6)
    # Alternate label columns''')
s=s.replace("ax.text(target,ypos[tip],display,fontsize=6.4,color=colors[subfamilies[locus]],va='center')", "ax.text(target,ypos[tip],display,fontsize=6.4,color=colors[subfamilies[locus]],va='center',fontweight='bold' if locus in ['19p12c','10q24.2','4q35.2_hg38'] else 'normal')")
s=s.replace("ax.set_xlabel('Substitutions per site',fontsize=7,labelpad=1)","ax.set_xlabel('Nucleotide differences per aligned base',fontsize=6.5,labelpad=1)")
s=s.replace("'Shared nucleotide sequences'","'Mean nucleotide difference (%)'")
s=s.replace("str(edge_map[key]),fontsize=7", "f'{edge_map[key]:.2f}' if edge_map[key]>=.1 else f'{edge_map[key]:.3f}',fontsize=6.1")
s=s.replace("'Figure_3_restored", "'Figure_3")
s=s.replace("ax.text(x,.94 if label", "ax.text(x,.94 if label")
# Keep chromosome focal label bold as in the reviewed Figure 3.
s=s.replace("ha='center',fontsize=6.3)","ha='center',fontsize=6.3,fontweight='bold' if label=='4q35.2' else 'normal')")
# No mutations or unrelated panels.
s=s[:s.index("svg=(INPUT/'Figure_S5")]
s=s.replace("coords=[(cx-.125,cy+.09),(cx+.125,cy+.09),(cx-.125,cy-.09),(cx+.125,cy-.09)]", "coords=[(cx-.125,cy+.12),(cx+.125,cy+.12),(cx-.125,cy-.12),(cx+.125,cy-.12)]")
s=s.replace("curve=.32 if i==0 else -.32", "curve=.45 if i==0 else -.45")
s=s.replace("'Other LTR family'", "'Unrooted trees'")
s=s.replace("fig.savefig(OUT/'Figure_3.svg'", """# Resolve collisions between bootstrap numerals and preserve focal underlines.
fig.canvas.draw()
renderer=fig.canvas.get_renderer()
for axis in fig.axes[:2]:
    occupied=[]
    for text in axis.texts:
        if not text.get_text().isdigit():continue
        original=text.get_position()
        placed=False
        for dx,dy in [(0,0),(-.006,0),(.006,0),(-.012,0),(.012,0),(0,.7),(0,-.7),(-.006,.7),(.006,-.7)]:
            text.set_position((original[0]+dx,original[1]+dy))
            bb=text.get_window_extent(renderer).expanded(1.1,1.05)
            if not any(bb.overlaps(other) for other in occupied):
                occupied.append(bb);placed=True;break
        if not placed:print('SUPPORT_LABEL_OVERLAP',text.get_text(),original)
    for text in axis.texts:
        if text.get_fontweight()!='bold':continue
        bb=text.get_window_extent(renderer)
        x1,y1=axis.transData.inverted().transform((bb.x0,bb.y0-1))
        x2,y2=axis.transData.inverted().transform((bb.x1,bb.y0-1))
        axis.plot([x1,x2],[y1,y2],color=text.get_color(),lw=.5)
fig.savefig(OUT/'Figure_3.svg'""")
s += "\nprint(json.dumps({'output':str(OUT),'trees':manifest['trees']}))\n"
(OUT/'render_figure3.py').write_text(s)
```
