## Supplementary material for "Structural polymorphism and population-variable coding capacity of HERV-K(HML-2) in human pangenomes": render_figure3.py

```
"""Restore all-locus trees and chromosome connections from the verified inputs."""
from pathlib import Path
from copy import deepcopy
from hashlib import sha256
import csv
import json
import os
import sys
os.environ.setdefault('MPLCONFIGDIR', '/private/tmp/hml2_restore_mpl')
sys.dont_write_bytecode = True
import matplotlib
matplotlib.use('Agg')
import matplotlib.pyplot as plt
from matplotlib.patches import FancyArrowPatch, FancyBboxPatch, Rectangle
from matplotlib.path import Path as MPath
from Bio import Phylo
import numpy as np
sys.path.insert(0,'source_worktree/project/manuscript')
import build_narrative_main_figures as owner

PROJECT = Path('source_worktree/project')
INPUT = PROJECT/'results/acroc_resolved_20260914'
PHY = INPUT/'phylogeny'
OUT = Path(__file__).resolve().parent
OUT.mkdir(exist_ok=True)
plt.rcParams.update({'font.family':'Arial', 'font.size':7, 'svg.fonttype':'none', 'pdf.fonttype':42})
owner.PHY = PHY
subfamilies = owner.load_subfamily_authority()
colors={'LTR5_Hs':'#007F73','LTR5A':'#CEAA36','LTR5B':'#D7733F','non-LTR5':'#444444'}
groups=[('Acrocentric Type I',['13p13','15p13b'],'#444444'),
        ('Acrocentric Type II',['15p13a','21p13','22p13'],'#444444'),
        ('1p36.21',['1p36.21a','1p36.21b','1p36.21c'],'#444444'),
        ('8p23.1',['8p23.1b','8p23.1c','8p23.1d','8p23.1e'],'#444444'),
        ('Xq28',['Xq28a','Xq28b'],'#444444')]
edges=list(csv.DictReader((INPUT/'Figure_3C_exact_nucleotide_edges.tsv').open(), delimiter='\t'))
edge_map={tuple(sorted((r['locus_1'],r['locus_2']))):float(r['percent_difference']) for r in csv.DictReader((OUT/'pairwise_differences_network.tsv').open(),delimiter='\t')}
manifest={'inputs':{},'trees':{},'edges':edges}
for path in [PHY/'hml2_pan_ltr_expanded_tree.nwk', PHY/'hml2_pan_orf_pol_tree.nwk', INPUT/'Figure_3C_exact_nucleotide_edges.tsv', PHY/'locus_subfamily.tsv']:
    manifest['inputs'][str(path)]=sha256(path.read_bytes()).hexdigest()

shared_loci = set.intersection(*(
    {tip.name.split('__')[0] for tip in Phylo.read(PHY/name,'newick').get_terminals()}
    for name in ('hml2_pan_ltr_expanded_tree.nwk','hml2_pan_orf_pol_tree.nwk')
))
manifest['comparison_loci'] = sorted(shared_loci)


def select_tree_tips(full):
    """Keep every modal locus tip and common-locus comparison neighbors."""
    keep=owner.representative_tip_names(full)
    for locus in ('19p12c','10q24.2'):
        query=next((t for t in full.get_terminals() if t.name.startswith(locus+'__hap1')),None)
        if query is not None and locus in shared_loci:
            candidates=[t for t in full.get_terminals()
                        if t.name.split('__')[0] in shared_loci - {locus}]
            shortest=min(full.distance(query,t) for t in candidates)
            keep.update(t.name for t in candidates if abs(full.distance(query,t)-shortest)<1e-10)
    return keep


fig=plt.figure(figsize=(6.5,7.25))
def draw_tree(rect, region, filename):
    ax=fig.add_axes(rect)
    full=Phylo.read(OUT/('LTR_bootstrap.nwk' if region=='LTR' else 'pol_bootstrap.nwk'),'newick')
    from analyze import split_key
    support_rows=list(csv.DictReader((OUT/'bootstrap_splits.tsv').open(),delimiter='\t'))
    support={tuple(r['split'].split(';')):float(r['bootstrap_percent']) for r in support_rows if r['region']==('LTR' if region=='LTR' else 'pol')}
    universe={x.name for x in full.get_terminals()}
    full.root_at_midpoint()
    full.rooted=False
    for c in full.get_nonterminals():
        c.confidence=support.get(split_key({x.name for x in c.get_terminals()},universe))
    keep=select_tree_tips(full)
    tree=deepcopy(full)
    for tip in list(tree.get_terminals()):
        if tip.name not in keep: tree.prune(tip)
    tree.ladderize()
    tips=tree.get_terminals()
    assert {x.name.split('__')[0] for x in tips} == {x.name.split('__')[0] for x in full.get_terminals()}
    ypos={tip:i for i,tip in enumerate(tips)}
    def locate(c):
        if c not in ypos: ypos[c]=sum((locate(c.clades[0]),locate(c.clades[-1])))/2
        return ypos[c]
    for c in tree.find_clades(order='postorder'):locate(c)
    depths=tree.depths()
    xmax=max(depths.values())
    for c in tree.find_clades():
        if c.clades:
            ax.plot([depths[c],depths[c]],[ypos[c.clades[0]],ypos[c.clades[-1]]],color='#777777',lw=.38)
        for child in c.clades:
            col=colors[subfamilies[child.name.split('__')[0]]] if child.is_terminal() else '#777777'
            ax.plot([depths[c],depths[child]],[ypos[child],ypos[child]],color=col,lw=.45)
    for c in tree.get_nonterminals():
        if c is not tree.root and c.confidence is not None and c.confidence >=70:
            ax.text(depths[c],ypos[c]+.52,str(round(c.confidence)),fontsize=4.4,ha='right',va='bottom',color='#333333',bbox={'fc':'white','ec':'none','pad':.03},zorder=6)
    # Alternate label columns, retaining every locus rather than selecting a few.
    for i,tip in enumerate(tips):
        locus=tip.name.split('__')[0]
        target=xmax*(1.06+(i%2)*.67)
        ax.plot([depths[tip],target*.99],[ypos[tip],ypos[tip]],color='#CCCCCC',lw=.28)
        display=owner.display_locus_name(locus)
        if sum(t.name.split('__')[0]==locus for t in tips)>1:display+=' c'+tip.name.split('__hap')[1].split('__')[0]
        ax.text(target,ypos[tip],display,fontsize=6.4,color=colors[subfamilies[locus]],va='center',fontweight='bold' if locus in ['19p12c','10q24.2','4q35.2_hg38'] else 'normal')
    ax.set(xlim=(-.01*xmax,2.42*xmax),ylim=(-1,len(tips)))
    ax.set_title(f'{region}   {len(set(t.name.split("__")[0] for t in tips))} loci',fontsize=9,pad=5)
    ax.spines[['top','left','right']].set_visible(False)
    ax.spines['bottom'].set_bounds(0,xmax)
    ax.tick_params(axis='y',left=False,labelleft=False)
    ax.set_xticks([0,round(xmax/2,2),round(xmax,2)])
    ax.tick_params(axis='x',labelsize=6.5,pad=1)
    ax.set_xlabel('Nucleotide differences per aligned base',fontsize=6.5,labelpad=1)
    manifest['trees'][region]={'loci':len(set(t.name.split('__')[0] for t in tips)),'full_tree_tips':len(full.get_terminals()),'selected_tips':[t.name for t in tips], 'selection':'Most frequent exact sequence cluster at every locus, plus nearest other-locus neighbors of modal 19p12c and 10q24.2 clusters restricted to loci represented in both LTR and Pol trees. Distances and pruning use the existing full tree without refitting.'}
    return ax

draw_tree([.035,.379,.455,.565],'LTR','hml2_pan_ltr_expanded_tree.nwk')
draw_tree([.545,.379,.445,.565],'Pol','hml2_pan_orf_pol_tree.nwk')
fig.text(.008,.976,'A',weight='bold',fontsize=12)
fig.text(.08,.325,'LTR5Hs',color=colors['LTR5_Hs'],fontsize=8)
fig.text(.265,.325,'LTR5A',color=colors['LTR5A'],fontsize=8)
fig.text(.425,.325,'LTR5B',color=colors['LTR5B'],fontsize=8)
fig.text(.59,.325,'Unrooted trees',color=colors['non-LTR5'],fontsize=8)

# Compact curved networks retain every validated edge and nucleotide count.
ax=fig.add_axes([.03,.025,.47,.266]);ax.set(xlim=(0,1),ylim=(0,1));ax.axis('off')
fig.text(.008,.306,'B',weight='bold',fontsize=12)
ax.text(.5,1.02,'Mean nucleotide difference (%)',ha='center',fontsize=9)
centers=[(.23,.77),(.74,.76),(.15,.25),(.53,.24),(.89,.23)]
all_edges=[]
for (title,loci,color),(cx,cy) in zip(groups,centers):
    ax.text(cx,cy+.205,title,fontsize=7,ha='center')
    if len(loci)==2: coords=[(cx-.10,cy),(cx+.10,cy)]
    elif len(loci)==3: coords=[(cx-.13,cy+.04),(cx+.13,cy+.04),(cx,cy-.115)]
    else: coords=[(cx-.125,cy+.12),(cx+.125,cy+.12),(cx-.125,cy-.12),(cx+.125,cy-.12)]
    for i,locus in enumerate(loci):
        x,y=coords[i];ax.scatter(x,y,s=15,color=color,zorder=4)
        label=locus if title.startswith('Acro') else locus[-1]
        ax.text(x,y+(.040 if y>=cy else -.04),label,ha='center',va='bottom' if y>=cy else 'top',fontsize=6.6)
    for i in range(len(loci)):
        for j in range(i+1,len(loci)):
            key=tuple(sorted((loci[i],loci[j])))
            if key not in edge_map:continue
            x1,y1=coords[i];x2,y2=coords[j]
            curve=.13 if (j-i)%2 else -.13
            if len(loci)==4 and {i,j} in [{0,3},{1,2}]:curve=.45 if i==0 else -.45
            patch=FancyArrowPatch((x1,y1),(x2,y2),arrowstyle='-',connectionstyle=f'arc3,rad={curve}',lw=1.2,color=color,shrinkA=3,shrinkB=3)
            ax.add_patch(patch)
            # The midpoint of the quadratic Bezier is displaced by half the curvature.
            mx=(x1+x2)/2+curve*(y2-y1)/2;my=(y1+y2)/2-curve*(x2-x1)/2
            ax.text(mx,my,f'{edge_map[key]:.2f}' if edge_map[key]>=.1 else f'{edge_map[key]:.3f}',fontsize=6.1,ha='center',va='center',bbox={'fc':'white','ec':'none','pad':.35},zorder=5)
            all_edges.append(key)
assert set(all_edges)==set(edge_map)

# Restore vertical ideograms and colored chromosome-spanning connections.
ax=fig.add_axes([.55,.025,.44,.266]);ax.set(xlim=(0,1),ylim=(0,1));ax.axis('off')
fig.text(.514,.306,'C',weight='bold',fontsize=12)
ax.text(.5,1.02,'Chromosomal connections',ha='center',fontsize=9)
chroms=['1','4','8','X','13','15','21','22']
xs=dict(zip(chroms,[.045,.175,.305,.435,.565,.695,.825,.955]));top=.75;bottom=.12;height=top-bottom;width=.039
stains={'gneg':'white','gpos25':'#D9DDE0','gpos50':'#AEB5BA','gpos75':'#737C82','gpos100':'#30363A','gvar':'#C8CDD0','stalk':'#E3E6E8','acen':'#B96868'}
for chrom in chroms:
    bands=owner.CYTOBANDS_HG38[chrom];total=bands[-1][0];x=xs[chrom];start=0
    for end,stain in bands:
        ax.add_patch(Rectangle((x-width/2,top-end/total*height),width,(end-start)/total*height,facecolor=stains[stain],edgecolor='none'))
        start=end
    ax.add_patch(FancyBboxPatch((x-width/2,bottom),width,height,boxstyle='round,pad=0,rounding_size=.008',facecolor='none',edgecolor='#555555',lw=.55))
    ax.text(x,.065,chrom,ha='center',fontsize=7)
marks={}
positions={'1p36.21a':('1',14.2e6),'1p36.21b':('1',14.2e6),'1p36.21c':('1',14.2e6),'8p23.1b':('8',9.55e6),'8p23.1c':('8',9.55e6),'8p23.1d':('8',9.55e6),'8p23.1e':('8',9.55e6),'Xq28a':('X',152e6),'Xq28b':('X',152e6),'13p13':('13',2.3e6),'15p13b':('15',2.1e6),'15p13a':('15',2.1e6),'21p13':('21',1.55e6),'22p13':('22',2.15e6)}
for title,loci,color in groups:
    for i,locus in enumerate(loci):
        chrom,coord=positions[locus];y=top-coord/owner.CYTOBANDS_HG38[chrom][-1][0]*height
        if title=='Acrocentric Type I' and chrom=='15':y+=.024
        if not title.startswith('Acro'): y+=(i-(len(loci)-1)/2)*.017
        marks[locus]=(xs[chrom],y)
        ax.plot([xs[chrom]-.03,xs[chrom]+.03],[y,y],color=color,lw=1.4)
    for i,l1 in enumerate(loci):
        for l2 in loci[i+1:]:
            if tuple(sorted((l1,l2))) not in edge_map:continue
            p1,p2=marks[l1],marks[l2]
            if p1[0]==p2[0]:
                p1=(p1[0]+.023,p1[1]);p2=(p2[0]+.023,p2[1]);curve=.75
            else: curve=-.60 if title.endswith('I') and not title.endswith('II') else .35
            ax.add_patch(FancyArrowPatch(p1,p2,connectionstyle=f'arc3,rad={curve}',arrowstyle='<->',mutation_scale=5,lw=.9,color=color,shrinkA=1,shrinkB=1))
for x,label in zip(xs.values(),['1p36.21','4q35.2','8p23.1','Xq28','13p13','15p13','21p13','22p13']):
    ax.text(x,.94 if label in ['1p36.21','Xq28','15p13','22p13'] else .885,label,ha='center',fontsize=6.3,fontweight='bold' if label=='4q35.2' else 'normal')
# The chromosome map also shows the broader Type-II relationship. Exact
# gene-sequence identity is the criterion for panel B, not for family membership.
# Current LTR and Pol trees place 4q35.2 nearest to acrocentric Type-II sequences.
four_y=bottom+.014
ax.plot([xs['4']-.03,xs['4']+.03],[four_y,four_y],color='#444444',lw=1.4)
# Bracket the Type-II group, rather than asserting a particular source chromosome.
group_y=.813
for chrom in ['15','21','22']:
    ax.plot([xs[chrom],xs[chrom]],[top+.018,group_y],color='#444444',lw=.8,ls='--')
ax.plot([xs['15'],.995],[group_y,group_y],color='#444444',lw=.9,ls='--')
ax.plot([xs['4']+.025,xs['4']+.047,xs['4']+.047,.995,.995],
        [four_y,four_y,.014,.014,group_y],color='#444444',lw=1.0,ls='--',clip_on=False)
# Resolve collisions between bootstrap numerals and preserve focal underlines.
fig.canvas.draw()
renderer=fig.canvas.get_renderer()
for axis in fig.axes[:2]:
    occupied=[]
    for text in axis.texts:
        if not text.get_text().isdigit():continue
        original=text.get_position()
        placed=False
        for dx,dy in [(0,0),(-.006,0),(.006,0),(-.012,0),(.012,0),(0,.7),(0,-.7),(-.006,.7),(.006,-.7)]:
            text.set_position((original[0]+dx,original[1]+dy))
            bb=text.get_window_extent(renderer).expanded(1.1,1.05)
            if not any(bb.overlaps(other) for other in occupied):
                occupied.append(bb);placed=True;break
        if not placed:print('SUPPORT_LABEL_OVERLAP',text.get_text(),original)
    for text in axis.texts:
        if text.get_fontweight()!='bold':continue
        bb=text.get_window_extent(renderer)
        x1,y1=axis.transData.inverted().transform((bb.x0,bb.y0-1))
        x2,y2=axis.transData.inverted().transform((bb.x1,bb.y0-1))
        axis.plot([x1,x2],[y1,y2],color=text.get_color(),lw=.5)
fig.savefig(OUT/'Figure_3.svg',facecolor='white')
fig.savefig(OUT/'Figure_3.pdf',facecolor='white')
fig.savefig(OUT/'Figure_3.png',dpi=450,facecolor='white')
plt.close(fig)
manifest['geometry_inches']=[6.5,7.25]
manifest['panel_C']='Schematic cytoband positions, chromosomes scaled separately. Solid double-headed connections encode exact sequence sharing. A dashed connection links 4q35.2 with the acrocentric Type-II group on the basis of the LTR and Pol relationships, without assigning a duplication direction or exact-sequence edge.'
(OUT/'Figure_3_manifest.json').write_text(json.dumps(manifest,indent=2)+'\n')

print(json.dumps({'output':str(OUT),'trees':manifest['trees']}))
```
