## Supplementary material for "Structural polymorphism and population-variable coding capacity of HERV-K(HML-2) in human pangenomes": xq28.py

```
from pathlib import Path
from hashlib import sha256
from collections import defaultdict
import csv,json,subprocess
from Bio import SeqIO
from Bio.Seq import Seq
OUT=Path(__file__).resolve().parent;P=OUT.parents[2]/'project/results/acroc_resolved_20260914/nucleotide_analysis'
rows=[r for r in csv.DictReader((P/'gene_sequence_observations.tsv').open(),delimiter='\t') if r['locus'] in ['Xq28a','Xq28b']]
seqs={r.id:str(r.seq).upper() for r in SeqIO.parse(P/'sequences.fa','fasta') if r.id in {r['query'] for r in rows}}
known={};raw_cache={};obs=[]
for r in rows:
 raw=seqs[r['query']];target=r['sha256'];L=int(r['length']);key=(raw,target)
 if key in raw_cache:start,strand,found=raw_cache[key]
 else:
  # All known exact Env strings are searched before a new hash scan.
  hits=[]
  for strand,s in [('+',raw),('-',str(Seq(raw).reverse_complement()))]:
   if target in known:
    start=s.find(known[target]);
    if start>=0:hits.append((start,strand,known[target]))
   else:
    for start in range(len(s)-L+1):
     frag=s[start:start+L]
     if sha256(frag.encode()).hexdigest()==target:hits.append((start,strand,frag))
  assert len(hits)==1,(r,len(hits));start,strand,found=hits[0];raw_cache[key]=(start,strand,found)
 known[target]=found;obs.append(dict(**r,oriented_source_start=start,source_strand=strand))
with (OUT/'Xq28_env_exact_sequences.fasta').open('w') as f:
 for key,s in known.items():f.write('>'+key+'\n'+s+'\n')
with (OUT/'Xq28_env_source_observations.tsv').open('w') as f:
 w=csv.DictWriter(f,fieldnames=list(obs[0]),delimiter='\t',lineterminator='\n');w.writeheader();w.writerows(obs)
print('Recovered',len(rows),'copies',len(known),'exact Env sequences')
with (OUT/'Xq28_env_aligned.fasta').open('w') as f, (OUT/'Xq28_mafft.log').open('w') as err:
 subprocess.run(['/usr/local/bin/mafft','--auto',str(OUT/'Xq28_env_exact_sequences.fasta')],stdout=f,stderr=err,check=True)
```
