## Supplementary material for "Structural polymorphism and population-variable coding capacity of HERV-K(HML-2) in human pangenomes": all_alleles.py

```
"""Compare every recovered source and retain regional site-frequency evidence."""
from pathlib import Path
from collections import Counter,defaultdict
import csv,json,re,itertools
import numpy as np
from Bio import SeqIO,Phylo
from Bio.Seq import Seq
from Bio.Phylo.TreeConstruction import DistanceTreeConstructor,DistanceMatrix

OUT=Path(__file__).resolve().parent
PROJECT=Path('/Users/Daniel/Documents/HML-2 manuscript work/project')
LOCI=['4q35.2_hg38','4p16.3a','15p13a','21p13','22p13']
def rows(p):
    with p.open() as f:return list(csv.DictReader(f,delimiter='\t'))
def write(name,data):
    with (OUT/name).open('w') as f:
        w=csv.DictWriter(f,list(data[0]),delimiter='\t');w.writeheader();w.writerows(data)
ad=[r for r in rows(PROJECT/'results/acroc_resolved_20260914/phylogeny/alignment_admission.tsv') if r['locus'] in LOCI]
meta={f'p{i:04d}':r for i,r in enumerate(ad)}
src={r.id:str(r.seq).upper() for r in SeqIO.parse(OUT/'genomic_sources.fa','fasta')}
hit=defaultdict(list)
for line in (OUT/'genomic_sources.paf').read_text().splitlines():
    h=line.split('\t');hit[h[0]].append(h)
records=[];arrays=[]
for k in meta:
    if k not in src or len(hit[k])!=1:continue
    h=hit[k][0];seq=src[k] if h[4]=='+' else str(Seq(src[k]).reverse_complement())
    q=int(h[2]) if h[4]=='+' else len(seq)-int(h[3]);t=int(h[7]);p=['-']*7283;mapped=0
    cigar=next(z[5:] for z in h[12:] if z.startswith('cg:Z:'))
    for n,op in re.findall(r'(\d+)([MID=X])',cigar):
        n=int(n)
        if op in 'M=X':p[t:t+n]=seq[q:q+n];t+=n;q+=n;mapped+=n
        elif op=='I':q+=n
        else:t+=n
    if mapped<4000:continue
    records.append({'key':k,**meta[k]});arrays.append(p)
assert len(records)==426
assert len({(r['sample'],r['haplotype'],r['Source_Identifier']) for r in records})==426
write('all_source_membership.tsv',records)
arr=np.array(arrays,dtype='U1');groups={loc:np.array([i for i,r in enumerate(records) if r['locus']==loc]) for loc in LOCI}
counts={loc:np.array([(arr[idx]==b).sum(axis=0) for b in 'ACGT']).T for loc,idx in groups.items()}
coverage={loc:counts[loc].sum(axis=1) for loc in LOCI}
common=np.ones(7283,dtype=bool)
for loc in LOCI:common &= coverage[loc]>=.9*len(groups[loc])
freq={loc:np.divide(counts[loc],coverage[loc][:,None],where=coverage[loc][:,None]>0,out=np.zeros((7283,4))) for loc in LOCI}
regions={'5p_LTR':(6,1000),'internal':(1000,6286),'3p_LTR':(6286,7283),'whole':(0,7283)}
distances=[];nearest=[]
for region,(a,b) in regions.items():
    keep=common.copy();keep[:a]=False;keep[b:]=False;n=int(keep.sum())
    pi={loc:float(np.mean((1-np.sum(freq[loc][keep]**2,axis=1))*coverage[loc][keep]/(coverage[loc][keep]-1))) for loc in LOCI}
    for loc in LOCI:
        distances.append({'region':region,'locus_a':loc,'locus_b':loc,'callable_sites':n,'dxy':pi[loc],'net_dxy':0.})
    for a,b in itertools.combinations(LOCI,2):
        d=float(np.mean(1-np.sum(freq[a][keep]*freq[b][keep],axis=1)))
        distances.append({'region':region,'locus_a':a,'locus_b':b,'callable_sites':n,'dxy':d,'net_dxy':d-(pi[a]+pi[b])/2})
    # Every source's closest source at another locus, with all ties retained.
    # Equal region mask, pairwise ACGT deletion within that mask.
    x=arr[:,keep];called=np.isin(x,list('ACGT'))
    for i,r in enumerate(records):
        joint=called & called[i]
        ns=joint.sum(axis=1);ms=((x!=x[i]) & joint).sum(axis=1)
        ds=np.divide(ms,ns,where=ns>0,out=np.full(len(records),np.inf))
        for j,z in enumerate(records):
            if z['locus']==r['locus']:ds[j]=np.inf
        best=float(ds.min());best_loci=sorted({records[j]['locus'] for j in np.flatnonzero(np.isclose(ds,best,rtol=0,atol=1e-12))})
        nearest.append({'region':region,'key':r['key'],'locus':r['locus'],'sample':r['sample'],'best_distance':best,'nearest_loci':'|'.join(best_loci)})
write('all_allele_regional_distances.tsv',distances);write('all_allele_nearest_sources.tsv',nearest)
aggregated=[]
for region in regions:
    for loc in LOCI:
        c=Counter(r['nearest_loci'] for r in nearest if r['region']==region and r['locus']==loc)
        for target,n in c.most_common():aggregated.append({'region':region,'locus':loc,'nearest_loci':target,'source_copies':n})
write('all_allele_nearest_summary.tsv',aggregated)
site_rows=[]
for pos in np.flatnonzero(common):
    modal={loc:'ACGT'[int(np.argmax(freq[loc][pos]))] for loc in LOCI}
    if len(set(modal.values()))<2:continue
    rr={'reference_position_1based':int(pos+1)}
    for loc in LOCI:
        rr[loc+'_base']=modal[loc];rr[loc+'_frequency']=float(max(freq[loc][pos]));rr[loc+'_called']=int(coverage[loc][pos])
    site_rows.append(rr)
write('locus_distinguishing_sites.tsv',site_rows)
fixed=[]
for a,b in itertools.combinations(LOCI,2):
    am=np.argmax(freq[a],axis=1);bm=np.argmax(freq[b],axis=1)
    is_fixed=common & (np.max(freq[a],axis=1)==1) & (np.max(freq[b],axis=1)==1) & (am!=bm)
    fixed.append({'locus_a':a,'locus_b':b,'fixed_differences':int(is_fixed.sum()),'positions_1based':','.join(str(x+1) for x in np.flatnonzero(is_fixed))})
write('fixed_locus_differences.tsv',fixed)
complete=np.all(np.isin(arr,list('ACGT')),axis=0)
exact=defaultdict(Counter)
for i,r in enumerate(records):exact[''.join(arr[i,complete])][r['locus']]+=1
shared=[]
for seq,c in exact.items():
    if len(c)>1:
        shared.append({'profile':f'shared_{len(shared)+1}','complete_columns':int(complete.sum()),**{loc:c.get(loc,0) for loc in LOCI}})
write('shared_complete_site_profiles.tsv',shared)
summary={'source_copies':len(records),'donors':len({r['sample'] for r in records}),'shared_90pct_coverage_sites':int(common.sum()),
  'all_source_complete_columns':int(complete.sum()),'cross_locus_exact_complete_site_profiles':len(shared),
  'regions':{region:int(common[a:b].sum()) for region,(a,b) in regions.items()},'locus_distinguishing_sites':len(site_rows),
  'method':'All 426 source sequences, equal locus weighting for region comparison, per-site nucleotide frequencies. Dxy is mean between-locus mismatch probability. Net Dxy subtracts half the sum of unbiased within-locus diversity estimates. Sites require 90% source callability in every locus. Negative net values are retained.'}
(OUT/'all_allele_summary.json').write_text(json.dumps(summary,indent=2))
print(json.dumps(summary,indent=2))
print('DISTANCES',json.dumps([r for r in distances if r['region']=='whole']))
print('NEAREST',json.dumps([r for r in aggregated if r['region']=='whole']))
```
