## Supplementary material for "Structural polymorphism and population-variable coding capacity of HERV-K(HML-2) in human pangenomes": flank_history.py

```
"""Reference flank genealogy using uniquely body-spanning primary alignments."""
from pathlib import Path
import csv,json,re,itertools,subprocess
import numpy as np
from Bio import SeqIO,Phylo
from Bio.Seq import Seq
from Bio.Phylo.TreeConstruction import DistanceTreeConstructor,DistanceMatrix

OUT=Path(__file__).resolve().parent
PROJECT=Path('/Users/Daniel/Documents/HML-2 manuscript work/project')
REF=PROJECT/'results/type2_insertion_ancestry_audit_20260802/sequences'
LOCI=['4q35.2_hg38','15p13a','21p13','22p13']
def write(name,rows):
    if rows:
        with (OUT/name).open('w') as f:
            w=csv.DictWriter(f,list(rows[0]),delimiter='\t');w.writeheader();w.writerows(rows)
seqs={};bounds={}
for loc,n in zip(LOCI,[7287,7283,7275,7279]):
    seq=str(next(SeqIO.parse(REF/f'{loc}.fa','fasta')).seq).upper()
    end=len(seq)-100000;start=end-n
    if loc.startswith('4q'):
        seq=str(Seq(seq).reverse_complement());start,end=len(seq)-end,len(seq)-start
    # Physical 8 kb flanks on both sides, before any alignment.
    seqs[loc]=seq[start-8000:end+8000];bounds[loc]=[8000,8000+n]
q=OUT/'reference_flank_windows.fa'
q.write_text(''.join(f'>{k}\n{s}\n' for k,s in seqs.items()))
t=OUT/'reference_flank_target.fa';t.write_text('>15p13a\n'+seqs['15p13a']+'\n')
p=subprocess.run(['/Users/Daniel/opt/anaconda3/bin/minimap2','-x','asm20','-c','--eqx','--secondary=no',str(t),str(q)],capture_output=True,text=True,check=True)
(OUT/'reference_flank_windows.paf').write_text(p.stdout)
hits={k:[] for k in LOCI}
for line in p.stdout.splitlines():
    h=line.split('\t')
    if int(h[7])<=3000 and int(h[8])>=20283:hits[h[0]].append(h)
projections={};evidence=[];indels=[]
for loc,s in seqs.items():
    assert len(hits[loc])==1,(loc,len(hits[loc]))
    h=hits[loc][0];assert h[4]=='+' and int(h[11])>=50
    cigar=next(x[5:] for x in h[12:] if x.startswith('cg:Z:'))
    qi,ti=int(h[2]),int(h[7]);projection=['-']*len(seqs['15p13a'])
    for size,op in re.findall(r'(\d+)([MID=X])',cigar):
        n=int(size)
        if op in 'M=X':projection[ti:ti+n]=s[qi:qi+n];qi+=n;ti+=n
        elif op=='I':
            indels.append({'locus':loc,'ref_position':ti,'operation':op,'length':n,'sequence':s[qi:qi+n]});qi+=n
        else:
            indels.append({'locus':loc,'ref_position':ti,'operation':op,'length':n,'sequence':seqs['15p13a'][ti:ti+n]});ti+=n
    projections[loc]=''.join(projection)
    evidence.append({'locus':loc,'query_length':len(s),'query_start':h[2],'query_end':h[3],'reference_start':h[7],'reference_end':h[8],'mapq':h[11]})
write('reference_flank_admission.tsv',evidence);write('reference_flank_indels.tsv',indels)
(OUT/'reference_flanks.aln.fa').write_text(''.join(f'>{k}\n{s}\n' for k,s in projections.items()))
arr=np.array([list(projections[k]) for k in LOCI]);common=np.all(np.isin(arr,list('ACGT')),axis=0)
regions={'upstream_5kb':(3000,8000),'element':(8000,15283),'downstream_5kb':(15283,20283),'both_flanks_5kb':None}
rng=np.random.default_rng(20260918);distances=[];patterns=[];boot=[]
for region,span in regions.items():
    keep=common.copy()
    if span:
        a,b=span;keep[:a]=False;keep[b:]=False
    else:
        keep[:3000]=False;keep[8000:15283]=False;keep[20283:]=False
    positions=np.flatnonzero(keep);x=arr[:,keep];n=len(positions)
    pair_sites={}
    for i,j in itertools.combinations(range(4),2):
        ds=(x[i]!=x[j]);pair_sites[i,j]=ds
        distances.append({'region':region,'locus_a':LOCI[i],'locus_b':LOCI[j],'mismatches':int(ds.sum()),'callable_sites':n,'p_distance':float(ds.mean())})
    # Four-point split scores. Exact sitewise patterns retained to explain support.
    split_names=['4q+15 | 21+22','4q+21 | 15+22','4q+22 | 15+21']
    score=np.array([pair_sites[0,1].astype(int)+pair_sites[2,3],pair_sites[0,2].astype(int)+pair_sites[1,3],pair_sites[0,3].astype(int)+pair_sites[1,2]])
    c=np.zeros(4,int)
    for _ in range(2000):
        ss=score[:,rng.integers(n,size=n)].sum(axis=1);w=np.flatnonzero(ss==ss.min());c[w[0] if len(w)==1 else 3]+=1
    for k,label in enumerate(split_names+['tie']):boot.append({'region':region,'split':label,'bootstrap_count':int(c[k]),'replicates':2000})
    for pi,pos in enumerate(positions):
        bases=x[:,pi]
        if len(set(bases))==1:continue
        row={'region':region,'reference_position_1based':int(pos+1),'element_relative_position':int(pos-8000+1)}
        row.update(dict(zip(LOCI,bases.tolist())));patterns.append(row)
write('reference_flank_distances.tsv',distances);write('reference_flank_site_patterns.tsv',patterns);write('reference_flank_split_bootstrap.tsv',boot)
print(json.dumps({'distances':distances,'bootstrap':boot},indent=2))
```
