## Supplementary material for "Structural polymorphism and population-variable coding capacity of HERV-K(HML-2) in human pangenomes": genomic_recovery.py

```
"""Restore genomic LTR boundaries before comparing the telomeric Type-II family."""
from pathlib import Path
from collections import Counter, defaultdict
from hashlib import sha256
import csv, json, subprocess, re, itertools
import numpy as np
from Bio import SeqIO, Phylo
from Bio.Seq import Seq
from Bio.Align import PairwiseAligner
from Bio.Phylo.TreeConstruction import DistanceTreeConstructor, DistanceMatrix

OUT=Path(__file__).resolve().parent
PROJECT=Path('/Users/Daniel/Documents/HML-2 manuscript work/project')
PHY=PROJECT/'results/acroc_resolved_20260914/phylogeny'
CAT=PROJECT/'results/rec_exon_boundary_correction_20260915/combined_hml2_orf_analysis.RESOLVED.REC_CORRECTED.tsv'
REF=PROJECT/'results/type2_insertion_ancestry_audit_20260802/sequences'
LOCI=['4q35.2_hg38','4p16.3a','15p13a','21p13','22p13']
def rows(p):
    with p.open() as f:return list(csv.DictReader(f,delimiter='\t'))
def write(name,data):
    if not data:return
    with (OUT/name).open('w') as f:
        w=csv.DictWriter(f,fieldnames=list(data[0]),delimiter='\t');w.writeheader();w.writerows(data)
def fasta(p):return {r.id:str(r.seq).upper() for r in SeqIO.parse(p,'fasta')}
def dist(a,b):
    pairs=[(x,y) for x,y in zip(a,b) if x in 'ACGT' and y in 'ACGT']
    return sum(x!=y for x,y in pairs),len(pairs)

refseqs={}
for loc,n in [('4q35.2_hg38',7287),('15p13a',7283),('21p13',7275),('22p13',7279)]:
    s=next(iter(fasta(REF/f'{loc}.fa').values()));end=len(s)-100000;start=end-n
    x=s[start:end]
    if loc=='4q35.2_hg38':x=str(Seq(x).reverse_complement())
    refseqs[loc]=x
target=refseqs['15p13a']
aligner=PairwiseAligner(mode='local',match_score=2,mismatch_score=-3,open_gap_score=-10,extend_gap_score=-0.5)
pair=aligner.align(target[:1200],target[-1200:])[0]
a0,a1=int(pair.coordinates[0,0]),int(pair.coordinates[0,-1])
b0,b1=[int(x)+len(target)-1200 for x in [pair.coordinates[1,0],pair.coordinates[1,-1]]]
assert a1-a0>850 and b1-b0>850,(a0,a1,b0,b1)
intervals={'5p':(a0,a1),'3p':(b0,b1)}
(OUT/'reference_LTR_boundary_alignment.txt').write_text(str(pair))
refpath=OUT/'genomic_reference.fa';refpath.write_text('>15p13a\n'+target+'\n')
admissions=[r for r in rows(PHY/'alignment_admission.tsv') if r['locus'] in LOCI]
catalog={r['ID_Full']:r for r in rows(CAT) if r['Locus'].removeprefix('HML-2_') in LOCI}
sources={};metadata={};excluded=[]
for i,r in enumerate(admissions):
    cr=catalog[r['ID_Full']]
    paths=[Path(cr['source_record_path_v3']),Path('/Users/Daniel/Documents/HML2_project_data/processed_loci')/cr['orig_Locus']/f"{r['ID_Full']}.fa"]
    candidates=[p for p in paths if p.is_file() and p.suffix.lower() in {'.fa','.fasta','.fna'}]
    if not candidates:
        excluded.append({'ID_Full':r['ID_Full'],'reason':'raw_source_not_local'});continue
    p=candidates[0];rs=fasta(p);assert len(rs)==1
    key=f'p{i:04d}';sources[key]=next(iter(rs.values()))
    metadata[key]={**r,'source_path':str(p),'source_sha256':sha256(p.read_bytes()).hexdigest(),'is_reference':0}
for loc,s in refseqs.items():
    key='ref_'+loc;sources[key]=s;metadata[key]={'locus':loc,'sample':'GRCh38' if loc.startswith('4q') else 'CHM13','ID_Full':key,'is_reference':1}
qpath=OUT/'genomic_sources.fa'
with qpath.open('w') as f:
    for k,s in sources.items():f.write(f'>{k}\n{s}\n')
r=subprocess.run(['/Users/Daniel/opt/anaconda3/bin/minimap2','-x','asm20','-c','--eqx','--secondary=no',str(refpath),str(qpath)],capture_output=True,text=True,check=True)
(OUT/'genomic_sources.paf').write_text(r.stdout)
hits=defaultdict(list)
for line in r.stdout.splitlines():
    h=line.split('\t');hits[h[0]].append(h)
extracted={};maps={};evidence=[];body_clusters=defaultdict(Counter)
for k,seq in sources.items():
    if len(hits[k])!=1:
        excluded.append({'ID_Full':metadata[k]['ID_Full'],'reason':f'{len(hits[k])}_primary_alignments'});continue
    h=hits[k][0];s=seq if h[4]=='+' else str(Seq(seq).reverse_complement())
    q=int(h[2]) if h[4]=='+' else len(seq)-int(h[3]);t=int(h[7]);mapping={}
    cigar=next(z[5:] for z in h[12:] if z.startswith('cg:Z:'))
    for n,op in re.findall(r'(\d+)([MID=X])',cigar):
        n=int(n)
        if op in 'M=X':
            for j in range(n):mapping[t+j]=q+j
            t+=n;q+=n
        elif op=='I':q+=n
        else:t+=n
    assert t==int(h[8]);maps[k]=mapping
    projection=''.join(s[mapping[i]] if i in mapping else '-' for i in range(len(target)))
    if len(mapping)>=4000 and not metadata[k]['is_reference']:
        body_clusters[metadata[k]['locus']][projection]+=1
    pairseq={}
    for end,(a,b) in intervals.items():
        idx=[mapping[t] for t in range(a,b) if t in mapping]
        if len(idx)<500:continue
        pairseq[end]=s[min(idx):max(idx)+1]
    if len(pairseq)!=2:
        excluded.append({'ID_Full':metadata[k]['ID_Full'],'reason':'fewer_than_500_mapped_bases_in_one_LTR'});continue
    extracted[k]=pairseq
    evidence.append({'key':k,'locus':metadata[k]['locus'],'sample':metadata[k]['sample'],'ID_Full':metadata[k]['ID_Full'],
       'is_reference':metadata[k]['is_reference'],'source_path':metadata[k].get('source_path',str(REF/(metadata[k]['locus']+'.fa'))),
       'source_sha256':metadata[k].get('source_sha256',''),'strand':h[4],'reference_start':h[7],'reference_end':h[8],
       'mapped_reference_bases':len(mapping),'ltr5_bases':len(pairseq['5p']),'ltr3_bases':len(pairseq['3p'])})
unique={}
for pairseq in extracted.values():
    for s in pairseq.values():unique.setdefault(s,f'g{len(unique):04d}')
up=OUT/'genomic_LTRs.unique.fa'
with up.open('w') as f:
    for s,k in unique.items():f.write(f'>{k}\n{s}\n')
rp=OUT/'genomic_LTRs.aln.fa'
r=subprocess.run(['/usr/local/bin/mafft','--quiet','--auto',str(up)],capture_output=True,text=True,check=True);rp.write_text(r.stdout)
aligned=fasta(rp);obs=[];clusters=defaultdict(Counter)
for ev in evidence:
    k=ev['key'];a,b=[aligned[unique[extracted[k][end]]] for end in ['5p','3p']]
    m,n=dist(a,b)
    if n<500:
        excluded.append({'ID_Full':ev['ID_Full'],'reason':'fewer_than_500_jointly_called_LTR_sites'});continue
    ob={**ev,'mismatches':m,'jointly_called_bases':n,'p_distance':m/n,'clock_low_My':m/n/.0045,'clock_high_My':m/n/.0024}
    obs.append(ob)
    if not ev['is_reference']:
        clusters[ev['locus'],'5p'][a]+=1;clusters[ev['locus'],'3p'][b]+=1
write('genomic_paired_LTR_observations.tsv',obs);write('genomic_exclusions.tsv',excluded)
summary=[]
for loc in LOCI:
    rr=[r for r in obs if r['locus']==loc and not r['is_reference']]
    if not rr:continue
    p=float(np.median([r['p_distance'] for r in rr]))
    summary.append({'locus':loc,'source_copies':len(rr),'donors':len({r['sample'] for r in rr}),
        'median_mismatches':float(np.median([r['mismatches'] for r in rr])),
        'median_callable':float(np.median([r['jointly_called_bases'] for r in rr])),
        'median_p':p,'min_p':min(r['p_distance'] for r in rr),'max_p':max(r['p_distance'] for r in rr),
        'clock_low_My':p/.0045,'clock_high_My':p/.0024})
write('genomic_paired_LTR_summary.tsv',summary)
body_modal={loc:c.most_common(1)[0][0] for loc,c in body_clusters.items()}
body_counts=[{'locus':loc,'source_copies':sum(c.values()),'unique_projected_sequences':len(c),'modal_count':c.most_common(1)[0][1]} for loc,c in body_clusters.items()]
write('genomic_body_counts.tsv',body_counts)
with (OUT/'genomic_body_modal.aln.fa').open('w') as f:
    for k,s in body_modal.items():f.write(f'>{k}\n{s}\n')
body_pairs=[]
for (a,sa),(b,sb) in itertools.combinations(body_modal.items(),2):
    m,n=dist(sa,sb);body_pairs.append({'query':a,'target':b,'mismatches':m,'jointly_called_bases':n,'p_distance':m/n,'clock_low_My':m/n/.0045,'clock_high_My':m/n/.0024})
write('genomic_body_pairwise.tsv',body_pairs)
ref_ltrs={}
for k,ends in extracted.items():
    if metadata[k]['is_reference']:
        for end,s in ends.items():ref_ltrs[metadata[k]['locus']+'__'+end]=aligned[unique[s]]
with (OUT/'reference_LTRs.aln.fa').open('w') as f:
    for k,s in ref_ltrs.items():f.write(f'>{k}\n{s}\n')
ref_pairs=[]
for (a,sa),(b,sb) in itertools.combinations(ref_ltrs.items(),2):
    m,n=dist(sa,sb);ref_pairs.append({'query':a,'target':b,'mismatches':m,'jointly_called_bases':n,'p_distance':m/n,'clock_low_My':m/n/.0045,'clock_high_My':m/n/.0024})
write('reference_LTR_pairwise.tsv',ref_pairs)
modal={f'{loc}__{end}':v.most_common(1)[0][0] for (loc,end),v in clusters.items()}
comparisons=[]
for (a,sa),(b,sb) in itertools.combinations(modal.items(),2):
    m,n=dist(sa,sb);comparisons.append({'query':a,'target':b,'mismatches':m,'jointly_called_bases':n,'p_distance':m/n,'clock_low_My':m/n/.0045,'clock_high_My':m/n/.0024})
write('genomic_modal_LTR_pairwise.tsv',comparisons)
with (OUT/'genomic_modal_LTRs.aln.fa').open('w') as f:
    for k,s in modal.items():f.write(f'>{k}\n{s}\n')
names=list(modal);mat=[]
for i,a in enumerate(names):
    line=[]
    for b in names[:i]:
        m,n=dist(modal[a],modal[b]);line.append(m/n)
    mat.append(line+[0.])
tree=DistanceTreeConstructor().nj(DistanceMatrix(names,mat));Phylo.write(tree,OUT/'genomic_modal_LTR_tree.nwk','newick')
manifest={'target_intervals_zero_based_half_open':intervals,'target_length':len(target),'input_admissions_sha256':sha256((PHY/'alignment_admission.tsv').read_bytes()).hexdigest(),
    'catalog_sha256':sha256(CAT.read_bytes()).hexdigest(),'paired_source_copies':len(obs),'unique_LTR_sequences':len(unique),
    'pairwise_clock_denominator_per_My':[.0024,.0045],'clock_reference':'Subramanian et al 2011, Methods, doi:10.1186/1742-4690-8-90',
    'clock_scope':'Raw nucleotide substitution distances only. Rate range is sensitivity, not a confidence interval. No source-event direction inferred from an unrooted tree.'}
(OUT/'genomic_manifest.json').write_text(json.dumps(manifest,indent=2))
print(json.dumps({'summary':summary,'references':[r for r in obs if r['is_reference']],'manifest':manifest},indent=2))
```
