## Supplementary figures and images for "Structural polymorphism and population-variable coding capacity of HERV-K(HML-2) in human pangenomes"

### Figure_3.pdf

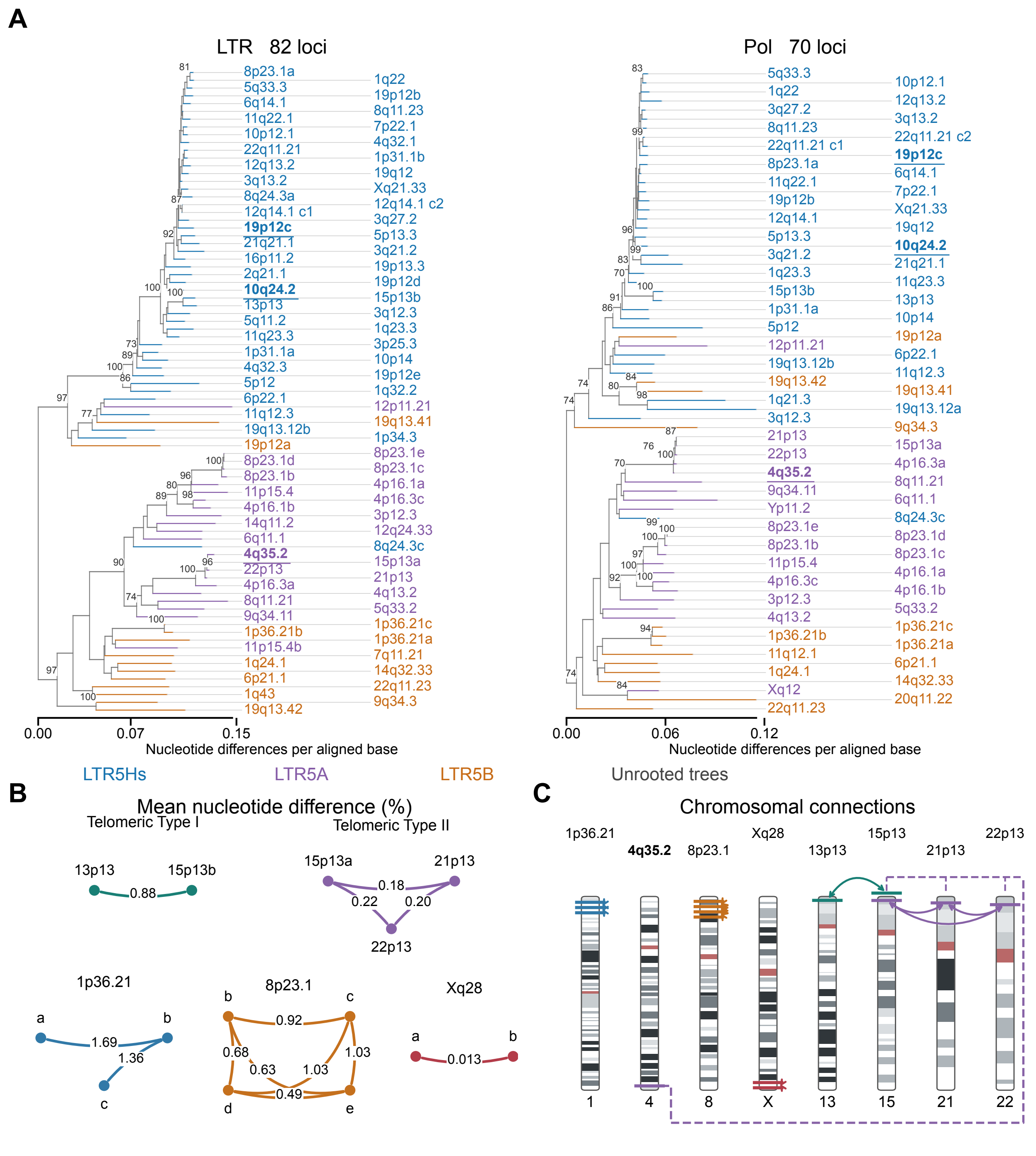

### Figure_3.png

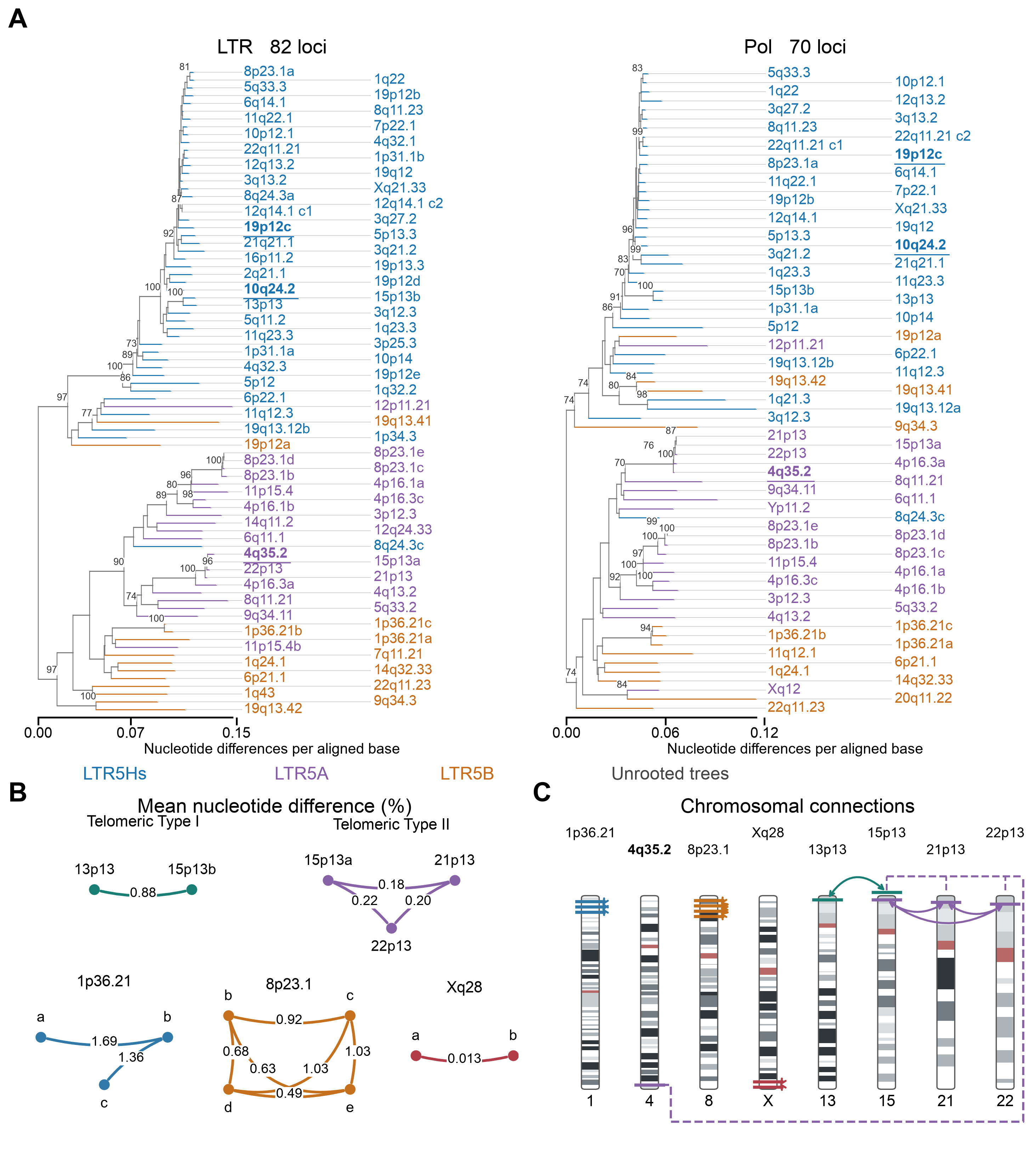

### Telomeric_phylogeny.pdf

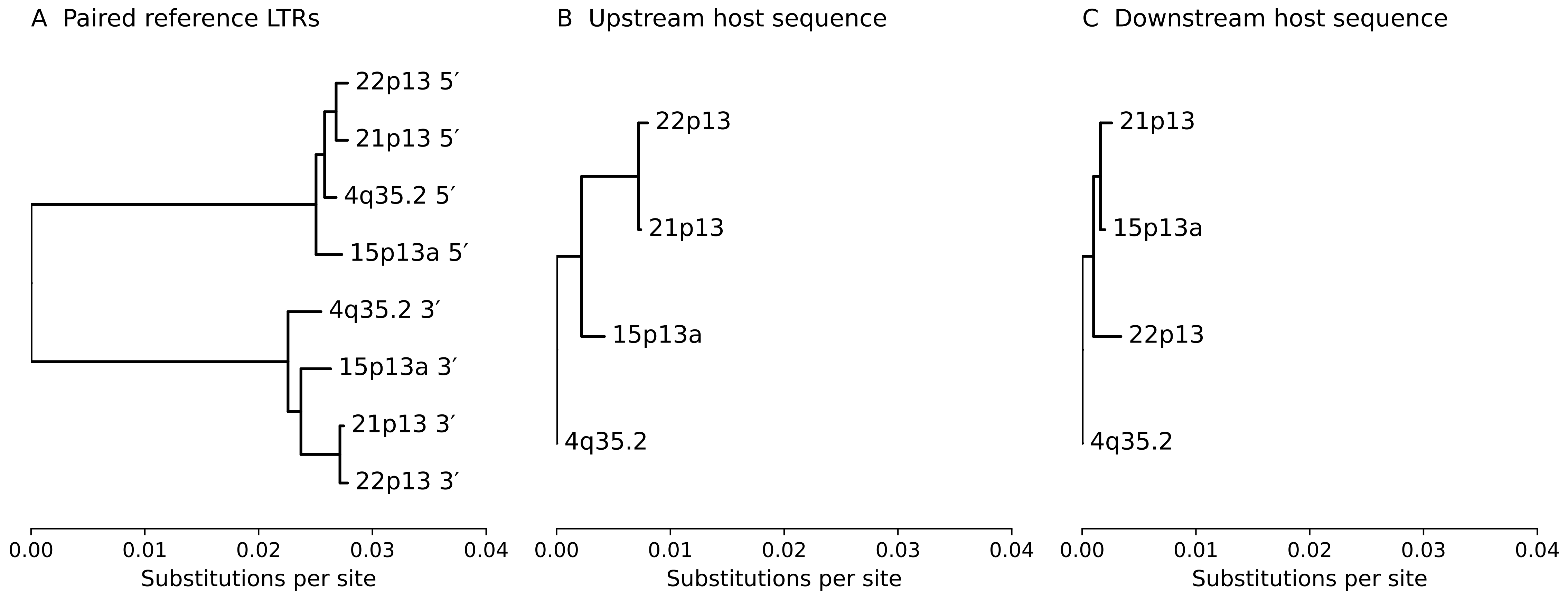

### Telomeric_phylogeny.png

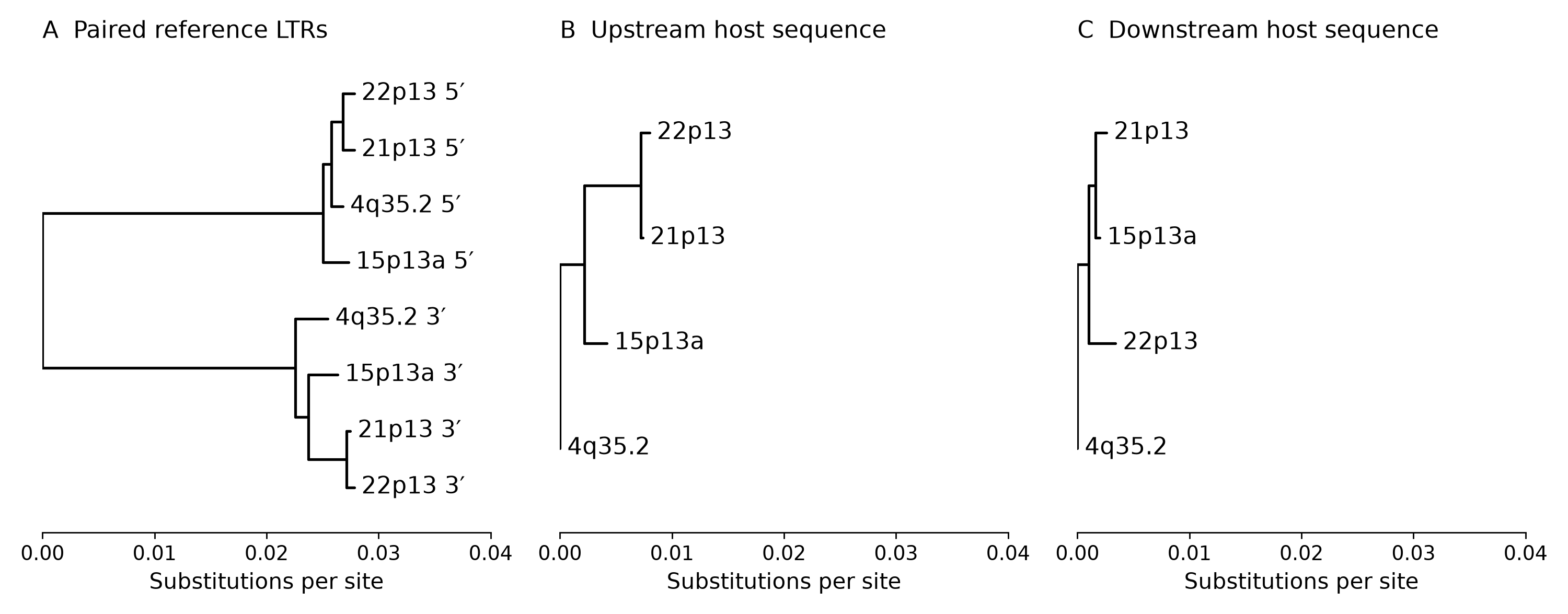
